# An integrative model of AMPA receptor trafficking reveals the central contribution of local translation in subtype-specific kinetics

**DOI:** 10.1101/2025.02.08.637220

**Authors:** Surbhit Wagle, Wouter J. Droogers, Attila Ignacz, Maximilian K. Kracht, Cornelius Bergmann, Anne Bührke, Daniel Choquet, Amparo Acker-Palmer, Nataliya Kraynyukova, Anne-Sophie Hafner, Erin M. Schuman, Tatjana Tchumatchenko

**Affiliations:** Universität Bonn, Universitätsklinikum Bonn, Institute for Experimental Epileptology and Cognition Research, Bonn, Germany; Present address: Bioengineering Department, Imperial College, London, United Kingdom; Donders Institute for Brain, Cognition and Behavior, Radboud University, 6500 GL Nijmegen, Netherlands; Interdisciplinary Institute for Neuroscience (IINS), Université de Bordeaux, UMR 5297 CNRS, 33076 Bordeaux, France; Buchmann Institute of Molecular Life Sciences (BMLS), Goethe Universität, Frankfurt Am Main, Germany; Janelia Research Campus, Howard Hughes Medical Institute, Ashburn, Virginia 20147, USA; Max Planck Institute for Brain Research, Synaptic Plasticity, Frankfurt Am Main, Germany; Bordeaux Imaging Center (BIC), Université de Bordeaux, UMS 3420 CNRS, US 4 INSERM, 33076 Bordeaux, France; Institute of Cell Biology and Neuroscience, Goethe Universität, Frankfurt Am Main, Germany

## Abstract

AMPA-type glutamate receptors (AMPARs) underlie most of the excitatory synaptic transmission in the brain and are crucial for implementing long-term synaptic plasticity. AMPARs form multi-protein complexes containing pore-forming subunits GluA1-4 and auxiliary subunits that regulate AMPAR function and trafficking. Here we show that protein synthesis and trafficking mechanisms determine distributions of AMPAR subtypes across neuronal compartments and their distinct responses to long-term potentiation. While GluA1 and GluA2 are mostly synthesized in the cell body, the auxiliary protein CNIH-2 is locally synthesized in dendrites, and its synthesis increases after the induction of long-term potentiation. Strikingly, CNIH-2 translation is required for the plasma membrane insertion of GluA2-containing receptors but not GluA1-homomeric AMPARs. Our model explains how CNIH-2 local translation allows for the selective trafficking of calcium-impermeable GluA2-containing AMPARs, making their response to plasticity cues slower and longer-lasting than that of calciumpermeable GluA1-homomeric AMPARs.

## Introduction

*α*-amino-3-hydroxy-5-methyl-4-isoxazolepropionic acid-type glutamate receptors (AMPARs) mediate most of the excitatory synaptic communication in the brain and play a central role in the expression of long-term potentiation (LTP). Controlling the number and activity of AMPARs in the postsynaptic density (PSD) effectively regulates the strength of excitatory synaptic transmission^1^. Many neurons in the brain, including pyramidal neurons in the cortex and hippocampus, have dendritic arbors that extend several hundreds of micrometers, allowing them to receive inputs from thousands of other neurons. These dendrites typically contain dendritic spines at a density of 1-2 spines per micron^2^. To detect excitatory inputs (i.e., glutamate release from the axons of presynaptic neurons), the postsynaptic neurons must ensure that their dendritic spines are populated with AMPARs. Which mechanisms do neurons employ to deliver AMPARs to distal sites and maintain their availability throughout the dendritic arbors? Previous experimental studies have shown that having abundant dendritic mRNA is a powerful strategy neurons employ to maintain a dendritic protein pool^3, 4^. Later theoretical work on plasticity-related protein Calcium-Calmodulin Kinase 2 alpha (CamK2a) showed that local dendritic protein synthesis significantly contributes to the homogenous distribution and high concentration of the protein throughout the dendritic arbors^5^. For the reasons outlined below, however, we still lack an understanding of the spatial and temporal mechanisms affecting AMPAR availability at synapses. AMPARs are macromolecular complexes comprising a diverse assembly of tetrameric core of GluA1-4 pore forming subunits^6–8^ as well as auxiliary subunits that are divided in three main families: the transmembrane AMPAR regulatory proteins (TARPs,*γ*-2,*γ*-3,*γ*-4,*γ*-5,*γ*-7, and *γ*-8)^9, 10^, the cornichon homologs (CNIH-2 and CNIH-3)^11^, GSG1L^12^ and the CKAMPs (CKAMP44 and CKAMP44-like proteins)^13, 14^. There are a variety of long-range trafficking possibilities for AMPARs, such as cytoplasmic or membrane diffusion and active transport, as well as local dynamics, including endo- and exocytosis and trafficking between the PSD and an adjacent part of the dendrite. These processes have different time scales and can be affected by the receptor subunit composition and the history of the preceding events at different dendritic locations. Additionally, the specific subunit composition can influence the functional properties of AMPARs, such as their conductance, gating kinetics, and localization density. How do long-range (i.e., a full dendrite) and local (i.e., a spine attached to a piece of dendrite) dynamics interfere or interact with one another? How does the specific subunit composition of AMPARs affect their functional and trafficking properties?

A major divergence in functional properties of AMPAR subtypes arises from the presence of the GluA2 subunit. GluA2-containing AMPARs are impermeable to Ca^2+15^ and are constitutively recruited during plasticity^16^. Conversely, GluA2-lacking AMPARs are Ca^2+^ permeable and can exhibit activity-regulated synaptic recruitment, thereby exhibiting differential recruitment rules at baseline and during synaptic plasticity^1, 17^. Recent work^18^ has suggested that individual subunits of AMPARs have higher mobility when not assembled into functional homo/heterotetramers. However, since a majority of AM-PARs in the neocortex and hippocampus are heterotetramers of Glua1 and GluA2 subunits^1^ this study will concentrate on these two types of AMPARs. For clarity, from now on, we refer to GluA1 homomers as GluA1-homomeric AMPARs and GluA1/2 and GluA2/3 heterotetramers as GluA2-containing AMPARs. Several studies have also suggested that synaptic plasticity demands are rapidly met by fast exocytosis of calcium-permeable GluA1 homotetramers^17^, which are later replaced by calcium-impermeable, GluA2-containing AMPARs. Interestingly, recent studies show that the incorporation of calcium-permeable AMPARs is crucial for the full expression of LTP^19^. In contrast, following chemically induced long-term potentiation (LTP), the exocytosis rate of GluA2-containing AMPARs at stimulated spines and the nearby dendrite increases, resulting in a higher surface concentration of those receptors. Specifically, following LTP induction, the concentration of GluA2-containing receptors slowly increases within the next 1-2 hours, saturates, and remains elevated for several hours^1^. Additionally, GluA1-homomeric AMPARs typically exhibit higher single-channel conductance compared to GluA2-containing receptors^20^. This difference in conductance can influence the strength and amplitude of synaptic responses mediated by these receptors. Also, because the GluA1 subunit possesses the majority of the phosphorylation sites, GluA2-containing receptors may be less dynamically regulated by phosphorylation than GluA1-homomeric receptors^21^. At the cell surface, GluA1-homomeric AMPARs show a tighter synaptic localization compared to GluA2-containing ones that appear more diffused and constitute a reserve pool of extrasynaptic AMPARs^22^.

In most neurons, different types of AMPAR auxiliary subunits co-exist. Such heterogeneity impacts the functional properties of AMPARs, such as their trafficking kinetics, stability in the plasma membrane and the PSD, and channel conductance^23^. All families of auxiliary proteins promote AMPAR surface expression, but each family modulates gating and pore properties differently. For example, CNIH-2 has been shown to slow receptor deactivation significantly - by up to fivefold - thereby extending the excitatory postsynaptic current duration, regardless of the GluA subunit combination^24^. As a result, synapses expressing CNIH-2-containing AMPARs exhibit a larger net depolarizing ion influx into the postsynaptic neuron. However, the precise role of CNIH-2 is still debated, with some seemingly conflicting findings. Initially, CNIH-2 was believed to assist AMPAR transport between the ER and Golgi apparatus (GA) in hippocampal CA1 neurons^25^. However, some studies suggest that CNIH-2 can bypass this early anterograde pathway and directly transport AMPARs to the plasma membrane^26^. Auxiliary subunit interactions can be dependent on the pore-forming subunit composition of AMPARs. For instance, CNIH-2 shows a stronger preference for GluA1/2 heteromers over GluA2/3 receptors^27^. Nonetheless, there are conflicting findings on whether CNIH-2 binds preferentially to GluA2-lacking or GluA2-containing AMPARs^28, 29^. Hence, it is still unclear how auxiliary subunits affect AMPAR availability at synapses and response to LTP.

Here, we have developed a mathematical model to understand how diverse transport mechanisms and their biologically plausible temporal changes impact the availability of receptors at different dendritic locations. At the start, we used experimental data from live-cell HaloTag labeling of surface and intracellular GluA2 endogenously tagged with HaloTag via CRISPR/Cas9-mediated gene editing to establish a well-characterized baseline model for long-range and local AMPAR trafficking. Using a series of assays that preserve AMPAR native stoichiometry and our model, we analyze which trafficking mechanisms are most critical for the availability of AMPARs at distal dendritic sites and how their exocytosis-to-endocytosis ratio is regulated across different dendritic locations. Finally, we examine the mechanisms that can explain a rapid decrease of GluA1-homomeric receptors and a sustained increase in GluA2-containing AMPAR concentration following cLTP induction. Our model suggests that a prolonged increase in GluA2-containing AMPAR exocytosis can account for the differences in the concentration dynamics of GluA1-homomers and GluA2-containing AMPARs. Interestingly, we show that the auxiliary subunit CNIH-2 is abundantly synthesized in dendrites, and chemical-LTP induction increases its synthesis. Moreover, interfering with CNIH-2 synthesis strikingly reduces the surface abundance of GluA2-containing AMPAR but does not significantly alter surface GluA1-homomers. This observation suggests that CNIH-2—an auxiliary subunit that specifically promotes GluA2 (but not GluA1) exocytosis—may be a critical factor underlying the differences in GluA1 and GluA2 temporal concentration profiles following cLTP induction. Taken together, our experimental and modeling results reveal a novel role the CNIH-2 local synthesis plays in diversifying glutamate receptor complexes in dendrites.

## Results

### Active transport of AMPARs can explain their homogeneous distribution along the dendrites

Diffusion and active transport are known trafficking mechanisms that shape AMPAR distribution along the neuronal dendrites. However, the local transport of AMPAR subunits is not very well understood. To understand the role of local synthesis in AMPAR availability in dendrites, we first investigated the distribution of the mRNAs coding for AMPAR subunits. We analyzed fluorescent insitu hybridization data to detect *Gria1* and *Gria2* mRNA-coding, respectively, for GluA1 and GluA2 AMPAR subunits (Fig. 1 and Fig. S1). First, we quantified the endogenous levels of *Gria1* and *Gria2* mRNA molecules in hippocampal cultured neurons using fluorescent in-situ hybridization (Fig. 1 and Fig. S1, and Table 1 for experimental techniques). We calculated the fractions of *Gria1* and *Gria2* localized in the cell body and in dendrites and compared them to *Camk2a* mRNAs^3, 5^ detected in the same neuron. While a majority of *Camk2a* mRNA was detected in dendrites (over 50% in both experimental datasets), both *Gria1* and *2* were localized primarily in neuronal somata (74% and 82%, respectively), with a smaller dendritic mRNA fraction of 26% and 18%, respectively (Fig. 1A-B and Fig. S1A-C). Our previous work shows that although *Camk2a* mRNA distribution declines towards the distal tips of dendrites, it still provides a sufficient pool for local synthesis^5^. To understand if the *Gria1* and *2* mRNA distributions behave similarly, we fitted the mRNA dendritic distributions with the exponential functions (Fig. 1C and Fig. S1C, Methods) and compared their relative abundance at 100 *µm* dendritic distance (1C and Fig. S1C, inset). At a dendritic distance of 100 *µm*, the distribution of *Camk2a* mRNA molecules showed a moderate decrease of 42%, while the *Gria1* and *Gria2* mRNA concentrations dropped by 96% and 98%, respectively (Fig. 1C, inset). The rapid decline and relatively low levels of both *Gria1* and *Gria2* mRNA with distance to soma challenge the possibility that local synthesis is at the core of maintaining AMPA receptor availability at distal sites as it does for *Camk2a*. We note that previous studies reported the local synthesis of AMPAR subunit GluA1 upon BDNF stimulation^30^ and secreted Amyloid precursor protein-alpha mediated LTP^31^ in proximal dendrites up to a few tens of microns. These results are consistent with the presence of *Gria1* and *Gria2* in dendrites in the vicinity of the soma and do not contradict the lack of local GluA1 and GluA2 synthesis in more distal locations observed in our analyses.

**Figure 1:**
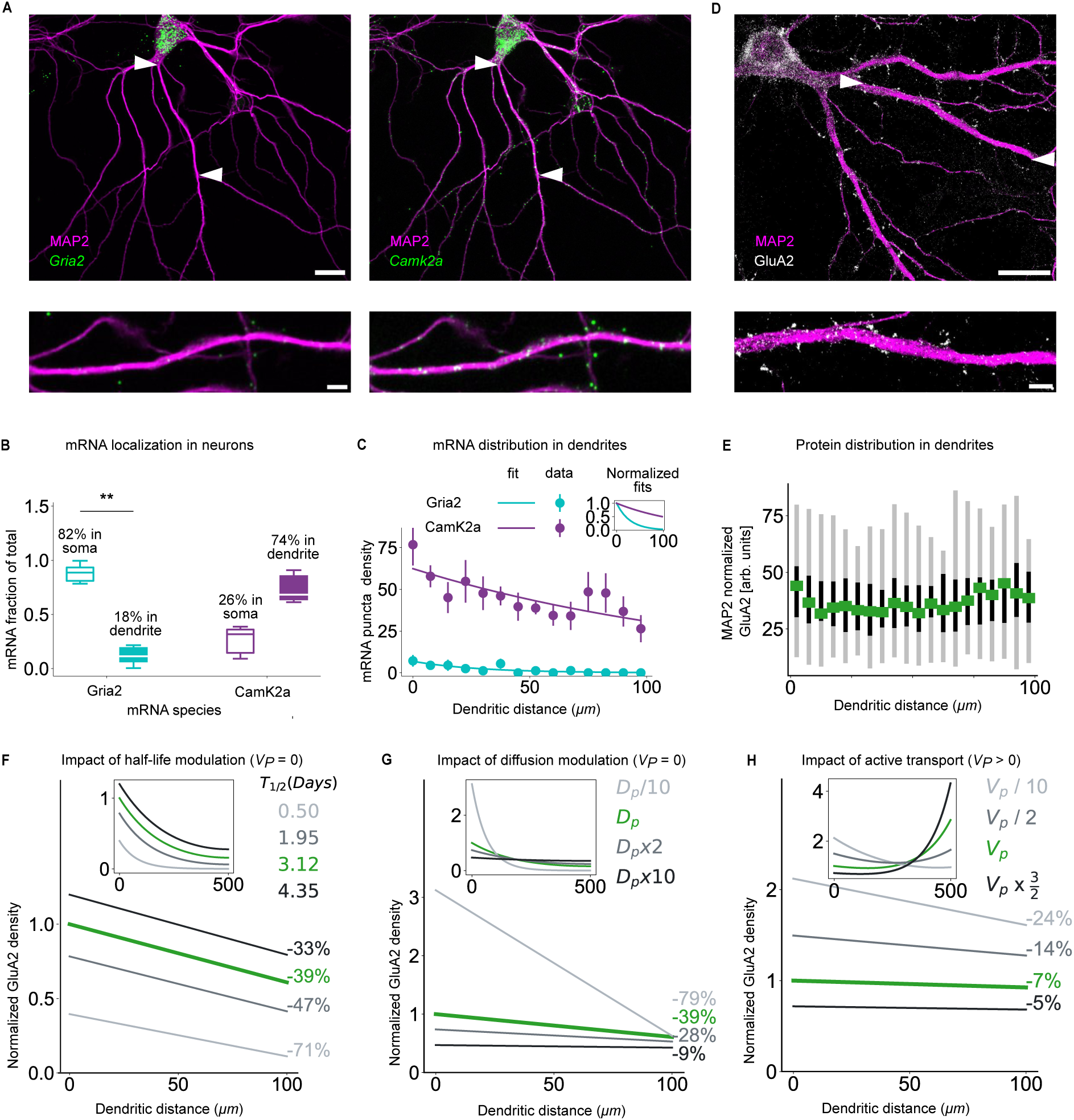
Active transport of AMPAR GluA2 subunits compensates for the lack of their local translation. **(A)** *Top*: Cultured rat hippocampal neurons at 18-21 DIV labeled using fluorescence *in situ* hybridization to detect *Gria2* (left, green) and *Camk2a* mRNAs (right, green) and immunostained for MAP2 (magenta). Scale bar = 20 *µ*m. *Bottom*: Zooming into a representative dendrite shows a smaller number of *Gria2* mRNA (left) compared to *Camk2a* mRNA (right). Scale bar = 5 *µ*m. **(B)** Somatic (hol-low) and dendritic (filled) fraction of total mRNA for *Gria2* and *Camk2a* is computed as a fraction in a compartment with respect to the total mRNA count in a neuron. Dendritic fractions were tested against 0.5 using two-sided one-sample Wilcoxon signed-rank tests (*Gria2*: *W* = 1, *P*_adj_ = 0.00586, *n* = 11 cells; *Camk2a*: *W* = 0, *P*_adj_ = 0.125, *n* = 5 cells). The between-mRNA comparison used a two-sided Wilcoxon signed-rank test on shared cells (*W* = 0, *P*_adj_ = 0.125; *n* = 5 paired cells). Holm adjustment was applied across the three comparisons. Boxes show the 25th–75th percentiles, with median lines and whiskers extending to the extreme values; Percentages indicate mean somatic and dendritic fractions. **(C)** Exponential fit of mRNA puncta density distribution for *Gria2* (*n* = 12 dendrites from 7 cells, exponent *Gria2* = −0.04 0.009) compared to the *Camk2a* distribution (*n* = 10 dendrites from 5 cells, exponent *Camk2a* = −0.006 0.001), see Methods for mRNA exponential fit. *Inset*: Normalized fitted *Gria2* mRNA distribution compared to *Camk2a*. Data points are presented as mean SEM. **(D)** *Top*: Cultured rat hippocampal neurons at DIV 18-21 processed for antibody labeling of GluA2 (white) and FI MAP2 (magenta). *Bottom*: Zoom into a representative dendrite shows a homogeneous GluA2 protein distribution along the first 100 *µ*m. Scale bar = 20 *µ*m. **(E)** MAP2 normalized GluA2 intensity (*n* = 35 dendrites, 5 *µ*m bins, black vertical lines show the 25th–75th percentiles, with median represented as green square and whiskers extending to the extreme as gray bars). **(F)-(H)** GluA2 distributions based on molecular dynamics model (Methods). **(F)**, **(G)** Active transport *V_p_* = 0. **(F)** The half-life (*T*_1*/*2_) of GluA2 protein increased over the reported maximum of 3.12 days cannot provide constant GluA2 distribution. **(G)** Increasing the reported diffusion constant (*D_p_* = 0.1 *µm*^2^*/s*) of GluA2 by a factor of 10 leads to a 9% GluA2 decrease at 100 *µ*m but does not support the increase at 500 *µ*m (inset). **(H)** Active transport of *V_p_* = 1.4 10*^−^*^3^ *µ*m/s leads to an observed 7% decrease in GluA2 protein concentration and increases concentration at 500 *µ*m (inset). **: p *≤* 0.01. Source data are provided as a Source Data file.

We investigated whether the density of GluA2-containing receptors in cultured hippocampal neurons (21 days *in vitro*) aligned with previously reported trends. Using protein antibody labeling, we visualized total GluA2-containing AMPARs after fixation and permeabilization (Fig. 1D). Receptor density was analyzed in the first 100 *µm* of primary dendrites by measuring fluorescence intensity over 5 *µm* segments. Our data did not show any major trend across the initial 100 *µm* of dendritic length. To explore the molecular mechanisms underlying the observed AMPAR distribution, we developed a computational model incorporating intracellular vesicle velocity (*V_p_*), diffusion rate (*D_p_*), and half-life (*λ_p_*) (Eq (4), Methods), using the biologically plausible parameter ranges (Table 2). Fitting the experimentally observed GluA2 distribution (Fig. 1E) to the model (Eq (4), Methods) revealed a marginal 7% decrease at 100 *µm* distance from the soma (Fig. 1E). Previous experimental studies examining AM-PAR distribution at a longer distance (100-250 *µm*)^32^ reported up to a two-fold increase in AMPAR-mediated currents along CA1 apical dendrites and attributed them to a proportional increase in AM-PAR concentration^32^. This phenomenon, known as distance-dependent scaling, suggests an accumulation of AMPARs toward the distal dendritic location. A subsequent study^33^ confirmed distance-dependent scaling specifically in AMPARs containing the GluA2 subunit. Building on these findings, we next examined how variations in half-life (*T*_1*/*2_), diffusion (*D_p_*), and active transport (*V_p_*) within biologically reported ranges (Table 2) might shape the AMPARs distribution. Specifically, we investigated parameter combinations that could reconcile our experimentally observed near-uniform AMPAR distribution close to the soma with the increasing AMPAR concentration trends reported in previous studies^32, 33^.

**Table 2:**
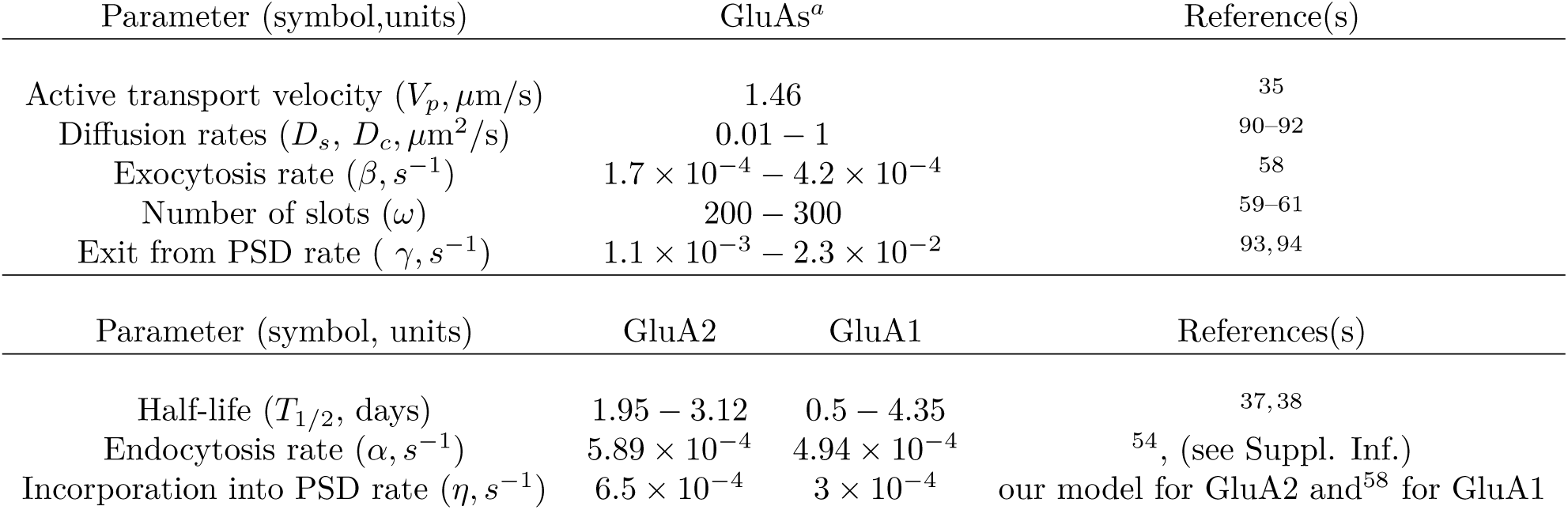
Experimentally reported parameters measured in dendrites of hippocampal cultured neurons; a) includes values that are comparable for both GluA1/2 subunits or if the value is reported only for one of the subunits.

| Parameter (symbol,units) | GluAs<sup>a</sup> |  | Reference(s) |
|---|---|---|---|
| Active transport velocity (V<sub>p</sub>, µm/s) | 1.46 |  | <sup>35</sup> |
| Diffusion rates (D<sub>s</sub>, D<sub>c</sub>, µm<sup>2</sup>/s) | 0.01 − 1 |  | <sup>90–92</sup> |
| Exocytosis rate (β, s<sup>−1</sup>) | 1.7 × 10<sup>−4</sup> − 4.2 × 10<sup>−4</sup> |  | <sup>58</sup> |
| Number of slots (ω) | 200 − 300 |  | <sup>59–61</sup> |
| Exit from PSD rate ( γ, s<sup>−1</sup>) | 1.1 × 10<sup>−3</sup> − 2.3 × 10<sup>−2</sup> |  | <sup>93, 94</sup> |
| Parameter (symbol, units) | GluA2 | GluA1 | References(s) |
| Half-life (T<sub>1/2</sub>, days) | 1.95 − 3.12 | 0.5 − 4.35 | <sup>37, 38</sup> |
| Endocytosis rate (α, s<sup>−1</sup>) | 5.89 × 10<sup>−4</sup> | 4.94 × 10<sup>−4</sup> | <sup>54</sup>, (see Suppl. Inf.) |
| Incorporation into PSD rate (η, s<sup>−1</sup>) | 6.5 × 10<sup>−4</sup> | 3 × 10<sup>−4</sup> | our model for GluA2 and<sup>58</sup> for GluA1 |

Previous modeling studies have shown that the active transport velocity strongly affects the distribution of molecules along the dendrite^5, 34^. AMPARs are composed of transmembrane proteins; thus, they are transported in vesicles in the cytoplasm. AMPAR-containing vesicles are actively moved inside the cytoplasm using microtubule-mediated transport. Those vesicles containing AMPARs move bidirectionally at a speed in the range of 1.46 *µm/s* in both anterograde and retrograde directions^35^. However, experimental studies do not report a net anterograde active transport velocity range for nonmodified receptors. Therefore, we first considered the active transport velocity an unknown parameter of our mathematical model (Supplementary Information). In the next two steps, we assumed that the net active transport was equal to zero (*V_p_* = 0) (anterograde and retrograde transport fully compensate for one another) and analyzed whether the half-life and diffusion parameter variations can explain the experimentally observed distribution of GluA2 receptors, which was approximately constant near the soma and should increase towards the dendritic end.

First, we analyzed how different values of the protein half-life parameter (*T*_1*/*2_) impact the shape of the GluA2 distribution. Since the range of experimentally reported diffusion parameters spans two orders of magnitude, varying from 0.01 to 1 *µm*^2^*/s* (Table 2), we first set an intermediate value of *D_p_* = 0.1 *µm*^2^*/s*, from^36^ in our simulations. We then assumed that the half-life of the GluA2-containing AMPARs was in the range from 1.95 to 3.12 days, as reported in previous experimental studies^37, 38^. When we set the half-life to 1.95 days, GluA2 concentration dropped by 47% at 100 *µm* dendritic distance. This drop was significantly reduced to 39% when setting the GluA2 half-life to its maximum reported value of 3.12 days. The larger half-life values lead to a longer lifetime of the corresponding molecules, allowing a larger number of them to reach further dendritic locations via diffusion. Nonetheless, a decrease of 39% in GluA2 concentration at 100 *µm* is much higher than our experimentally observed decrease of 7%. Further increasing the GluA2 half-life to 4.35 days (half-life reported for GluA1, Table 2) still resulted in a 33% drop at 100 *µm* dendritic distance (Fig. 1F, Fig. S2A) which further decreased towards the end of model dendritic length (Fig. 1F, inset, Fig. S2A, inset). While increasing the GluA2 half-life did raise its concentration in dendrites, it could not reproduce the experimentally measured GluA2 distribution.

We next set GluA2 half-life to the maximally reported value of 3.12 days^37^ and explored how modulating the receptor diffusion in cellular membranes could shape its distribution. Interestingly, while diffusion is a rather fast parameter (see Table 2), because it is bidirectional, it is relatively slow at moving receptors through the long distances of dendritic arbors^39^. In addition, at the cell surface, dendritic spines further slow down receptor trafficking^40^. Indeed, AMPARs can diffuse in the plasma membrane at a rate up to 1 *µm*^2^*/s*^41,42^. However, the diffusion drops to 10*^−^*^2^*µm*^2^*/s* in synaptic compartments^43^. In addition, receptors are also highly mobile in intracellular organelle membranes and particularly in the endoplasmic reticulum that forms a continuous organelle throughout the neuronal dendritic arbor^44^. To understand the impact of diffusion on the distribution of GluA2, we analyzed the diffusion constants in the range from 10*^−^*^2^ to 1 *µm*^2^*/s* starting from the experimentally reported diffusion constant of *D_p_* = 0.1 *µm*^2^*/s*^36^. For the experimentally reported diffusion constant of *D_p_* = 0.1, we observed a 39% GluA2 decrease at a 100 *µm* dendritic distance (Fig. 1G, Fig. S2B). We divided the diffusion constant by a factor of 10, as reported by^43^, which led to a large drop of 79% in GluA2 concentration (Fig. 1G). Increasing the diffusion constant by a factor of two led to a 28% concentration reduction. Multiplication of the diffusion constant *D_p_*by a factor of 10 led to a 9% decrease at 100 *µm* dendritic distance. This decrease is comparable with our experimentally observed reduction in GluA2 concentration (Fig. 1G). However, this value corresponds to the upper bound of the experimentally reported value of diffusion rate in the dendritic shaft, and the presence of spines leads to a lower net diffusion rate^45^. In addition, similar to the effect observed for half-life (*T*_1*/*2_) values at a larger dendritic distance, different diffusion constants also resulted in a decrease towards the distal tip in our model (Fig. 1G, inset, Fig. S2B, inset). However, these model predictions differ from previous experimental reports on AMPAR distribution at longer distances (100 350 *µm*). Since increasing neither the GluA2 half-life nor diffusion coefficient was sufficient to reproduce distance-dependent scaling of AMPARs, we next considered the possibility that the active transport velocity of GluA2 along microtubules plays a role. We had previously set the active transport velocity to be zero as transport along microtubules in dendrites is known to be bidirectional^46^. In contrast, in the next step, we set the active transport velocity *V_p_* to a positive value, resulting in a biased anterograde movement of intracellular vesicles toward the dendritic tips.

To test this idea, we set the receptor half-life to 3.12 days and fitted the active velocity constant *V_p_* and the diffusion constant *D_p_* to first match our experimentally measured GluA2 fluorescence distribution. The best fit active velocity value was *V_p_* = 1.4 10*^−^*^3^ *µm/s*, and the diffusion constant was *D_p_* = 0.22 *µm*^2^*/s*. With these physiologically plausible values for our parameters, we obtained a distribution of GluA2-containing receptors that displayed a 7% drop at 100 *µm* dendritic distance (Fig. 1H, Fig. S2C). Strikingly, the best-fitted model also predicted a steady increase in receptor concentration that reached 180% at 500 *µm* dendritic distance (Fig. 1H, inset). To understand how a lower value of the active transport velocity might affect the global distribution of GluA2 in our model branch, we divided its value by a factor of 2 and 10. As the active transport velocity was lowered, we observed an accumulation of receptors near the somatic regions (Fig. 1H). When dividing the active transport velocity by a factor of 10, we observed that the gradual increase of GluA2 concentration in distal dendrites was lost (Fig. 1H, inset). Interestingly, multiplying the fitted active transport parameter by a small factor of 3/2 led to a 500% larger GluA2 concentration towards a dendritic tip, showing that even minimal increases in the active transport velocity can lead to a large accumulation of GluA2 molecules at distal dendritic sites (Fig. 1H, inset and Fig. S3).

In conclusion, our experimental data show that *Gria1* and *Gria2* mRNAs, coding for the two most abundant AMPAR pore-forming subunits GluA1 and GluA2, are predominantly translated in the somatic compartment under basal conditions. This observation reduces the likelihood that local translation underlies the reported increase in GluA2 concentration at distal dendritic locations. In our simulations, only by modulating the active transport velocity of GluA2 intracellular vesicles could we recapitulate the experimentally observed distribution of GluA2 along dendrites. Since changes in AM-PAR numbers close to and at synapses are a crucial step during the expression of synaptic plasticity, we then explored how exoand endocytosis shape AMPAR distribution in the plasma membrane and cytoplasm.

### Endogenous concentrations of AMPARs in the cytoplasm, cell membrane, and postsynaptic density

The simulations presented in the previous section indicate that active transport is crucial for delivering AMPARs to distal dendritic sites. Since the number of AMPARs in the postsynaptic density (PSD) correlates with the synaptic strength, our next goal was to understand the mechanisms under-lying AMPAR insertion in the PSD under basal and plasticity conditions. To this end, we analyzed AMPAR trafficking between the cytoplasm (i.e., intracellular compartment), the plasma membrane (i.e., neuronal surface), and the PSD.

AMPAR exocytosis occurs near the synaptic site (spine head or local dendritic shaft), where the AM-PARs can enter the PSD via lateral diffusion in the plasma membrane^47^. Thus, modeling the rate and location for endo- and exocytosis and synaptic trapping can provide a better understanding of the impact of these processes on synaptic plasticity. Notably, the trafficking speed and the residence time (lifetime in a compartment) of transmembrane proteins differ when they are in the plasma membrane or the cytoplasm^40^. Thus, modeling intracellular and surface proteins separately is crucial to estimate their redistribution accurately. Here, we extended the previous version of the model to examine AMPAR distribution in different dendritic compartments: the cytoplasm, the plasma membrane, and the PSD. With this model, we analyzed how endocytosis, exocytosis, and diffusional trapping at PSD shape the AMPAR distribution between the three compartments. We distinguish between mobile AMPARs in the cytoplasm (*P_c_*), mobile receptors in the plasma membrane in extrasynaptic regions (*P_s_*), and the immobile receptors at PSDs (*P*_psd_)

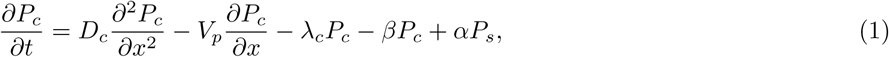

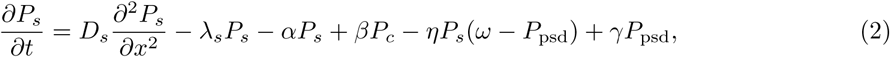

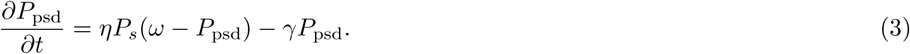

In this model, the intracellular diffusion (*D_c_*), net motor-based transport (*V_p_*), and receptor degradation rate (*λ_c_*) shape the distribution in the cytoplasm along the dendrites. Furthermore, we assumed that the AMPARs in the plasma membrane move exclusively via lateral diffusion (*D_s_*) - Brownian movement. For local trafficking between surface and cytoplasm, we introduced coupling terms of exocytosis and endocytosis rates (*β* and *α*, respectively). Synaptic uptake can impact the availability of the AMPARs at different dendritic locations. Surface receptors can get trapped by binding to anchoring proteins (e.g., PSD-95) in the PSD with an incorporation rate of *η*. Trapped receptors will eventually unbind from anchoring proteins and exit the PSD at the rate of *γ*. Finally, we assumed that each synapse has a maximum capacity (*ω*) for holding AMPARs, determined by the number of the so-called “slots” that are postsynaptic density structures consisting of proteins such as PSD-95 that anchor AMPARs in the PSD and restrict their movement^29, 48, 49^. For simplicity, we did not consider the nanoscale organization of these slots, which can influence the diffusional trapping of AMPARs inside the PSD (see^50^). To constrain parameters for our model, we performed live-cell HaloTag labeling of surface and intracellular GluA2 endogenously tagged with HaloTag using CRISPR/Cas9-mediated gene editing (Fig. 2A;^51^). Surface Halo-GluA2 was labeled using an impermeable HaloLigand, which saturates all extracellular HaloTags^52^. Subsequently, intracellular Halo-GluA2 was labeled with a permeable HaloLigand. The HaloLigand dyes Janelia Fluor JF549 and JF635 that were used to label GluA2 do not have the same brightness, which leads to a biased measurement of GluA2 distribution. To overcome this limitation, we labeled half of the neurons with JF549i (impermeable HaloLigand) and then JF635, and the other half with JF635i and then JF549 thereby effectively mitigating any biases. We then determined the trafficking parameters under basal conditions by fitting the distribution of the GluA2-containing AMPARs at the surface and in the cytoplasm (Fig. S4) to the steady state distribution Eqs. (9)-(11) of the model in Eqs. (1)-(3). We first computed the exocytosis and endocytosis rates, which ensure continuous cycling of the GluA2-containing AMPARs between the plasma membrane and the cytoplasm. To determine the exocytosis rate, we used our theoretical prediction in Eq. (19) showing that the ratio between the exoand endocytosis rates (*β*/*α*) is given by the ratio between the total surface to intracellular GluA2 fluorescence intensity (sGluA2/iGluA2). To determine this ratio, we first computed it in two sets of neurons (Fig. 2B, C). In the dataset where JF549i was used to label the surface GluA2, the mean ratio of sGluA2 to iGluA2 was comparable in dendrite (mean std: 4.98 2.5) and somata (mean std: 5.2 8.61, W = 587, P = 0.0571, see Fig. 2B). On the other hand, in the dataset where we used JF635i to label surface GluA2 the ratio of surface to intracellular GluA2, was significantly lower in somata than dendrites (Fig. 2C, somata vs dendrites: 0.08 0.06 vs 0.33 0.13, W = 0, P = 2.78 10*^−^*^7^). Finally, to account for the biases due to the different HaloTag dyes used between the two datasets we computed the geometric mean of the two datasets (see Methods section). Finally, our calculations led to an estimated mean ratio of 1.29 (95% CI (1.17, 1.41)) in dendrites (Fig. 2D). The estimated ratio in somata (mean 0.64, 95% CI (0.45, 0.83)) was lower than in dendrites. The observation that the ratio is lower in somata compared to dendrites has also been reported by Rathje and colleagues^53^. Using this ratio in dendrites (Fig. 2D) and the formula in Eq. (19), we computed the exocytosis rate to be *β* = 7.73 10*^−^*^4^*s^−^*^1^, which is larger than the endocytosis rate *α* = 6 10*^−^*^4^*s^−^*^1^ that we adopted from experiments reported by Rosendale and colleagues^54^. Interestingly, this estimated ratio was comparable across different dendritic lengths (Fig. S5).

**Figure 2:**
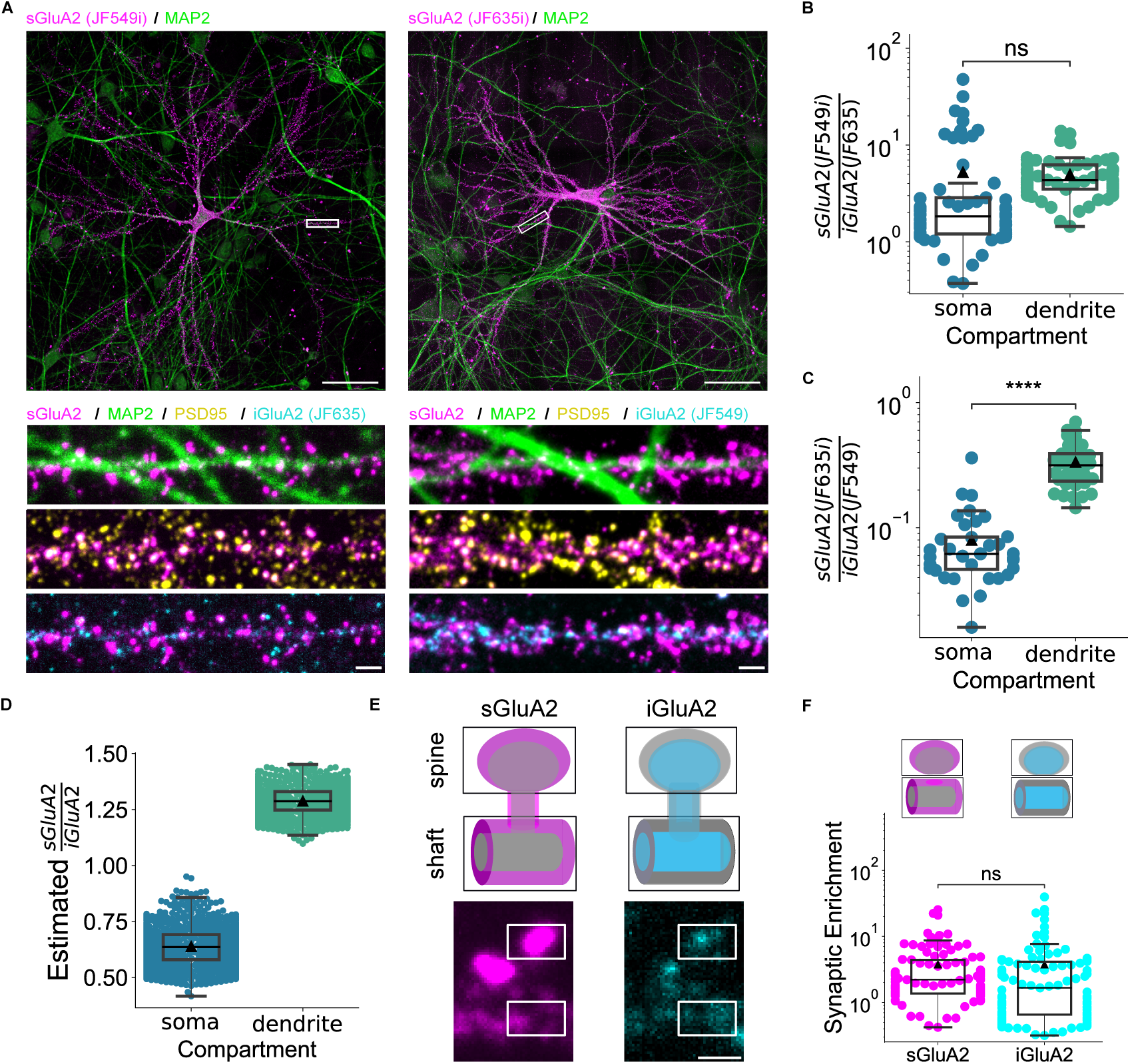
The endogenous distribution of surface and intracellular GluA2-containing AMPARs. **(A)** *Top*: Cultured rat hippocampal neurons at 21 DIV visualized using CRISPR/Cas9 by endogenously tagging GluA2 with HaloTag. Dendrites are labeled using immunostaining to detect MAP2 (green). Surface HaloTag-GluA2 (magenta) was live-cell labeled using the cell-impermeable dyes JF549i (left) and JF635i (right). Scale bars 50 *µ*m. *Bottom*: Zooms of representative dendrites labeled for surface HaloTag-GluA2 (magenta), MAP2 (green), PSD95 (yellow) and intracellular HaloTag-GluA2 (cyan). Scale bars 2 *µ*m. **(B)** Fluorescent intensity ratio between surface (JF549i) and intracellular (JF635) GluA2 was equal to 5.2 8.61 in somata (mean std) and 4.98 2.5 in dendrites (mean std, *W* = 587, *P* = 0.0571, *n* = 57 paired cells). **(C)** Fluorescent intensity ratio between surface (JF635i) and intracellular (JF549) GluA2 was equal to 0.08 0.06 in somata (mean std) and 0.33 0.13 in dendrites (mean std, *W* = 0, *P* = 2.78 10*^−^*^7^, *n* = 35 paired cells). **(D)** The estimated mean of sGluA2 to iGluA2 as the geometric mean of the ratios in **(B)** and **(C)** yielded a mean value of 0.64 (95% CI (0.45, 0.83)) in somata and mean of 1.29 (95% CI (1.17, 1.41)) in dendrites, CI was estimated via bootstrapping 1000 subsamples. **(E)** Schematic of the surface (top, left, magenta) and intracellular areas (top, right, cyan) of a synapse. Example images (bottom) show dendritic segments with spine and corresponding shaft ROIs (scale bar 1 *µ*m). **(F)** The surface (top, left) and intracellular (top, right) synaptic enrichment (SE) was computed as the background subtracted mean fluorescent intensity corresponding to the PSD95 spine-associated ROIs divided by the shaft (magenta for the surface and cyan for the cytoplasm area, respectively SE_sGluA2_ = 3.82 4.34, SE_iGluA2_ = 3.74 6.08, *W* = 1985, *P* = 0.351; *n* = 94 paired cell averages). Significance of the pair-wise comparisons were assessed using a two-sided Wilcoxon signed-rank test. Boxes indicate the interquartile range, horizontal lines indicate medians, and black triangles indicate means. Whiskers extend to the most extreme values. Individual points represent cell-level measurements (B, C, F). Source data are provided as a Source Data file.

We note that the exact sites of AMPAR exo- and endocytosis will likely impact the speed of the AMPAR insertion and removal from the PSD, thus affecting the AMPAR responses to plasticity cues. However, the precise localization of those sites (i.e., near the PSD or on the dendritic shaft) remains an active source of debate. Exocytosis of AMPARs has been reported to occur primarily at peri-synaptic sites^55^, on the dendritic shaft^56^, or both^57^. Importantly, the closer the exocytosis sites to the PSD, the shorter the distance AMPARs need to diffuse to reach the PSD and participate in synaptic transmission. Furthermore, spine necks act as a bottleneck that limits the surface diffusion of AMPARs^58^ and, hence prolongs response times. Similarly, the exact site of the AMPAR endocytosis is still a topic of active research, with recent evidence suggesting AMPAR internalization at the peri-synaptic endocytic zone^54^. For simplicity we decided to use the dendritic shaft as the only site of endocytosis and exocytosis in our model.

AMPARs enter the PSD via lateral diffusion following exocytosis to the cell surface. Once AMPARs reach a PSD, they can be trapped through interactions with scaffolding proteins. Intracellular signaling pathways, including those involving phosphorylation, regulate these interactions. AMPARs leave the PSD by unbinding from scaffolding sites and diffusing away. To inform the model parameterization of GluA2-containing AMPAR incorporation into the PSD, we analyzed their enrichment in spine-associated regions relative to neighboring dendritic shafts (Fig. 2F). We calculated the ratio of background-subtracted mean fluorescence intensity in PSD95-defined spine-associated ROIs to that in neighboring dendritic shaft ROIs. The mean ratios were 3.82 4.34 for surface GluA2 and 3.74 6.08 for intracellular GluA2 (mean SD), with no statistically significant difference detected between the two distributions (*p* = 0.351). These measurements describe relative fluorescence within the selected ROIs; they do not directly quantify receptor concentration, the fraction of receptors located within PSDs, or receptor immobilization. Additionally, an independent dataset based on live-cell surface labeling with ATTO643-conjugated Fab fragments derived from the monoclonal anti-GluA2 antibody 15F1 yielded a comparable surface spine-associated/shaft fluorescence ratio in primary hippocampal neurons (Fig. S6), supporting consistency across labeling approaches.

We used the surface fluorescence ratio as an approximate phenomenological constraint on baseline receptor partitioning in the model. Its correspondence to the model’s PSD-to-extrasynaptic receptor ratio is a simplifying assumption, because confocal imaging does not distinguish PSD-localized receptors from receptors elsewhere on the spine membrane. With the exit rate fixed at *γ* = 2.3 10*^−^*^2^ s*^−^*^1^, we fitted the incorporation parameter to obtain *η* = 6.5 10*^−^*^4^ s*^−^*^1^. This estimate is conditional on the assumed mapping between fluorescence and model receptor partitioning, the exit rate, and the prescribed number of scaffolding slots. We fixed the slot number at 250, within the range reported for scaffolding sites^59–61^. Under this parameterization, approximately 30% of slots were occupied at base-line, consistent with the experimental estimate reported by Clavet-Fournier and colleagues^62^. Finally, we fitted the steady state equations in Eq. (13) to estimate global transport parameters (diffusion constants *D_s_*, *D_c_*, and net active transport velocity *V_p_*) from the normalized GluA2 fluorescent intensity (Fig. S4). We obtained the best-fit values for diffusion coefficients *D_s_* = 0.01 *µm*^2^*/s*, *D_c_* = 0.4 *µm*^2^*/s*, and the active transport velocity of *V_p_* = 0.9 *×* 10*^−^*^3^ *µm/s*. The model parameters obtained here are comparable to those obtained for the model shown in Fig. 1. With these fitted parameters our extended model predicted a roughly 15% increase in total AMPAR density at 500 *µm* which is in line with the increase in AMPAR density measured in-vivo (Fig. S13,^63^).

Overall, using the experimentally measured AMPAR distribution and extended molecular dynamics model, we characterized the local and global trafficking parameters of endogenous AMPARs. The inferred exocytosis rate (*β*) constant exceeded the prescribed endocytosis (*alpha*) rate constant, while integrated exocytic and endocytic receptor fluxes balanced at steady state. Notably, the exocytosis-to-endocytosis ratio remains approximately constant along the dendrite. The coexistence of two regimes “live longer - travel slower” in the neuron plasma membrane and “live shorter - travel faster” in the cytoplasm - seem to balance each other such that it enables GluA2-containing AMPAR availability at distal dendritic sites and their stability near synapses. Next, we used the molecular dynamics model to study temporal changes in AMPAR concentration due to the regulation of local trafficking upon plasticity induction.

### GluA1-homomeric and GluA2-containing AMPARs elicit distinct temporal response upon LTP

Modeling AMPAR temporal dynamics is crucial for understanding the time scales of synaptic plasticity. One of the challenges in understanding AMPAR temporal dynamics is that the onset and duration of LTP-related processes depend on the specific subunit composition of AMPARs involved. So far, we have focused our experimental investigations mainly on GluA2-containing AMPARs. However, recent studies report that the GluA2-lacking or calcium-permeable AMPARs are essential for LTP^64, 65^. Calcium-permeable AMPARs get trapped rapidly during a short and transient window after the start of LTP induction - we note that the most abundant calcium-permeable AMPAR poreforming complex is a GluA1 homotetramer^1^. To predict and compare the temporal responses of the GluA2-containing AMPARs and GluA1 homomers, we compared our model predictions with experimental studies. We used data from two independent studies to isolate the unique behavior of GluA1 homomers and GluA2-containing receptors.

First, we used our model to simulate the experimental data from Tanaka and Hirano^66^ in which the overexpression of GluA1 promoted the formation of GluA1-homomeric AMPARs. We started with the steady state distribution of GluA1-homomeric AMPARs using the parameters in Table 3, and then simulated cLTP induction. During cLTP induction, these receptors increased in the plasma membrane and, to a greater extent, in the PSD, peaking at the end of the cLTP stimulation using glycine. Following the chemical stimulation, the concentration of GluA1-homomeric receptors gradually decreased. Incorporating the cLTP protocol from^66^ in our model, we tested the model’s ability to recapitulate the temporal response of the GluA1 homomers upon cLTP induction (see Fig. 3A and Methods).

**Figure 3:** A constant increase in exocytosis rate upon LTP can explain temporal dynamics of GluA1-homomeric AMPARs but not GluA2-containing AMPARs. **(A)** cLTP induction led to 3.5-fold increase in exocytosis rate (*β*) (^66^) and 30% increase in synaptic hopping-in rate (*η*) of GluA1-homomeric AMPARs. The mathematical model matches the amplitude and subsequent decrease in response of GluA1-homomeric AMPARs reported in^65^. **(B)** Concentration of GluA2-containing AMPARs increases within two hours following cLTP induction^62^. Long lasting increase of constant magnitude in the exocytosis rate cannot explain experimental data. Data points are presented as mean SD. Solid lines represent mean response curves (A, B). **(C)** Pairwise relation between parameter change upon LTP induction and model performance against data, (see Fig. S8 and Fig. S9). GluA1-homomeric AMPAR data is from^66^, GluA2-containing AMPAR data is from^62^. Source data are provided as a Source Data file.

**Table 3:**
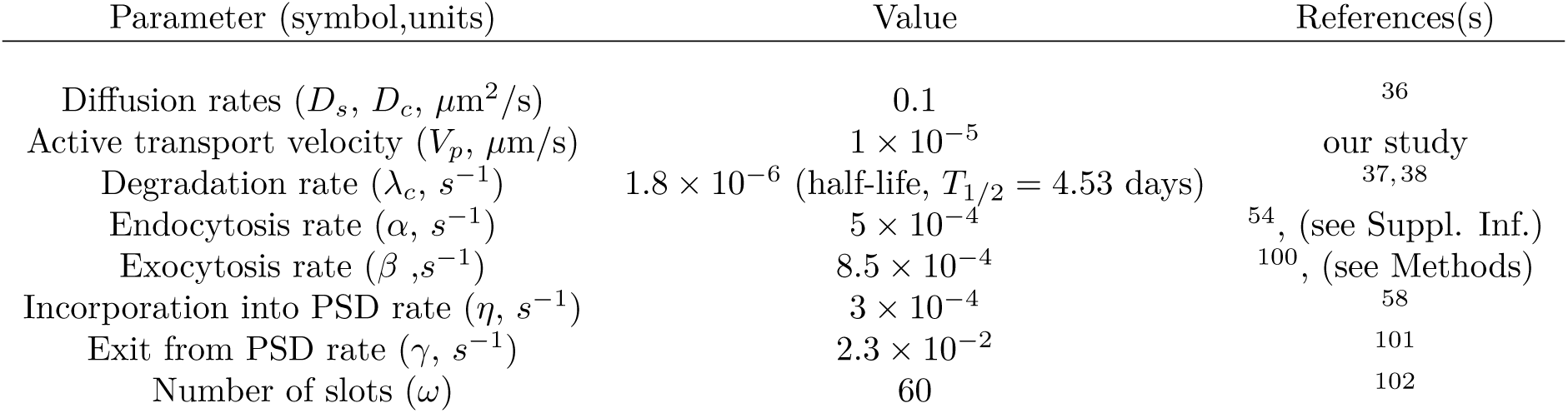
Parameter values used for steady-state, baseline model of GluA1-homomeric AMPARs.

According to Tanaka and Hirano, the GluA1 exocytosis frequency increases immediately after cLTP induction but returns to basal level when glycine is washed out (Fig. 3A, S7)^66^. Importantly, changes in GluA1 exocytosis rate are consistent with other independent studies (e.g.,^57^), indicating that the homomeric GluA1 AMPAR exocytosis rate increases immediately upon cLTP induction and lasts only during the stimulation phase. We simulated an instantaneous 3.5-fold increase of the GluA1 exocytosis rate (*β*) homogeneously throughout the dendrite during the 10 minutes of cLTP induction (Fig. S7). Our simulation showed that a 3.5-fold increase in exocytosis rate for 10 minutes leads to an increase in GluA1 concentration in the plasma membrane and in PSDs (Fig. S8A). However, when modifying the exocytosis rate alone, the model predicted a transiently higher concentration of GluA1 in the dendrite compared to the PSDs, which contradicted experimental observations of Tanaka and Hirano^66^. To address this, we adjusted the model to promote GluA1 incorporation in PSDs by increasing both the exocytosis rate and the PSD incorporation rate (*η*) upon cLTP induction. The prolonged increase in incorporation rate reflects increased receptor binding to scaffold proteins, associated with phosphorylation of GluA1 and stargazin C-terminal domains and increased synapse size^1, 62, 67^. We simulated an instantaneous 1.3-fold increase in the incorporation rate together with a 3.5-fold transient increase in the exocytosis rate (Fig. 3A). Using these parameters, our model was able to replicate the experimentally observed magnitude and temporal profile of GluA1 homomeric AMPARs (Fig. 3A and C). Notably, neither exocytosis nor the incorporation rate up-regulation alone could reproduce the experimentally observed temporal profile of GluA1 surface concentration (Fig. S8A and B). Interestingly, our model predicted that 30 minutes following cLTP induction, the concentration of GluA1 in the plasma membrane in extrasynaptic areas dropped back to the baseline level and decreased below the baseline after approximately one hour while the synaptic GluA1 concentration reached an equilibrium concentration of 15% above the baseline (Fig. 3A).

Second, changes in synaptic efficacy induced by cLTP are known to last for several hours. As described above, these long-lasting changes have been attributed to an increase in the concentration of GluA2-containing AMPARs that, according to a recent report, can last for at least two hours^62^. Clavet-Fournier and colleagues used super-resolution microscopy and antibodies against endogenous GluA2 to monitor the concentration of GluA2 in PSDs after cLTP induction. Their experiments showed increased synaptic GluA2-containing receptors that persisted beyond the stimulation period and reached roughly 50% increase two hours after cLTP induction. To replicate these findings, we incorporated the experimental data for GluA2-containing receptors into our model^62^ (Fig. 3B). However, as we observed in the case of GluA1, increasing the exocytosis and PSD incorporation rates enhanced receptor concentration in PSDs during stimulation but was insufficient to sustain the increased concentration beyond the stimulation period (Fig. 3B, Fig. S9). In fact, the model failed to match both the kinetics and large concentration increase of GluA2-containing receptors (Fig. 3B and C). This suggests that additional mechanisms with effects lasting beyond the stimulation phase were required for the prolonged GluA2 accumulation in PSDs.

As an initial attempt to adjust our model, we investigated whether increasing the exocytosis rate beyond the stimulation phase can lead to a long-lasting increase in GluA2-containing AMPAR concentration in PSD (Fig. S9). Our model predicted that the increase in the receptor concentration lasted roughly as long as the increase in the exocytosis rate. Moreover, a direct change in exocytosis rate showed faster kinetics of GluA2-AMPAR change compared to the data. This suggests that in the case of GluA2-containing receptors, the increase in exocytosis rate is, on the one hand, limited or slow and, on the other hand, lasting for several hours despite the arrest of the stimulation. Strikingly, delayed, and long-lasting responses are typically expected when protein synthesis is recruited upon stimulation^68^. However, we showed previously that GluA2 mRNA is not present at high levels in dendrites (Fig. 1).

### Auxiliary subunit CNIH-2 is locally translated and preferentially mediates GluA2-containing receptor exocytosis

We shifted our focus to other components of the AMPAR macromolecular complexes, namely, auxiliary subunits. Leveraging the translatome dataset published by Glock and colleagues^4^, we confirmed the low levels of AMPAR pore-forming subunits *Gria1-2* mRNAs in dendrites but identified one of the AMPAR auxiliary subunits Cornichon-2 (*Cnih2*) as an abundant dendritic mRNA. To validate this finding, we analyzed *Cnih2* mRNA localization in cultured rat hippocampal neurons using in situ hybridization and compared it to *Camk2a* mRNA. Remarkably, the distribution of *Cnih2* mRNA closely resembled that of *Camk2a* (Fig. 4A). Approximately 50% of *Cnih2* mRNA was localized in the dendrites, similar to the 60% observed for *Camk2a* mRNA (Fig. 4B).

**Figure 4:**
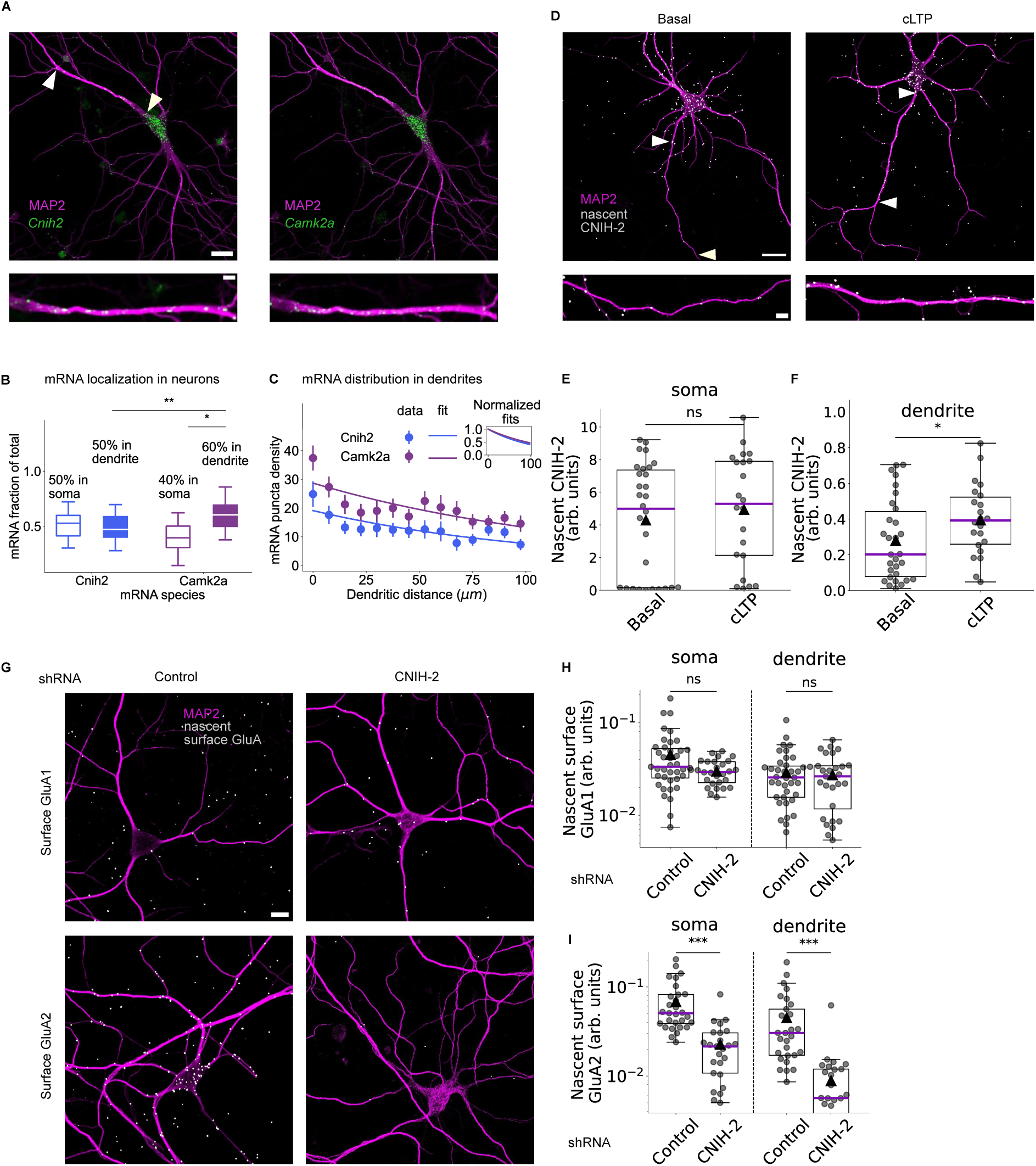
Local translation of CNIH-2 regulates the trafficking of GluA2-containing AMPA receptors to the neuronal surface. **(A)** *Top*: Cultured rat hippocampal neurons (18-21 DIV) processed for FISH against *Cnih2* (left, green) and *Camk2a* mRNA (right, green) and fluorescently immunostained (FI) MAP2 (magenta). Scale bar = 20 *µ*m. *Bottom*: Zooming into a representative dendrite shows a smaller number of *Cnih2* mRNA (left) compared to *Camk2a* mRNA (right). Scale bar = 5 *µ*m. **(B)** Somatic (hollow) and dendritic (filled) fraction of total mRNA for *Cnih2* and *Camk2a*. Dendritic fractions were tested against 0.5 using two-sided one-sample Wilcoxon signed-rank tests (*Cnih2*: *W* = 166, *P*_adj_ = 0.581, *n* = 27 cells; *Camk2a*: *W* = 25, *P*_adj_ = 0.0168, *n* = 19 cells). The between-mRNA comparison used a two-sided Wilcoxon signed-rank test on shared cells (*W* = 21, *P*_adj_ = 0.00508; *n* = 19 paired cells). Holm adjustment was applied across the three comparisons. **(C)** Exponential fit of mRNA puncta density distribution for *Cnih2* (*n* = 44 dendrites, exponent *Cnih2* = −0.009 0.002) compared to the *Camk2a* distribution (*n* = 44 dendrites, exponent *Camk2a* = −0.007 0.002), see Methods for mRNA exponential fit. *Inset*: Normalized fitted *Cnih2* mRNA distribution compared to *Camk2a*. Data points are presented as mean SEM. **(D)** Representative images of hippocampal cultured neurons showing the distribution of the newly synthesized CNIH-2 after 1 hr of metabolic labeling with AHA (*left column*) and an increase of CNIH-2 new proteins in dendrites after cLTP induction (*right column*). Scale bar 20 *µ*m in neuron images, 5 *µ*m in dendrite images. **(E)** Box plots showing the density of newly synthesized CNIH-2 in the soma. **(F)** Same as (E) but in dendrites. Basal and cLTP nascent CNIH-2 densities were compared using two-sided Mann– Whitney *U*-tests in soma (E: *U* = 272, *P* = 0.287) and dendrites (F: *U* = 218, *P* = 0.0389); *n* = 30 basal and 22 cLTP cells per compartment. **(G)** Cultured hippocampal neurons showing newly synthesized surface GluA1 and GluA2 subunits in neurons expressing an shRNA against CNIH-2 or a control shRNA. Without CNIH-2 proteins, newly synthesized GluA2-containing AMPARs are absent at the somatic and dendritic surface. Scale bar 20 *µ*m. (H) Box plots show the abundance of newly synthesized GluA1 (top) and (I) GluA2 (bottom) containing receptors at the cell surface in soma and dendrite. Control shRNA and *Cnih2* shRNA groups were compared using two-sided Mann–Whitney *U*-tests: GluA1 **(H)**, soma (*U* = 566, *P* = 0.804) and dendrites (*U* = 689, *P* = 0.0701; *n* = 39 control and 28 knockdown cells); GluA2 (I), soma (*U* = 672, *P* = 8.04 10*^−^*^8^) and dendrites (*U* = 664, *P* = 1.77 10*^−^*^7^; *n* = 29 control and 25 knockdown cells). *: p 0.05, ***: p 0.001. Boxes indicate the interquartile range, and whiskers extend to the most extreme values. Horizontal lines indicate medians (B, E, F, H and I); black triangles indicate means and individual points represent cell-level measurements (E, F, H and I). Source data are provided as a Source Data file.

Fitting *Cnih2* and *Camk2a* mRNA distributions with exponential decay functions (Fig. 4C, Methods) revealed comparable reductions at 100 *µm* from the soma: 59% for *Cnih2* and 52% for *Camk2a*. Next, we investigated whether CNIH-2 protein was indeed synthesized locally. To assess de novo CNIH-2 protein synthesis, we performed fluorescence non-canonical amino acid tagging coupled with proximity ligation assay (FUNCAT-PLA)^69^. FUNCAT-PLA couples the incorporation of a non-canonical amino acid, azidohomoalanine (AHA), a methionine derivative, during protein synthesis with an antibody-based detection of a protein-of-interest; the coincidence of the signals is detected with proximity ligation assay. Neurons were incubated for 1 hour with AHA (4 mM) or Methionine as a control and then fixed for further processing (see Methods).

FUNCAT-PLA revealed widespread CNIH-2 synthesis in somata and throughout dendrites, with approximately 70% of the nascent protein localized to dendrites (Fig. S10A-D). In the Methionine control condition, the FUNCAT-PLA signal was essentially absent (Fig. S10A-C). Together, these results indicate that CNIH-2 is locally synthesized in the dendrites.

To examine whether CNIH-2 contributes to the activity-dependent trafficking of AMPARs to the neuronal plasma membrane, we applied a cLTP protocol (see Methods) and tracked newly synthesized CNIH-2 proteins using FUNCAT-PLA (Fig. 4D). cLTP induction resulted in a significant, approximately twofold increase in the dendritic CNIH-2 protein levels within an hour (Fig. 4F), while no significant changes were observed in neuronal somata (Fig. 4E). This experiment shows the direct regulation of an AMPAR auxiliary subunit protein level by activity within the dendrites. We next asked how dendritic CNIH-2 synthesis regulates AMPAR trafficking to the neuronal surface. To this end, we optimized the FUNCAT-PLA pipeline^69^ to label nascent GluA1 and GluA2-containing receptors localized in the plasma membrane (see Methods). To investigate whether CNIH-2 synthesis promotes the trafficking of AMPAR to the cell surface, we used a shRNA strategy to reduce *Cnih2* transcript levels. We transfected hippocampal cultured neurons with CNIH-2 shRNA or a control shRNA at 8 days in vitro (DIV). After 11 days of expression of the shRNA, neurons were incubated for 2 hours with AHA (4 mM) or methionine as a control, followed by surface labeling with GluA1 or GluA2 antibodies prior to fixation. The efficiency of CNIH-2 expression knockdown after 11 days of expression was assessed by both in situ hybridization against *Cnih2* mRNA and FUNCAT-PLA to detect nascent CNIH-2 protein (Fig. S11). Reduced CNIH-2 expression had no effect on the surface expression of the GluA1 subunit (Fig. 4G and H) but dramatically reduced the surface expression of GluA2 subunits in both the cell soma and the dendritic arbor (Fig. 4G and I). These findings suggest that CNIH-2 is critical in driving the surface expression of GluA2-containing AMPARs. To determine whether CNIH-2 also promotes surface trafficking of GluA2-containing AMPAR during LTP, we used the shRNA strategy to reduce the *Cnih2* levels and treated cultured hippocampal neurons for 5 min with glycine and picrotoxin in absence of Mg2+ to chemically induce LTP. In neurons expressing CNIH-2 shRNA we did not observe any increase in GluA2 surface expression after cLTP induction (see Fig. S12), indicating that CNIH-2 expression is necessary for both basal and LTP-induced surface insertion of GluA2-containing AMPARs.

### CNIH-2 local synthesis-driven exocytosis can explain the slow and persistent response of GluA2-containing AMPARs upon LTP

Building on the results from the previous section, we then incorporated CNIH-2 in our model as a regulator, under the control of local protein synthesis, of GluA2-containing AMPAR exocytosis rate. Using our previously developed model^5^, we examined steady-state distributions and concentrations of CNIH-2 protein and its mRNA under basal and plasticity conditions (see also Supplementary Information). In the model, we simulated LTP at 10 spines (2% of spines) distributed across our model dendrite. For these spines, a prolonged 2-hour up-regulation of CNIH-2 local synthesis led to an approximately 10-fold increase in CNIH-2 protein concentration in the vicinity of stimulated spines (Fig 5A). While the CNIH-2 concentration, in the vicinity of stimulated spines, peaked during the up-regulation of its local protein synthesis, a 5-fold elevated CNIH-2 concentration was still present 5 hours after the start of the simulation (Fig. 5A).

**Figure 5:**
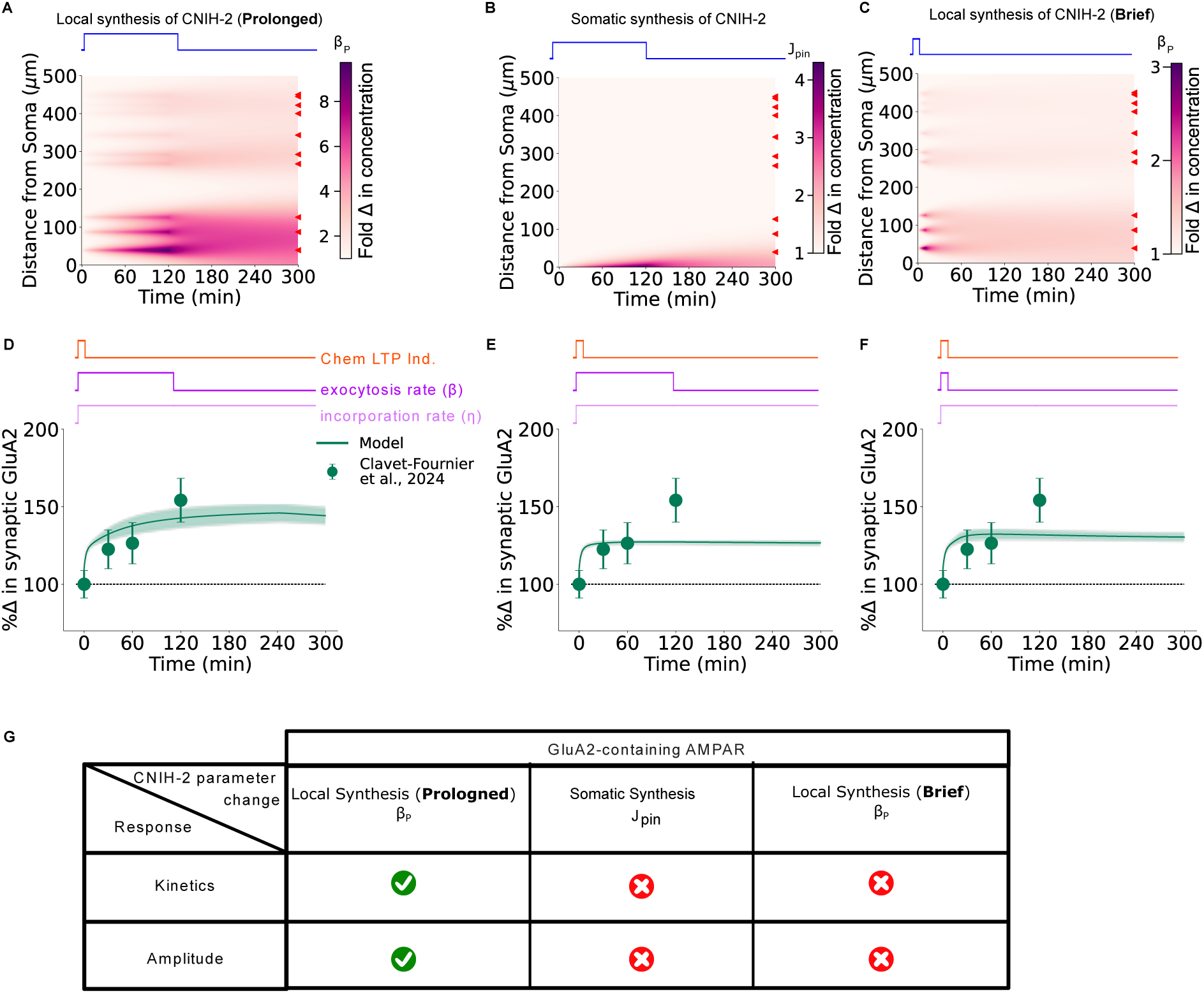
Sustained local synthesis-driven exocytosis can explain the time course of cLTP-induced changes in GluA2-containing AMPARs. **(A, B, C)** Spatiotemporal distribution of CNIH-2 protein following: **(A)** prolonged, 2 hours up-regulation of its local synthesis, **(B)** prolonged, 2 hours up-regulation of somatic flux, **(C)** brief, 10 minutes up-regulation of local synthesis rate. In case of local synthesis upregulation **(A&C)**, the local synthesis rate *β_p_* was increased near the location of stimulated spines (red triangles) which led to increase in CNIH-2 concentration in the vicinity of stimulated spines. On the contrary, somatic flux upregulation did not lead to any considerable increase in CNIH-2 concentration at the stimulated spines. **(D, E, F)** The time- and space-dependent increase in CNIH-2 concentration was used to model an increase in the GluA2 exocytosis rate. In addition, we stepwise increased the synaptic incorporation rate *η* by 70% (*η^cLTP^ /η* = 1.7). **(D)** The resulting GluA2-containing AMPAR distribution matched the temporal response kinetics and the amplitude reported in^62^ only when local CNIH-2 synthesis rate was increased for the prolonged duration. Both **(E)** somatic flux increases and **(F)** brief local synthesis upregulation, did not match the change in synaptic GluA2 profile, with concentration saturation at a much lower concentration than observed in data. Data points are presented as mean SD. Solid line represents mean response curve with the shaded region representing the 95% CI (D, E, F). **(G)** Pairwise relation between CNIH-2 trafficking kinetics upon chemical LTP induction and model performance against data from^62^. Source data are provided as a Source Data file.

In contrast, enhanced CNIH-2 synthesis confined to the somatic region led to a marginal increase in CNIH-2 dendritic concentration of roughly 4-fold in proximal dendrite after 2 hours of up-regulation but, importantly, this increase was reduced to 2-fold after 5 hours (Fig. 5B). Finally, a brief 10-minute up-regulation of CNIH-2 translation led to a moderate 3-fold increase in CNIH-2 concentration after 10 minutes in the proximal dendritic region. This increase dropped to 1.5-fold after 5 hours (Fig. 5C). Next, we incorporated time-dependent changes in CNIH-2 distribution in our mathematical model of AMPAR trafficking. Specifically, we simulated an increase in the exocytosis rate of GluA2-containing AMPARs driven by the CNIH-2 local synthesis combined with a 1.7-fold increase in the PSD incorporation rate (*η*) after cLTP induction (Fig. 5D). Importantly, we assumed that the exocytosis rates (*β*) varied in time and space proportionally to the modeled time-dependent changes in the CNIH-2 distribution (Fig. 5A, Eq. S14 in Supplementary Information). With these parameters, our model reproduced the experimental results reported by Clavet-Fournier and colleagues^62^, showing a gradual increase in GluA2 subunits in PSDs that reached 150% 2 hours after cLTP induction and remained elevated after 5 hours (Fig. 5D, G). Does *de novo* CNIH-2 synthesis need to occur in dendrites to obtain a long-lasting increase in GluA2-containing AMPAR concentration in PSDs? To determine whether dendritic CNIH-2 synthesis was necessary for the sustained GluA2 increase, we restricted increased CNIH-2 synthesis to the somatic region in our next simulation and used changes in CNIH-2 dendritic concentration as the change in exocytosis rate. Additionally, we also increased the incorporation rate by a factor of 1.7. Strikingly, our model predicted a 25% increase in GluA2 in the PSDs of stimulated spines after 5 hours, far below the experimentally reported concentration^62^ (Fig. 5E, G). This 25% increase was due to the increased synaptic incorporation. These results highlight the critical role of sustained local protein synthesis, specifically of CNIH-2, in controlling the kinetics of GluA2-containing receptors in PSDs during long-term potentiation. Finally, we tested whether a brief 10-minute CNIH-2 local synthesis increase could recapitulate experimental results (Fig. 5F, G). This brief upregulation failed to match the amplitude of changes in synaptic GluA2. These findings suggest local synthesis of CNIH-2 as the mechanism behind activity-dependent GluA2 changes and highlight the importance of the long-lasting increase in CNIH-2 local synthesis and a concomitant increase in the receptor incorporation rate. We additionally evaluated an alternative LTP induction paradigm in our model, in which CNIH-2 local synthesis was upregulated across all spines for either 2 hours or 10 minutes (Fig. S16A-B). In both conditions, the model predictions failed to recapitulate the experimental observations reported by^62^ (Fig. S16C-D). This discrepancy can be attributed to distinct mechanistic limitations in each case. During prolonged (2-hour) upregulation of CNIH-2 synthesis, the intracellular GluA2 pool was rapidly depleted, resulting in an insufficient reservoir for receptor recycling (Fig. S16E, middle panel). By contrast, brief (10-minute) upregulation of CNIH-2 synthesis produced only a modest increase in CNIH-2 levels, which was insufficient to drive the level of exocytosis required to support synaptic plasticity (Figure S16F).

Based on our experimental measurements and mathematical modeling, we replicated the fast and transient synaptic incorporation of GluA1 homomers as a result of fast and transient changes in their exocytosis coupled with a modest rise in synaptic receptor incorporation rate. We also showed that the slow but persistent synaptic changes in GluA2-containing AMPARs can be explained by a long-lasting up-regulation of the local synthesis of its auxiliary subunit CNIH-2 (2 hours), again coupled with a modest rise in the receptor synaptic incorporation rate (*η*). Thus, our integrative approach suggests a novel model of AMPAR delivery to synapses upon chemical LTP induction, where GluA1-homomers are exocytosed immediately after LTP induction while GluA2-containing AMPARs exocytosis slowly picks up relying on CNIH-2-local translation and lasts for hours (Fig. 6).

**Figure 6:**
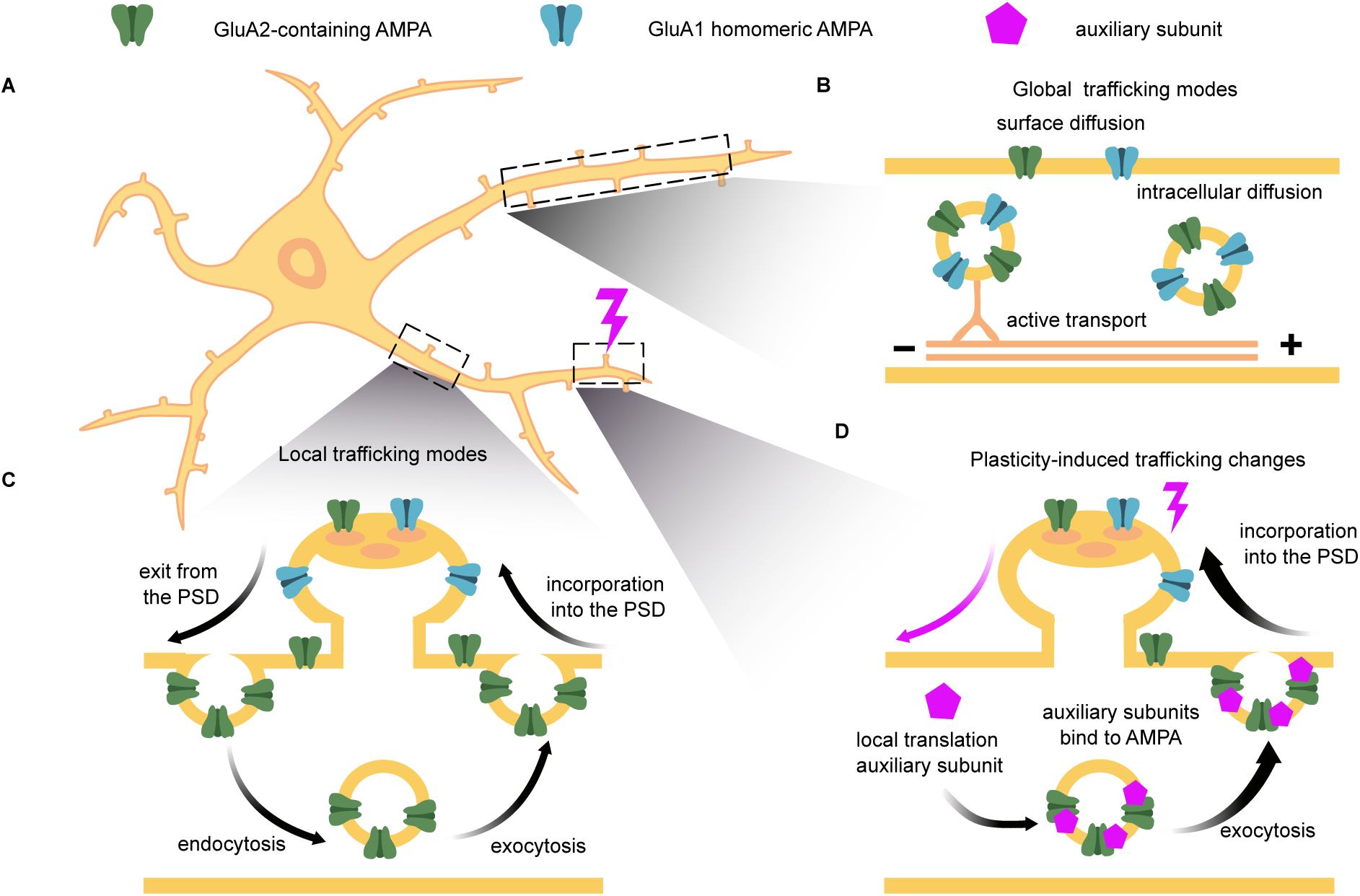
Proposed model of trafficking of AMPARs subtypes upon LTP induction. **(A)** Synapses distributed along the dendritic arbors require GluA1and GluA2-containing AMPARs for LTP. The time scales of different AMPAR trafficking mechanisms range from seconds to hours. **(B)** On the global scale of a neuron, both AMPARs move via fast bidirectional active transport intracellularly and slow passive diffusion on the cell surface or inside the cytoplasm. **(C)** The local trafficking modes, including slow endocytosis and exocytosis of AMPARs, help to maintain a sufficient pool of extrasynaptic receptors. Under the base-line parameterization used here, the model assigns a larger surface-to-intracellular receptor ratio to GluA1-homomeric AMPARs than to GluA2-containing AMPARs, corresponding to a larger intracellular fraction for the latter. The synaptic concentration of AMPARs is further tuned by their diffusion towards the synapse and binding to PDZ domain scaffold proteins at the PSD. **(D)** Our results indicate a novel mechanism specific to GluA2-containing AMPARs that requires local translation of an auxiliary subunit, CNIH-2, that can in turn modify the exocytosis rate of the AMPARs, bringing them closer to synaptic sites.

## Discussion

Our study employed an integrative approach combining experiments and computational modeling to unravel the role of the fast and slow kinetics in global and local trafficking of AMPARs. This approach provides novel insights into the molecular mechanisms underlying the long-recognized but poorly understood temporal differences in synaptic insertion dynamics of the two primary AMPAR subtypes - GluA1 homomers and GluA2-containing heteromers. Notably, we present the first evidence of how the local dendritic translation of the AMPAR auxiliary subunit CNIH-2 regulates the dynamics of GluA2-containing receptors at synapses, unveiling novel perspectives on synaptic function and plasticity (Fig 6).

Synthesis of AMPAR subunits from the corresponding mRNA can occur in various neuronal compartments. Previous studies^3, 70^ have identified *Gria1* and *Gria2* mRNAs in dendrites, suggesting that local protein synthesis can provide a plausible mechanism for rapid protein availability in dendrites. Additionally, previous studies found that the translation rate of AMPARs is location-specific and can differ under basal conditions and during plasticity. In fact,^31, 70, 71^ reported the dynamic regulation of AMPAR mRNA subunits in synapses. However, these prior analyses have predominantly focused on proximal parts of dendritic arbors near the cell somata. In our earlier work, we also observed the dynamic regulation of the newly synthesized GluA1 subunits in dendrites during homeostatic scaling^17^. These experiments quantified the total newly synthesized GluA1 subunits after 3 hours of metabolic labeling using AHA^69^. However, the rapid processes of diffusion and intracellular trafficking made it challenging to distinguish whether the newly synthesized subunits originated from the soma or the dendrites^5^.

In this study, we quantified the relative distribution of AMPAR subunits mRNA in the somatic and dendritic compartments of neurons under basal conditions. Our results revealed that mRNA coding for the pore-forming subunits GluA1 and GluA2 were predominantly somata-enriched in pyramidal neurons (Fig. 1A-C and Fig. S1). Our results are in line with those of Glock and colleagues^4^, who investigated the translatome of neurons by comparing the soma layer and the neuropile (containing mainly dendrites, axons, and glia) from the CA1 region of the hippocampus. The RNA footprint data show very small coverage for *Gria1* and *Gria2* in the neuropil but a large coverage in the soma layer^4^. Despite the mRNA concentration rapidly decreasing with the distance from the soma, the GluA2 subunit exhibits an almost constant distribution along the dendrites (Fig. 1E). Interestingly, in a recent study, authors showed that ribosomes positioned near AMPA receptor complexes at synapses support local translation of postsynaptic proteins, indicating that dendrites can synthesize PSD components and auxiliary subunits such as CNIH-2 despite predominantly somatic AMPAR subunit production^72^. Using our model, we demonstrate that these seemingly contradictory observations can be explained by a biased anterograde active transport of GluA2-containing AMPARs. Specifically, our results indicate that only biased active transport can lead to AMPAR density increase with the distance from the soma - a phenomenon reported in prior AMPAR studies as distance-dependent scaling of AMPARs^32, 33^. We note that the increase of 180% at 500 *µm*, though consistent with previous literature, should be interpreted qualitatively. For example, in the HaloTag-GluA2 data we observed only a modest increase of 15% (Fig. S13,^63^). Hence, the precise increase in AMPAR density requires a more careful characterization of active transport. We also explored the impact of other mechanisms, including passive diffusion rates and protein half-life. However, we found that these factors could not account for the observed distance-dependent scaling of AMPARs. This distance-dependent increase has been reported for other ion channels, such as HCN channels and Kv4.2 potassium channels^73, 74^.

While the increase in concentration of Kv4.2 along the dendrite is attributed to its active transport, no evidence of active transport was reported for HCN channels. We hypothesize that biased active transport can be a common mechanism for ion channels and other transmembrane proteins to achieve a distance-dependent concentration increase along dendritic arbors.

Previous experimental studies have shown that endocytosis and exocytosis of AMPARs take place throughout the whole dendritic tree^54, 56^. However, the precise balance between these two processes remains poorly characterized. We used sequential labeling with cell-impermeable and cell-permeable HaloTag ligands to visualize surface and intracellular GluA2-containing AMPARs independently (Fig. 2). By defining the surface-to-intracellular GluA2 fluorescent intensity ratio, we quantified the net balance between exocytosis and endocytosis rates in our model. Our computational model demonstrates that, under basal conditions, the rate of exocytosis is slightly higher than that of endocytosis, leading to a smaller pool of intracellular GluA2-containing AMPARs. Quantitatively, we estimate that approximately 44% of these receptors are located intracellularly, consistent with the previous experimentally reported range (30% - 70%)^75–77^. Furthermore, we found that this balance between exocytosis and endocytosis is consistently maintained throughout the dendritic tree (Fig. S5). We note that AMPAR can exist in several functional pools intracellularly, including a rapid recycling pool in which receptors are endocytosed and reinserted via recycling endosomes on timescales of minutes^78^. In addition, AMPARs also populate slower intracellular reservoirs, including late endosomal and degradative compartments, which can be mobilized under specific conditions^79^. These pools are not strictly segregated but dynamically exchange, and are often described in terms of fast recycling versus more stable intracellular reservoirs rather than discrete vesicle populations. In the present model, these intracellular compartments are not explicitly distinguished and are instead captured through effective trafficking rates.

Additionally, image analysis revealed higher surface GluA2 levels in PSD95-defined spine-associated ROIs than in neighbouring dendritic shaft ROIs. However, the spatial resolution of confocal imaging does not allow us to determine whether these receptors are localized within the PSD or on the membrane surrounding the PSD inside the spine. Plasticity responses of AMPARs can differ depending on their subunit compositions^1^. GluA1 homomers exhibit fast surface insertion and accumulation in PSDs^57, 66^, whereas GluA2-containing, heteromeric AMPARs exhibit slower synaptic accumulation^62, 66^. Using our model, we investigated the temporal dynamics of synaptic AMPAR density changes for these two receptor subtypes (Fig. 3). For GluA1 homomers, an immediate increase in exocytosis combined with a modest rise in the synaptic incorporation rate was sufficient to replicate the observed rapid and transient accumulation. These results are encouraging as our model could replicate the GluA1-homomeric dynamics without an explicit fitting to the experimental data. vIn contrast, GluA2-containing heteromers required a time-dependent change in the exocytosis rate to match the experimentally reported concentration, while a short-term increase alone was insufficient. These distinct timescales of AMPAR subtypes may contribute to the temporal organization of synaptic plasticity at the circuit level, with rapid changes in synaptic strength followed by slower, more sustained modifications of network connectivity. By modulating the ratio of these receptors, a neuron may gate between the fast and slow plasticity at the circuit level. We propose that this timedependent exocytosis may be driven by local changes in the CNIH-2 concentration - a protein known to regulate AMPAR trafficking and synaptic accumulation. However, our model reproduced the AMPAR plasticity response only when the stimulation was restricted to a small fraction of spines ( 2%), substantially lower than observed under chemical LTP ( 40% structural potentiation, see^82^). This discrepancy arises from strong resource constraints in the model, where intracellular receptor availability is rapidly depleted, limiting AMPAR insertion to a small subset of spines. Consequently, the model captures a resource-limited, high-threshold regime of plasticity and underestimates the broader synaptic recruitment seen under cLTP.

Interestingly, *Cnih2* mRNA is highly abundant throughout the entire length of neuronal dendrites (Fig. 4A-C), and a previous study using ribosome profiling confirmed that this dendritic mRNA undergoes local translation^4^. Using metabolic labeling with AHA for 1 hour, we observed that most newly synthesized CNIH-2 proteins localized to dendrites with their synthesis increasing after cLTP induction (Fig. 4D-F).

To explore the role of CNIH-2 in AMPAR trafficking, we used shRNAs targeting *Cnih2* mRNA. Unexpectedly, this led to a specific decrease in GluA2-containing receptors at the plasma membrane (Fig. 4G-I), confirming that CNIH-2 synthesis is crucial for GluA2-containing AMPAR surface trafficking. Our computational model further showed that a local up-regulation of CNIH-2 synthesis could explain the amplitude and temporal dynamics of GluA2-containing AMPARs (Fig. 5). In contrast, increasing the somatic influx of CNIH-2 resulted in only minimal effects on AMPAR concentration at the cell surface and in synapses, highlighting the critical role of the local CNIH-2 synthesis in regulating AMPAR trafficking under activity-dependent conditions. The molecular mechanisms driving the increase in local CNIH-2 synthesis upon LTP induction remain an active area of research. One key molecule implicated in memory maintenance via regulation of GluA2-containing AMPARs trafficking is protein kinase M*ζ* (PKM*ζ*)^83^. Notably, experimental studies have shown that PKM*ζ* controls dendritic protein synthesis through phosphorylation-mediated inhibition of Pin1 (Peptidyl-prolyl cis-trans isomerase NIMA-interacting 1), a suppressor of protein synthesis^84^. Future experimental work can explore whether PKM*ζ* regulates GluA2-containing AMPARs trafficking by modulating dendritic translation of CNIH-2, providing new insights into molecular mechanisms of memory maintenance.

The model introduced in this study provides a foundation that can be extended in multiple directions in future work. First, AMPAR subtype trafficking is multi-factorial and can be regulated by additional mechanisms such as phosphorylation of the GluA1 subunit at serine residues 831 and 845^1^, dynamic binding with scaffolding proteins including GRIP and PICK1^85, 86^, and interaction with other auxiliary subunits such as TARPs^9^. While our findings highlight the key role of CNIH-2, these other pathways likely act in parallel to shape the overall trafficking profile. Future studies should examine how CNIH-2 regulation integrates with other modulatory mechanisms to drive AMPAR subtype-specific trafficking. Importantly, Schwenk and colleagues^12^ have demonstrated a physical association between CNIH proteins and AMPAR subunits in native complexes, suggesting a possibility of such a coupling. An important next step would be to confirm how CNIH-2 couples to the GluA2-containing transport vesicles using co-transport assays and live imaging. Second, we considered our model dendrite to be a reduced, one-dimensional cylinder. Future work can extend this framework and incorporate explicit branching geometry. A prior study^39^ showed that membrane-anchored proteins are 35% more likely to pass branch points and reach distal dendritic sites than cytosolic proteins. For AMPARs, however, continuous endo- and exocytosis tends to re-balance this surface bias, and crucially, our analysis indicates that active transport dominates passive diffusion in supplying receptors to distal sites. Furthermore, because microtubule number is approximately conserved across branchpoints^87, 88^, our one-dimensional reduction provides a plausible, morphology-averaged baseline for detailed neuronal arbors. Finally, our study focused on chemically induced LTP, a widely used and well-established protocol for eliciting robust, long-lasting potentiation and for constraining intracellular signalling models. Nonetheless, integrating physiologically patterned activity protocols will be an important direction for future work to fully assess model generalisability.

Overall, our findings elucidate alternative routes for AMPAR delivery to synapses (Fig. 6). Previous work has identified unconventional secretory processing of neuronal surface proteins, including AMPARs with Endo H-sensitive glycans. In contrast, GluA2-containing heteromers, synthesized at the surface of the somatic ER, may bypass GA membranes and reach the neuronal plasma membrane directly in dendrites. This process appears to be mediated by the local synthesis of CNIH-2, resulting in AMPARs with Endonuclease H-resistant glycans, as shown in previous work^89^. While a complete understanding of the molecular interplay between AMPAR auxiliary subunits remains to be uncovered, our findings provide insights into how distinct AMPAR populations are selectively recruited to synapses under different conditions. This selective recruitment may have profound implications for therapeutic strategies aimed at restoring or modulating AMPAR synaptic transmission and plasticity - processes that are often disrupted in neurological and neurodegenerative disorders.

## Methods

### Ethics statement

All animal experiments were conducted in accordance with applicable national and institutional guide-lines. Experiments conducted in Germany complied with German national animal care guidelines and those of the Max Planck Society or Goethe University Frankfurt and are in compliance with international laws and policies (DIRECTIVE 2010/63/EU; German animal welfare law; FELASA guidelines). The animals were euthanized according to annex 2 of § 2 Abs. 2 Tierschutz-Versuchstier-Verordnung and animal numbers were reported to the local authorities (Regierungspräsidium Darmstadt). Experiments conducted in the Netherlands complied with Dutch national animal care guidelines and were approved by the Centrale Commissie Dierproeven (CCD) and the local animal welfare body under animal experiment project 2021-0040. Experiments conducted in France complied with European guidelines for the care and use of laboratory animals and the guidelines of the University of Bordeaux animal experimentation committee (CE50; animal facility authorisations A3306940 and A33063941).

### Experimental protocols

#### Cultured neurons

Dissociated rat hippocampal cultures were prepared and maintained as described previously^97^. Briefly, we dissected hippocampi from postnatal day 0-1 rat pups of either sex (Sprague-Dawley strain; Charles River Laboratories), dissociated them with papain (Sigma) and plated them at a density of 40 10^3^ cells/*cm*^2^ on poly-D-lysine coated glass-bottom Petri dishes (MatTek). For HaloTag experiments, we dissected hippocampi from E18 rat embryos of either sex (Sprague-Dawley strain; Janvier), dissociated using trypsin (Gibco) and plated at a density of 30 10^3^ cells/*cm*^2^ on poly-D-lysine and laminin coated 18-mm glass coverslips (VWR).

Neurons were maintained and matured in a humidified atmosphere at 37°C and 5% CO_2_ in growth medium (Neurobasal-A supplemented with B27 and GlutaMAX-I, life technologies) for 18-21 days in vitro (DIV) to ensure synapse maturation. For HaloTag experiments, medium was refreshed every 7 days with BrainPhys neuronal medium (StemCell technologies) with SM1 (StemCell technologies) and 1x Pen/Strep (Gibco). All experiments complied with national animal care guidelines and the guidelines issued by either the Max Planck Society or the Dutch authorities (CCD), and were approved by local authorities. For transfection, DIV7-11 neurons were transfected using Effectene (Qiagen). Transfected cells were maintained until DIV19 for experiments. For HaloTag experiments, DIV7 neurons were transfected using lipofectamine 2000 (ThermoFisher) with the Halo-GluA2 donor plasmid and the Halo-GluA2 gRNA plus spCas9. Per coverslip, 3.3 *µ*l lipofectamine, 1000 ng of gRNA+Cas9 plasmid and 500 ng of donor plasmid were diluted in 200 *µ*l BrainPhys. Neurons were incubated in 200 *µ*l conditioned medium plus 200*µ*l transfection mix for 1.5-2h at 37°C and 5% CO_2_. After incubation, coverslips were transferred back in conditioned medium plus half a volume of fresh BrainPhys medium.

#### Cloning of plasmids

To replace the SEP tag of the TKIT SEP-GluA2 knock-in (a generous gift from Richard Huganir, Addgene #169440,^51^), we first amplified the HaloTag from the Halo-Homer1 knock-in plasmid (a generous gift from Harold MacGillavry, Addgene #183446,^98^) using PCR with primers 5’-gcAGCagaGCAGAAATCGGTACTGGCTTTCCATTCG-3’ and 5’-aGccTccCccTccaagGCCGGAAATCTCGAGCGTgGAC-3’ and amplified the SEP-GluA2 backbone without the SEP tag using PCR with primers 5’-TCGAGATTTCCGGCcttggAggGggAggCtcAggcggaggtgggtcgggtg-3’ and 5’-GAAAGCCAGTACCGATTTCTGCtctGCTgccaccgccaccactagatatgc-3’. Subsequently, we performed Gibson assembly using a vector:insert ratio of 1:5. Competent cells were transformed and grown, individual colonies were picked and plasmid DNA was isolated. Correct clones were identified using whole-plasmid sequencing (Plasmidsaurus).

#### HaloTag staining, immunostaining, image acquisition and analysis

Coverslips with cultured hippocampal neurons (DIV 21) were incubated for 10 minutes in a 50-*µ*l droplet of conditioned medium with 0.2*µ*M cell-impermeable HaloTag Ligands (JF635i or JF549i; Promega) at 37°C and 5% CO_2_.

Then, cell-permeable HaloTag Ligands (JF635 or JF549; Promega) were added at a concentration of 0.2*µ*M and neurons were again incubated for 10 minutes at 37°C and 5% CO_2_. After placing the coverslips back in conditioned medium, they were washed and fixed once at room temperature in DPBS containing calcium and magnesium before being fixed in 4% paraformaldehyde in PBS (pH = 7.4) for 10 minutes on ice. Subsequently, neurons were washed with DPBS 3 times, blocked in blocking buffer (4% NGS/PBS with 0.1% Triton X-100) for 30 min at 37°C, and incubated with mouse anti-PSD95 antibody (1:1000, ThermoFisher) and chicken anti-MAP2 antibody (1:1000, Synaptic Systems) in blocking buffer at room temperature for 2 hours. After washing 3 times with DPBS, neurons were incubated with anti-mouse AF488 antibody (1:500, Abcam) and anti-chicken AF405 antibody (1:500, Abcam). Three images per coverslip were acquired on a Nikon SoRa spinning disk microscope fitted with a 40x/1.30 oil objective, 405, 488, 561 and 638 laser lines, and an ORCA-Fusion BT CMOS camera (Hamamatsu). Z stacks spanning the entire volume of neurons were obtained with 0.5 *µ*m z-steps and with the 2.8x SoRa magnifier engaged. Channels were separated and collapsed to a sum intensity projection in ImageJ. Manual annotation of dendritic tree using segmented line tool and somata using polygonal ROIs was performed. Signal intensity profile in dendritic ROIs was measured using a constant width and integrated density was calculated for somatic ROIs.

#### In situ hybridization in cultured neurons

In situ hybridization was performed using QuantiGene ViewRNA kit from Affymetrix (now Thermo Fisher Scientific) mostly following the provider instructions. In brief, cells (DIV 18-24) were fixed for 20 min at room temperature using a 4% paraformaldehyde (PFA) solution (4% paraformaldehyde, 2.5% Sucrose, in lysine-phosphate buffer). The Proteinase K treatment was omitted in order to preserve the integrity of the dendrites. After permeabilization using the provider’s detergent buffer for 2 min, cells were directly incubated for 3h at 40°C with detection probes. Incubations with pre-amplification, amplification and detection probes were reduced to 40 min each. After completion of in situ hybridization, cells were washed with PBS and incubated in blocking buffer (4% goat serum in PBS) for 1hr. Neurons were subsequently processed for immunofluorescence using standard methods as described below.

#### Immunofluorescence in cultured neurons

All steps were performed at room temperature, unless stated otherwise. Glass bottom dishes with attached neurons (DIV 18-21) were fixed in paraformaldehyde 4% in lysine phosphate buffer pH7.4 containing 2.5% sucrose for 15-20 min. For simple immunofluorescence, neurons were permeabilized for 10 min in PBS + 0.5% Triton-X 100. After 2 quick washes in PBS, neurons were then incubated in blocking buffer (4% goat serum in PBS) for 30 min. Then neurons were incubated for 1 hr or overnight with primary antibodies, and after 3 washes of 5 min, they were incubated for 1.5 hr with secondary antibodies. We used the following antibodies: guinea pig anti-MAP2 (Synaptic Systems, 1:2000), mouse anti-biotin (Sigma, 1:1000), rabbit anti-biotin (Cell Signaling, 1:1000), mouse anti-Puromycin (Kerafast, 1:2500), homemade rabbit anti-Puromycin (1:250), homemade anti-GluA1 N-terminal domain (targeted sequence QWRTSDSRDHTRVDWKRPKC; KO validated by western blot) (fixed sample 1/500; live-labeling 1/200), mouse anti-GluA2 N-terminal domain from Eric Gouaux previously used in Nair et al., 2013 (fixed sample 1/100; live-labeling 1/500), rabbit anti-CNIH-2 (Synaptic Systems, 1/200).

#### Chemical long-term potentiation (cLTP)

For cLTP experiments, the original medium of cultured neurons was replaced for 5 min by artificial cerebrospinal fluid (ACSF) at 37°C. In control condition, the ACSF contained Ca2+ (2 mM) and Mg2+ (2 mM) and supplemented with B27 and MEM Amino Acids (50X) (Thermo Fisher Scientific). In the cLTP-induction condition, the ACSF differed from the control-ACSF as follows: Ca2+ (3 mM), Mg2+ (0 mM), glycine (200 *µ*M), picrotoxin (100 *µ*M). After the 5 min induction the cells were placed in growth medium containing either methionine (Met) or L-Azidohomoalanine (AHA) at 4 mM for 1 hr.

#### Fluorescence non-canonical amino acid tagging

To investigate the distribution of newly synthesized proteins we used a recently developed method combining fluorescence non-canonical amino acid tagging with the proximity ligation assay (FUNCAT-PLA)^69^. Cultured neurons were incubated for 1 hr with growth medium in which Met was replaced by AHA (4 mM). In control conditions the growth medium contained Met (4 mM) and no AHA.

#### Puromycylation

To determine the sites of synthesis of AMPAR components we used a recently developed method combining puromycylation and proximity ligation assay (Puro-PLA)^69^. Cultured neurons were labeled with 10 *µ*M puromycin (Sigma-Aldrich) for 2 min. In control experiments cells were treated with 40 *µ*M of the protein synthesis inhibitor anisomycin (Sigma-Aldrich) for 30 min prior to and during puromycin labeling.

#### Proximity ligation assay

Detection of newly synthesized proteins by proximity ligation was carried out using anti-puromycin or anti-biotin antibodies (in case of AHA metabolic labeling) in combination with protein-specific antibodies against our proteins of interest (tom Dieck et al., 2015)(complete list of antibodies in the Immunofluorescence section). We used Duolink reagents (Sigma) and followed the protocol provided by the manufacturer with some modifications described below. We routinely used rabbit PLAplus and mouse PLAminus probes amplification and label probe binding. Briefly, after washing the primary antibodies 3 times in PBS, PLA probes were applied in 1:10 dilution in PBS with 4% goat serum for 1 h at 37 °C, washed several times with wash buffer A (0.01 M Tris, 0.15 M NaCl, 0.05% Tween 20) and incubated for 30 min with the ligation reaction containing the circularization oligos and T4 ligase prepared according to the manufacturer’s recommendations (Duolink Detection reagents Red, Sigma) in a prewarmed humidified chamber at 37°C. Amplification and label probe binding was performed after further washes with wash buffer A with the amplification reaction mixture containing Phi29 polymerase and the fluorophore-labeled detection oligo prepared according to the manufacturer’s recommendations (Duolink Detection reagents Red, Sigma) in a prewarmed humidified chamber at 37 °C for 100 min. Amplification was stopped by three washes in wash buffer B (0.2 M Tris, 0.1 M NaCl, pH 7.5). For better signal stability, cells were kept in wash buffer B at 4°C until imaging.

#### Image acquisition

Cultured cells were imaged using Zeiss LSM780/880 confocal microscopes using a 63x oil objective NA 1.4 and a 40x oil objective NA 1.3. Z-stacks spanning the entire volume of imaged neurons were obtained.

#### Puncta detection

Cell culture images were analyzed to assess mRNA and newly synthesized protein distributions in somata and dendrites using a custom built Python tool called SpyDen^96^. The code and instruction manual for SpyDen is freely-available for download online https://github.com/meggl23/SpyDen.

#### Statistical analysis

All data are shown as the mean s.e.m. unless stated otherwise. Box plots show the median, quartile and whisker using lines and mean using triangle. Swarm plot shows the individual values. We performed all the data analysis using Python programming language. We did not assume that the data were normally distributed, hence we used 2-sided Mann-Whitney (rank-sum) test for independent sample comparisons and 2-sided Wilcoxon signed-rank test for pairwise comparisons, unless stated otherwise.

### Model for the combined distribution of surface and cytoplasmic proteins in dendrites (related to section Active transport of AMPARs can explain their homogeneous distribution along the dendrite and Fig. 1)

Under basal conditions, AMPARs are primarily produced in the soma and move along the dendrites via active intracellular transport as well as diffusion within both the plasma membrane and intracellular compartments. The following boundary value problem describes the dynamics of a protein (*P*) along the dendrites shaped by the diffusion (*D_P_*), net active transport velocity (*V_P_*) and degradation (*λ_P_*)

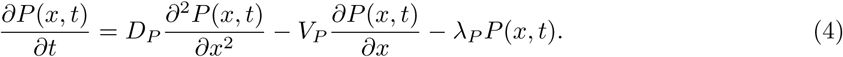

We assume that the concentration of the protein (*P*) satisfies the boundary conditions

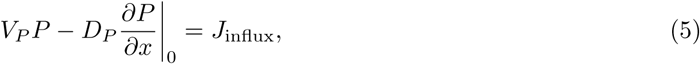

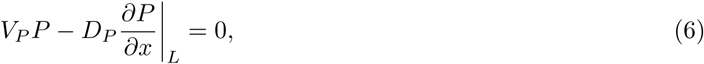

whereby *J*_influx_ *>* 0 is the rate of protein influx in the dendrite from the soma and Eq (6) means that no proteins come out of or enter at the dendritic tip located at the distance *L* from the soma.

The steady state solution of the Eq (4) (i.e., solution under the assumption 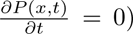 is a sum of two exponential functions

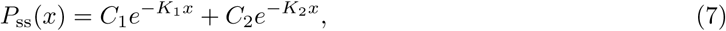

whereby 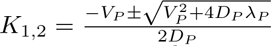. We compute the constants *C, C* using the boundary conditions in Eqs (5), (6) and obtain the steady state dendritic distribution of the protein

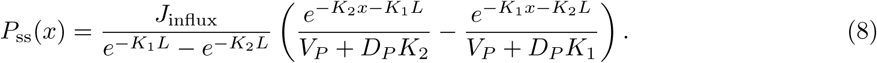

### Model for GluA2-containing AMPAR concentration in surface, cytoplasm and PSD (related to section Endogenous concentrations of AMPARs in the cytoplasm, cell membrane, and postsynaptic density and Fig 2)

To compute the steady-state solutions of the model in Eq (1) - (3), we assume that the time derivatives of the functions *P_s_*, *P_c_* and *P*_psd_ are equal to 0 and obtain the following system of ODE

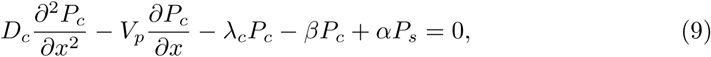

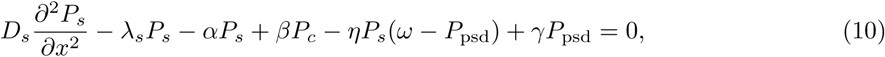

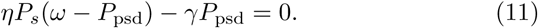

The relation in Eq (11) implies that steady state of the synaptic proteins (*P*_psd_) is related to the steady state concentration of the proteins in the cell membrane via

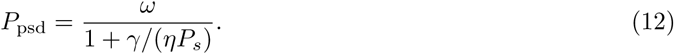

Combining Eqs, (10) and (11), we reduce the system in Eqs 9-11 to two equations

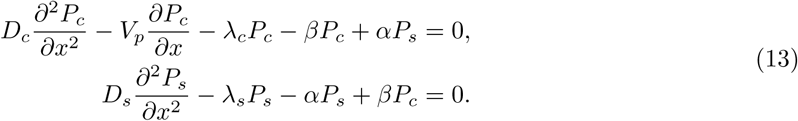

We assume that the dendrite has the length *L* and the protein molecules can not escape the dendrite at the point *x* = *L*, i.e., the functions *P_c_* and *P_s_* fulfill the following boundary conditions

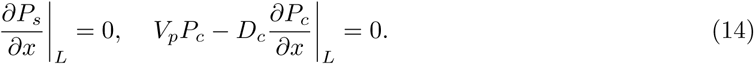

Also, we assume that the proteins *P_c_* produced in the soma are released in the cytoplasm at *x* = 0 with a constant influx rate *J*_cin_ and reach the plasma membrane *P_s_* only via exocytosis. Hence we assume no-flux from the soma into the dendrite, i.e.,

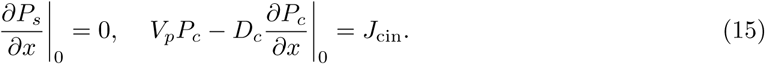

We solve Eqs (13) numerically using Python library *solve_bvp* which computes the numeric solution of the ODE system subject to two-point boundary conditions.

### Fitting model to the data

#### Fitting mRNA distributions

To estimate the decrease in *Gria1*, *Gria2*, *Cnih2*, and *Camk2a* mRNA distributions along a dendritic branch, we binned the empirical data (bin size = 7.5 *µ*m) and fitted an exponential function of the form *f* (*x*) = *A e^−Bx^* to the binned data. This function describes the distribution of mRNA along the dendritic branch, where the independent variable *x* denotes the position along the branch. The parameters *A >* 0 and *B >* 0 were fitted using the *minimize* function from the *lmfit* Python package to minimize the sum of squared residuals between the data and the model.

#### Fitting *D_P_* and *V_P_* for total GluA2 model

To estimate the global transport parameters for the model, we numerically minimized the error between normalized distribution of the first 100 *µ*m dendritic distribution from the model and experimental data over the two-dimensional parameter space to fit the experimental data. For both, experimental data and model, a binning (bin size = 5 *µ*m) and normalization by the first bin value was done before the optimization. The fitting was done using the *minimize* function of *lmfit* python package for nonlinear least square minimization between the data and the model. We note that in all of our simulations, we assumed the length of the model dendrite to be *L* = 500 *µm* and then performed the fitting.

#### Calculating surface to intracellular intensity ratio

To quantify total surface and intracellular GluA2 levels, z-stack images were sum-projected, and the integrated density was calculated from manually defined somatic and dendritic regions of interest (ROIs). Dendritic ROIs were defined as segmented lines of constant width, while somatic ROIs were defined as polygons.

Let *S* and *I* denote the measured surface and intracellular AMPAR intensities, respectively, within a dendritic segment (or somatic region) of area *A*. Background-subtracted intensities were computed as:

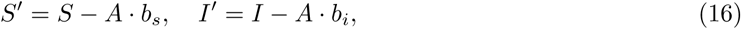

where *b_s_*and *b_i_* represent the mean background intensity values for the surface and intracellular GluA2 channels, respectively.

#### Calibration of HaloTag fluorescence using dye switching

To account for differences in fluorophore brightness, we performed a dye-switching calibration. First, we computed the ratio of surface to intracellular GluA2 intensity using JF635i (surface) and JF549 (intracellular), denoted as *R*_1_. We then repeated the measurement using the inverse labeling configuration — JF549i (surface) and JF635 (intracellular) — to obtain *R*_2_.

To mitigate bias arising from differences in fluorophore intensity, the final ratio was defined as the geometric mean of *R*_1_ and *R*_2_:

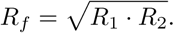

To estimate the mean and variability of *R_f_*, we performed bootstrap resampling by drawing 1000 subsets from the original ratio measurements.

#### Spine-head and dendritic shaft segmentation

Spine and dendritic regions of interest (ROIs) were derived from a supervised segmentation of the PSD95-intrabody channel, using Labkit as a FIJI plugin^99^. The resulting segmentation was transformed to binary masks corresponding to PSD95 spineassociated-positive structures and PSD95 spine-associated-negative dendritic compartments, based on user-defined class ranges. PSD95 spine-associated ROIs masks were further processed by watershed segmentation to separate adjacent puncta.

Analyses were performed within manually annotated dendritic branch ROIs, which were spatially expanded to include the PSD95 spine-associated ROIs masks. Candidate PSD95 spine-associated ROIs were identified as PSD95 spine-associated-positive structures that satisfied predefined size and circularity criteria. They were filtered based on their fractional overlap with the original branch ROI so that only objects below a specified overlap value were retained.

Finally, spine ROIs (and their corresponding dendritic ROIs) with mean surface GluA2 (sGluA2) intensity below background levels were excluded from further analysis, ensuring that only spines exhibiting detectable surface GluA2 were retained. (See Fig. S15, for an example of “empty/non-specific” spine that was filtered out during this criterion).

For each accepted spine ROI, a corresponding dendritic ROI was defined by iteratively expanding a circular region centered on the spine until it intersected the dendritic mask. The intersecting dendritic signal within this local neighborhood was then converted into a dendritic ROI, yielding paired spine– dendrite regions for subsequent quantification. All measurements were performed on a per-pair basis across imaging channels. The measured values were then background-subtracted, followed by PSD95 spine-associated ROIs/dendrite ratio calculation which was finally averaged within individual cells as Synaptic Enrichment (SE).

#### Exocytosis rate for the extended GluA2 model

To calculate net exocytosis rate at a steady state we computed the ratio of surface GluA2 fluorescence 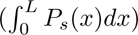 to the intracellular GluA2 fluorescence 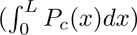 as described above. The following proof shows that in our model this ratio is indeed equal to the ratio between the exocytosis and endocytosis rates (*β/α*). Integrating Eqs. (13) from 0 to *L*, we obtain

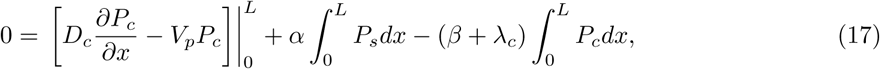

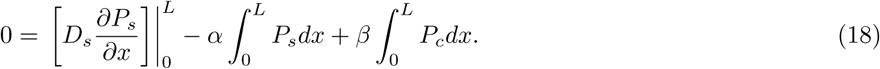

Using the boundary conditions from Eqs. (14 - 15), we obtain

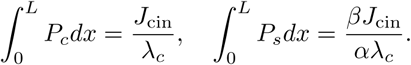

Hence, we compute the ratio between the exocytosis and endocytosis rates as

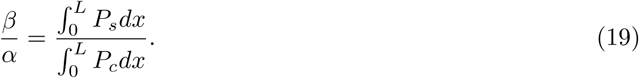

We used the endocytosis rate as described in Table 2 and *λ_s_*= 0. For the GluA1-homomer version of the model, we used a ratio of 1.7 to obtain the exocytosis rate. This ratio was reported in^100^.

#### Fitting *D_s_*, *D_c_* and *V_p_* in the extended model

The exocytosis rate was calculated using the above mentioned method before fitting the global transport parameters, *D_s_*, *D_c_* and *V_p_*. To estimate the parameters we numerically minimized the error between the normalized model-predicted dendritic distribution and the normalized experimental data over the first 100 *µ*m of the dendrite.

#### Fitting *η* in the extended model

All parameters of the extended model were either computed as described in previous sections or set to an experimentally reported value before we estimated the incorporation rate *η* (see, Table 2). To fit *η* in the extended model (Eqs (1)-(3)), we calculated the mean ratio of *P*_psd_ and *P_s_* over the entire length of our model dendrite using *minimize* function to reduce the difference between this ratio and the mean surface synaptic enrichment. We treated the measured surface spine-associated/shaft fluorescence ratio as an approximate phenomenological constraint. Its correspondence to the model’s PSD-to-extrasynaptic receptor ratio is a simplifying assumption; the fitted *η* is conditional on this mapping and the fixed slot number.

#### Baseline model of GluA1-homomeric AMPARs

To generate the steady-state trafficking dynamics of GluA1-homomeric AMPARs, we used the same model as for GluA2-containing AMPARs described in Eqs. (1)-(3). For the GluA1 model we used the parameters in Table. 3, which were either previously reported or estimated in this study. These parameters provide different distributions of the GluA1-homomers and GluA2-containing AMPARs as reported experimentally (see Fig. S14)

## Acknowledgments

We would like to thank all Tchumatchenko, Hafner, Acker-Palmer, Schuman and Choquet lab members for fruitful discussions and helpful feedback.

## Author Contributions

SW performed the simulations and analyzed the simulations and experimental data. SW built the model with theoretical and conceptual input from NK and TT and experimental feedback from EMS, ASH, and TT. The experimental data shown in Figures 1 and 4 were acquired by ASH in EMS’s lab at the Max Planck Institute for Brain Research. The experimental data shown in Figure 2 were acquired by WJD in ASH’s lab. AB performed the shRNA validation experiments S11 in EMS’s lab at the Max Planck Institute for Brain Research under the supervision of ASH and EMS. CNIH-2 protein data Fig. S10 were acquired by ASH at Radboud University. AI performed the endogenous AMPAR staining. DC supervised AI. SW, NK, ASH, and TT wrote the original draft of the manuscript, and all authors edited the manuscript. Data in S13 was analyzed by CB. MKK and AAP provided useful feedback on the modeling work.

## Data availability

The experimental and computational data underlying our study is available at: https://github.com/surbhit21/mRNAAnalysisCodenData, citation^103^, https://github.com/surbhit21/AMPARSimulationsCode, citation ^104^, https://gin.g-node.org/surbhitw/GluAsimulations.git, citation^105^, https://gin.g-node.org/surbhitw/CNIH2simulations.git, citation^106^.

## Code availability

The code to run the CNIH-2 translation and AMPAR simulations, analyze the experimental data, simulated data, and generate the figures in this manuscript is deposited on the following GitHub repositories: https://github.com/surbhit21/mRNAAnalysisCodenData^103^, https://github.com/surbhit21/AMPARSimulationsCode^104^.

## Key resources table (Table 1)

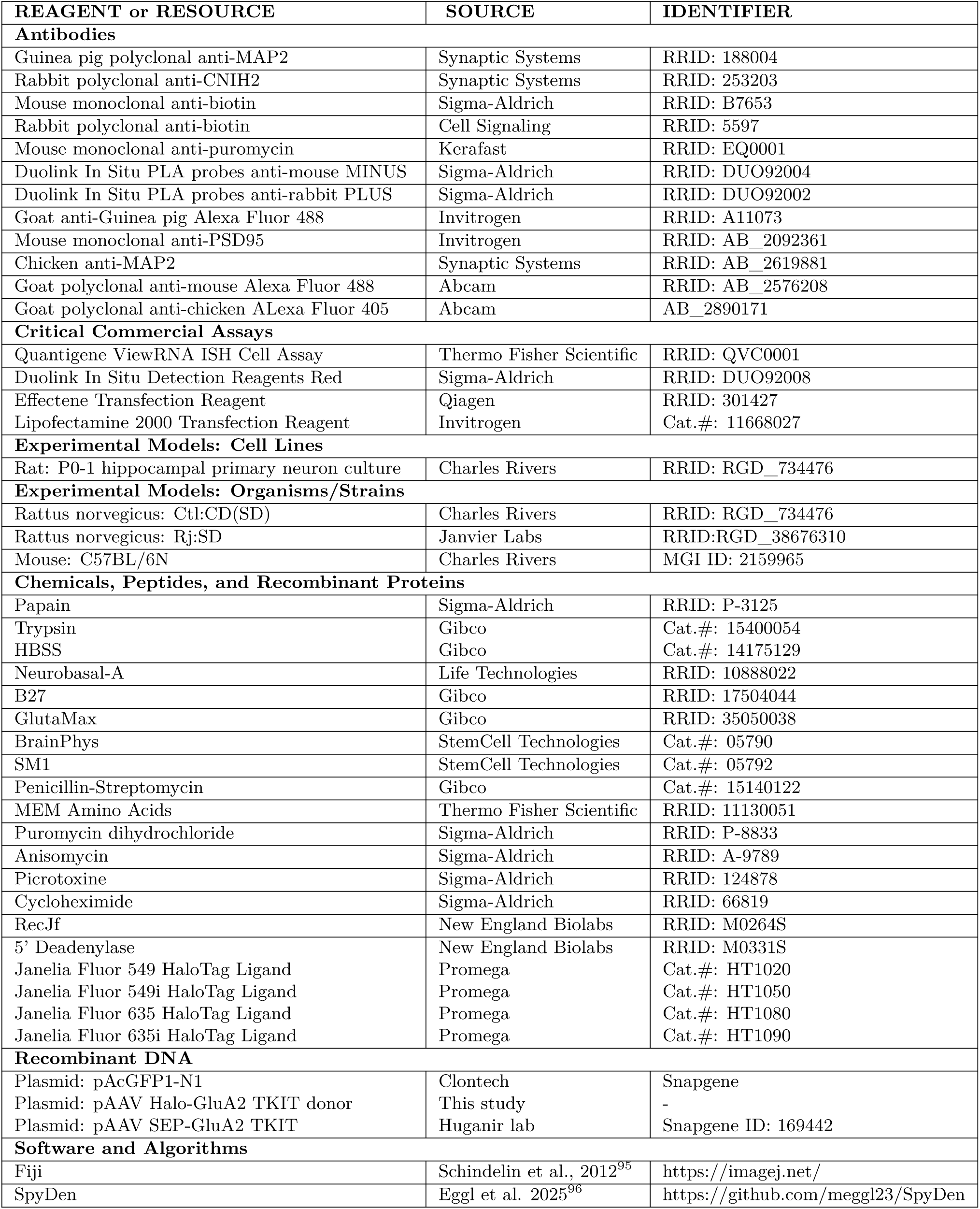

| **REAGENT or RESOURCE** | **SOURCE** | **IDENTIFIER** |
|---|---|---|
| **Antibodies** |  |  |
| Guinea pig polyclonal anti-MAP2 | Synaptic Systems | RRID: 188004 |
| Rabbit polyclonal anti-CNIH2 | Synaptic Systems | RRID: 253203 |
| Mouse monoclonal anti-biotin | Sigma-Aldrich | RRID: B7653 |
| Rabbit polyclonal anti-biotin | Cell Signaling | RRID: 5597 |
| Mouse monoclonal anti-puromycin | Kerafast | RRID: EQ0001 |
| Duolink In Situ PLA probes anti-mouse MINUS | Sigma-Aldrich | RRID: DUO92004 |
| Duolink In Situ PLA probes anti-rabbit PLUS | Sigma-Aldrich | RRID: DUO92002 |
| Goat anti-Guinea pig Alexa Fluor 488 | Invitrogen | RRID: A11073 |
| Mouse monoclonal anti-PSD95 | Invitrogen | RRID: AB_2092361 |
| Chicken anti-MAP2 | Synaptic Systems | RRID: AB_2619881 |
| Goat polyclonal anti-mouse Alexa Fluor 488 | Abcam | RRID: AB_2576208 |
| Goat polyclonal anti-chicken ALexa Fluor 405 | Abcam | AB_2890171 |
| **Critical Commercial Assays** |  |  |
| Quantigene ViewRNA ISH Cell Assay | Thermo Fisher Scientific | RRID: QVC0001 |
| Duolink In Situ Detection Reagents Red | Sigma-Aldrich | RRID: DUO92008 |
| Effectene Transfection Reagent | Qiagen | RRID: 301427 |
| Lipofectamine 2000 Transfection Reagent | Invitrogen | Cat.#: 11668027 |
| **Experimental Models: Cell Lines** |  |  |
| Rat: P0-1 hippocampal primary neuron culture | Charles Rivers | RRID: RGD_734476 |
| **Experimental Models: Organisms/Strains** |  |  |
| Rattus norvegicus: Ctl:CD(SD) | Charles Rivers | RRID: RGD_734476 |
| Rattus norvegicus: Rj:SD | Janvier Labs | RRID:RGD_38676310 |
| Mouse: C57BL/6N | Charles Rivers | MGI ID: 2159965 |
| **Chemicals, Peptides, and Recombinant Proteins** |  |  |
| Papain | Sigma-Aldrich | RRID: P-3125 |
| Trypsin | Gibco | Cat.#: 15400054 |
| HBSS | Gibco | Cat.#: 14175129 |
| Neurobasal-A | Life Technologies | RRID: 10888022 |
| B27 | Gibco | RRID: 17504044 |
| GlutaMax | Gibco | RRID: 35050038 |
| BrainPhys | StemCell Technologies | Cat.#: 05790 |
| SM1 | StemCell Technologies | Cat.#: 05792 |
| Penicillin-Streptomycin | Gibco | Cat.#: 15140122 |
| MEM Amino Acids | Thermo Fisher Scientific | RRID: 11130051 |
| Puromycin dihydrochloride | Sigma-Aldrich | RRID: P-8833 |
| Anisomycin | Sigma-Aldrich | RRID: A-9789 |
| Picrotoxine | Sigma-Aldrich | RRID: 124878 |
| Cycloheximide | Sigma-Aldrich | RRID: 66819 |
| RecJf | New England Biolabs | RRID: M0264S |
| 5’ Deadenylase | New England Biolabs | RRID: M0331S |
| Janelia Fluor 549 HaloTag Ligand | Promega | Cat.#: HT1020 |
| Janelia Fluor 549i HaloTag Ligand | Promega | Cat.#: HT1050 |
| Janelia Fluor 635 HaloTag Ligand | Promega | Cat.#: HT1080 |
| Janelia Fluor 635i HaloTag Ligand | Promega | Cat.#: HT1090 |
| **Recombinant DNA** |  |  |
| Plasmid: pAcGFP1-N1 | Clontech | Snapgene |
| Plasmid: pAAV Halo-GluA2 TKIT donor | This study | - |
| Plasmid: pAAV SEP-GluA2 TKIT | Huganir lab | Snapgene ID: 169442 |
| **Software and Algorithms** |  |  |
| Fiji | Schindelin et al., 2012<sup>95</sup> | https://imagej.net/ |
| SpyDen | Eggl et al. 2025<sup>96</sup> | https://github.com/meggl23/SpyDen |

## Supplemental Information

### GluA2 live labelling by Fabs directly coupled to ATTO643 in hippocampal cultures

#### Primary hippocampal neuron culture

All experiments were performed in accordance with the European guidelines for the care and use of laboratory animals, and the guidelines issued by the University of Bordeaux animal experimental committee (CE50; Animal facilities authorizations A3306940, A33063941). Primary hippocampal cultures were prepared from E18 rat embryos of Sprague-Dawley rats, according to^90^, modified from^107^, at a density of 200,000 cells per 60 mm dish on poly-L-lysine pre-coated coverslips. After 2h the coverslips were flipped in the ‘sandwich configuration’ in dishes containing confluent astrocytes. The cultures were maintained at 37°C and 5% CO2 in Neurobasal PLUS medium (Thermofisher Scientific cat # A3582901) supplemented with 2mM L-glutamine and B27 PLUS Neuronal supplement (Thermofisher Scientific cat # A3582801). After 3 days, cytosine arabinoside “Ara-C” (5 M) was added to inhibit glial cell proliferation.

#### Neuronal transfection

Neurons were transfected using the calcium phosphate method. Briefly, plasmid DNA was diluted in water, followed by addition of 2.5 M CaCl_2_. BBS buffer (50 mM BES, 280 mM NaCl, 1.5 mM Na_2_HPO_4_) was then added, and the mixture was incubated for 15–20 min at room temperature to allow precipitate formation. The mixture was applied dropwise to the neuronal cultures. After 20 min incubation, coverslips were washed twice with HBSS and returned to their original culture dishes.

#### Surface GluA2 labelling

Surface GluA2 subunits were labelled using purified Fab fragments derived from the IgG 15F1 monoclonal antibody (E. Gouaux), directly conjugated to ATTO643 at a degree of labelling of 0.85. For saturated live-labelling experiments, Fab fragments were applied at a 1/200 dilution in conditioned medium for 6 min. Cells were fixed using pre-warmed 4% paraformaldehyde (PFA) with sucrose in PBS for 10 min at room temperature. Samples were washed three times with PBS, and residual PFA was quenched using 50 mM NH_4_Cl for 5 min. Coverslips were washed three additional times with PBS and mounted using Fluoromount mounting medium.

#### Epifluorescence microscopy

Fixed samples were imaged using a Leica DM5000B epifluorescence microscope (Leica Microsystems) equipped with a 63 oil immersion objective (NA 1.40). Filter sets included FITC (Leica GFP ET; excitation 450–490 nm, emission 500–550 nm), TRITC (Leica RHOD ET; excitation 534–558 nm, emission 580–620 nm), and Cy5 (Leica Y5; excitation 590–650 nm, emission 665–735 nm).

Excitation was provided by a SOLA SE 6-LCR-SC LED light source (Lumencor; 350–650 nm). Images were acquired using an ORCA-Flash 4.0 V2 camera (Hamamatsu) across six focal planes with a step size of 0.3 *µ*m. All acquisitions were performed using identical parameters.

#### Image analysis

Image analysis was performed using FIJI (NIH) softwares, with custom made macros. Max-projected images were segmented by a pre-trained pixel classification model in Labkit^109^, to detect Xph20 (intrabody against PSD95,^108^) spots in the dendritic spines, and dendritic branch within manually given regions of interest (ROI). GluA2 labelling intensity was measured in these putative PSD spots as well as in a dendritic segment and background with no cellular signal. Then the background-subtracted intensity within the detected putative PSD-spots, the background-subtracted dendritic signal, and their ratio were calculated.

### Time-dependent solution of the extended AMPAR trafficking model

To study plasticity-dependent changes in AMPAR concentration in different compartments (for both GluA1 homomeric and GluA2-containing heteromeric receptors), we used the original time-dependent model in Eqs. (1)-(3) and computed its solution numerically. To this end, we discretized the spatial variable of the PDE (Δ*x* = 1 *µ*m) and transformed the system of PDE into a system of ODEs

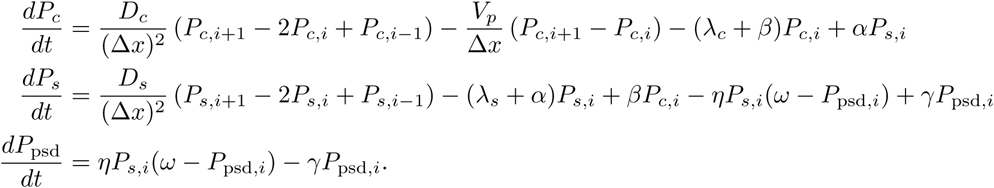

We determined *P_s_*, *P_c_* at all intermediate spatial nodes (*i* = 1*, …, L* 1). For the boundary nodes (*i* = 0*, L*), we used the boundary conditions Eqs. (14)-(15) at *x* = 0 and *x* = *L* and obtained

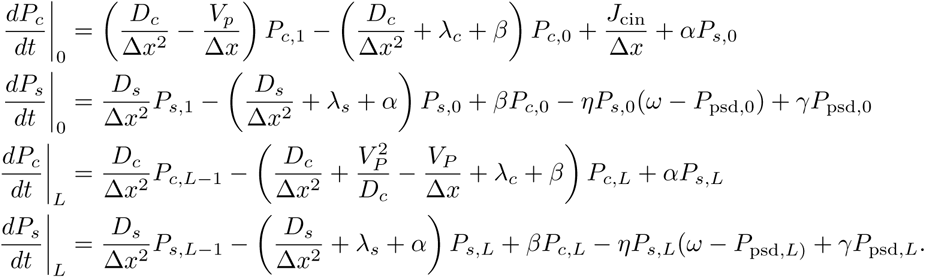

We used the scipy.solve_ivp function to compute *P_c_* and *P_s_* with the explicit Runge-Kutta method of order 5(4).

### Trafficking dynamics and steady state distribution of CNIH-2 mRNA

The production, degradation and transport dynamics of CNIH-2 mRNA molecules can be described by the model proposed in^5^

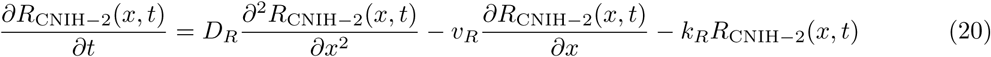

First, we determined the steady state distribution for the Eq. (20). Setting the time-derivative to 0 we obtained

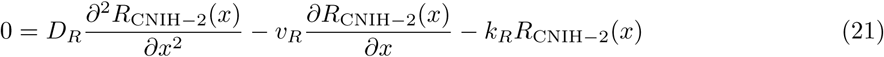

The general solution,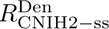, of the Eq. (21) reads

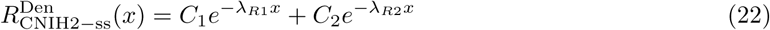

where 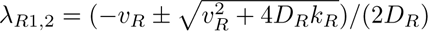. We calculated the constants *C*_1_*, C*_2_ using the following boundary conditions:

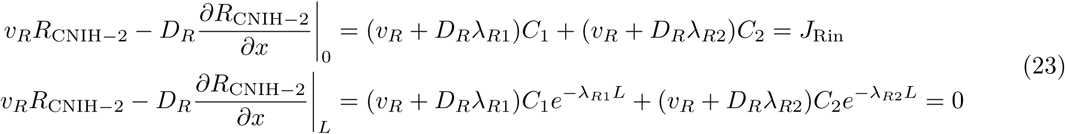

and derived the steady state distribution 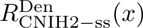. Here *J*_Rin_ is the constant rate of influx of

mRNA from soma into the dendrite. The second boundary condition describes the no-flux boundary condition. We obtained

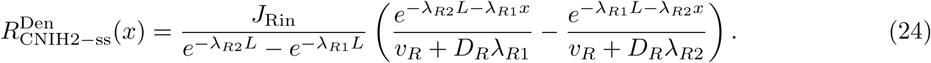

### Trafficking dynamics and steady state distribution of CNIH-2 protein

The CNIH-2 protein model takes into account the CNIH-2 somatic translation rate, local dendritic translation rate, diffusion, and active transport. We used the protein trafficking model introduced in^5^ for CamK2a protein to model the CNIH-2 protein dynamics. We obtained

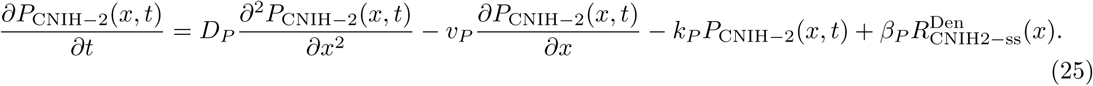

First, we determined the steady state of the Eq. (25) by setting the time-derivative to 0

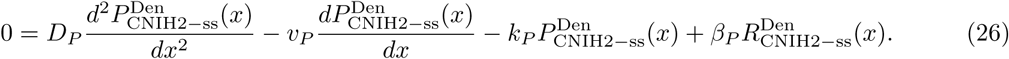

The general solution 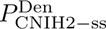 of the Eq. (26) reads

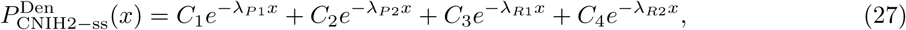

whereby

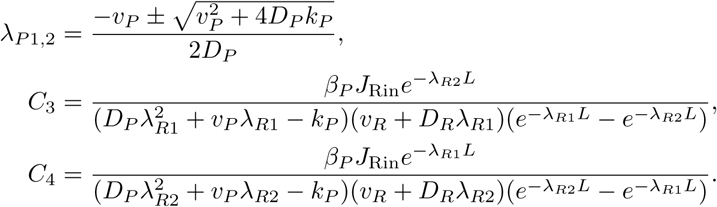

We used the following boundary conditions to obtain the constants *C*_1_ and *C*_2_ in Eq. (27)

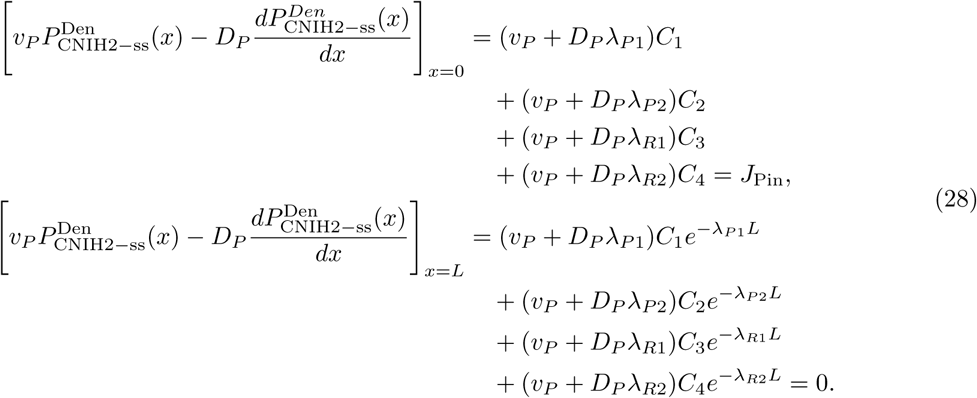

The final steady state CNIH-2 protein distribution reads

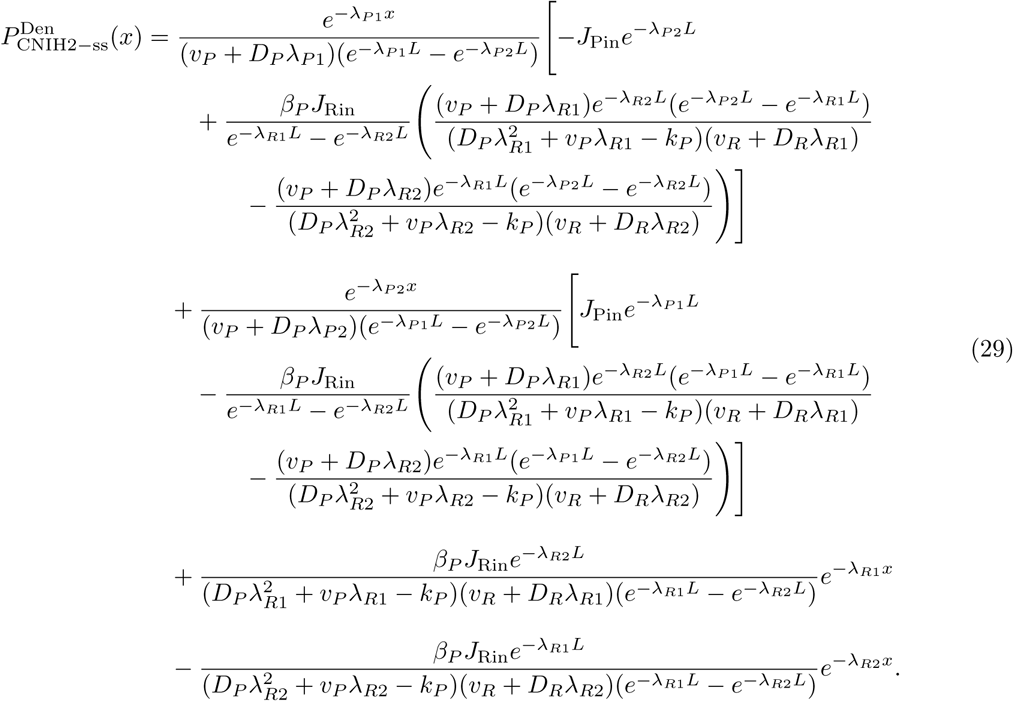

### Time-dependent solution of CNIH-2 subunit trafficking model

How does the concentration of CNIH-2 evolve in our model dendrite over time? To calculate the spatiotemporal distribution of CNIH-2 protein (*P*_CNIH_*_−_*_2_(*x, t*)), we return to the original partial differential equation of CNIH-2 in Eq. 25. In order to simulate the spatio-temporal dynamics of CNIH-2 during basal state and upon cLTP induction, we first discretized the spatial variable *x* of the model and converted PDE to a time-dependent ODE

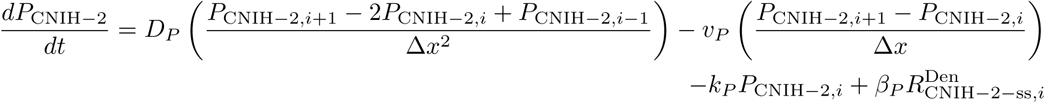

We computed coefficients corresponding to *P_i_*, *P_i−_*_1_ and *P_i_*_+1_ and rewrote the last equation as

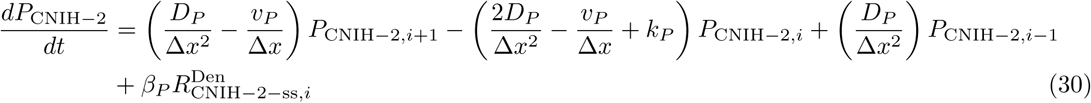

Now we could determine *P* at all intermediate nodes (*i* = 1*, …, L* 1) except the boundary nodes *i* = 0*, L*. For the boundary nodes, we used boundary conditions at *x* = 0 and *x* = *L*

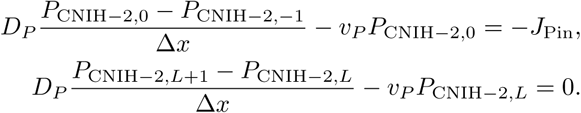

and determined the temporal evolution of the CNIH-2 protein at the boundary points and obtained the differential equations

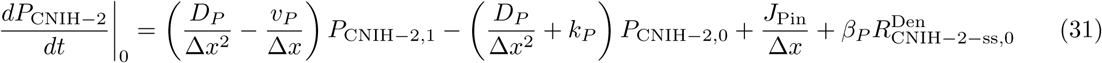

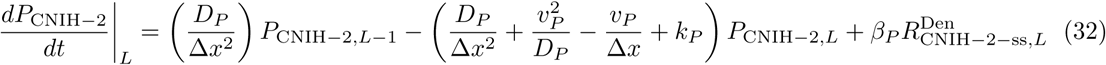

Finally, we used the *solve*_*ivp* function of the scipy Python library to integrate the ODE in Eqs. (30)-(32) and calculated *P*_CNIH_*_−_*_2_ as a function of dendritic location and time.

### CNIH-2 local translation upregulation upon plasticity and exocytosis of GluA2-containing AMPARs

In case of cLTP-driven translation up-regulation (Fig. 5A,C), we multiplied the basal translation rate (*β_P_*) at 10 randomly selected spines throughout the dendrite by the factor of 3000 for 2 hours or 10 minutes following cLTP induction. This is in-line with previous reports of translation up-regulation which peaks at 30 minutes and sustains for 2 hours following cLTP induction^110, 111^. After 2 hours (or 10 minutes) of simulation time, the local translation rate was reset to the basal level.

For the increased somatic influx simulation (in Fig. 5B, E), we increased the in-flux rate *J*_Pin_ by a factor of 3000. After 2 hours of simulation time, we reset the influx rate to a baseline value.

For Fig. S22A&B, the local translation rate (*β_p_*) was upregulated in the complete dendrite by a factor of 500 for either 2 hours or 10 minutes, respectively.

Next, we considered the local concentration of CNIH-2 to be a direct regulator of GluA2-containing AMPAR exocytosis. Hence, for plasticity simulations, we changed the AMPARs exocytosis rate proportionally to the modification in CNIH-2 concentration. More precisely, we normalized the CNIH-2 concentration after plasticity induction by the baseline concentration of CNIH-2 and multiplied it with the basal exocytosis rate

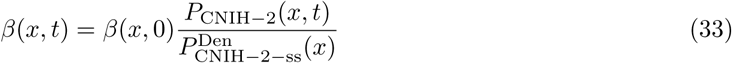

Here, *β*(*x, t*) represents the exocytosis rate as a function of dendritic location and time.

### Estimation of endocytosis rates from published live imaging

Usually, studies investigating receptor internalization, endocytosis rate is measured in terms of events per *µm*^2^ per minute. For the consistency with trafficking parameters in our models, we converted endocytosis rate to represent an event per *µm*^2^ per minute. To this end, we multiplied the reported rate with the number/concentration of receptors per vesicle and the area of the ROIs. For example,^54^ report the endocytosis rate to be (2.1 1.1) 10*^−^*^3^ events per *µm^−^*^2^ per minute for GluA1 and (2.5 0.4) 10*^−^*^3^ events per *µm^−^*^2^ per minute for GluA2 vesicles. These rates were measured in a circular ROI of radius 300 *nm*. The estimated number of AMPARs per vesicle is 50^112, 113^. Using these values we obtain the mean endocytosis rates:

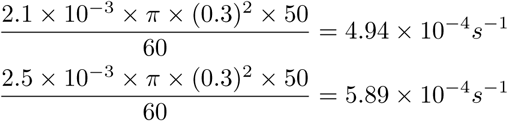

for GluA1 and GluA2 AMPA receptors, respectively.

## Supplementary Figures

**Figure S7:**
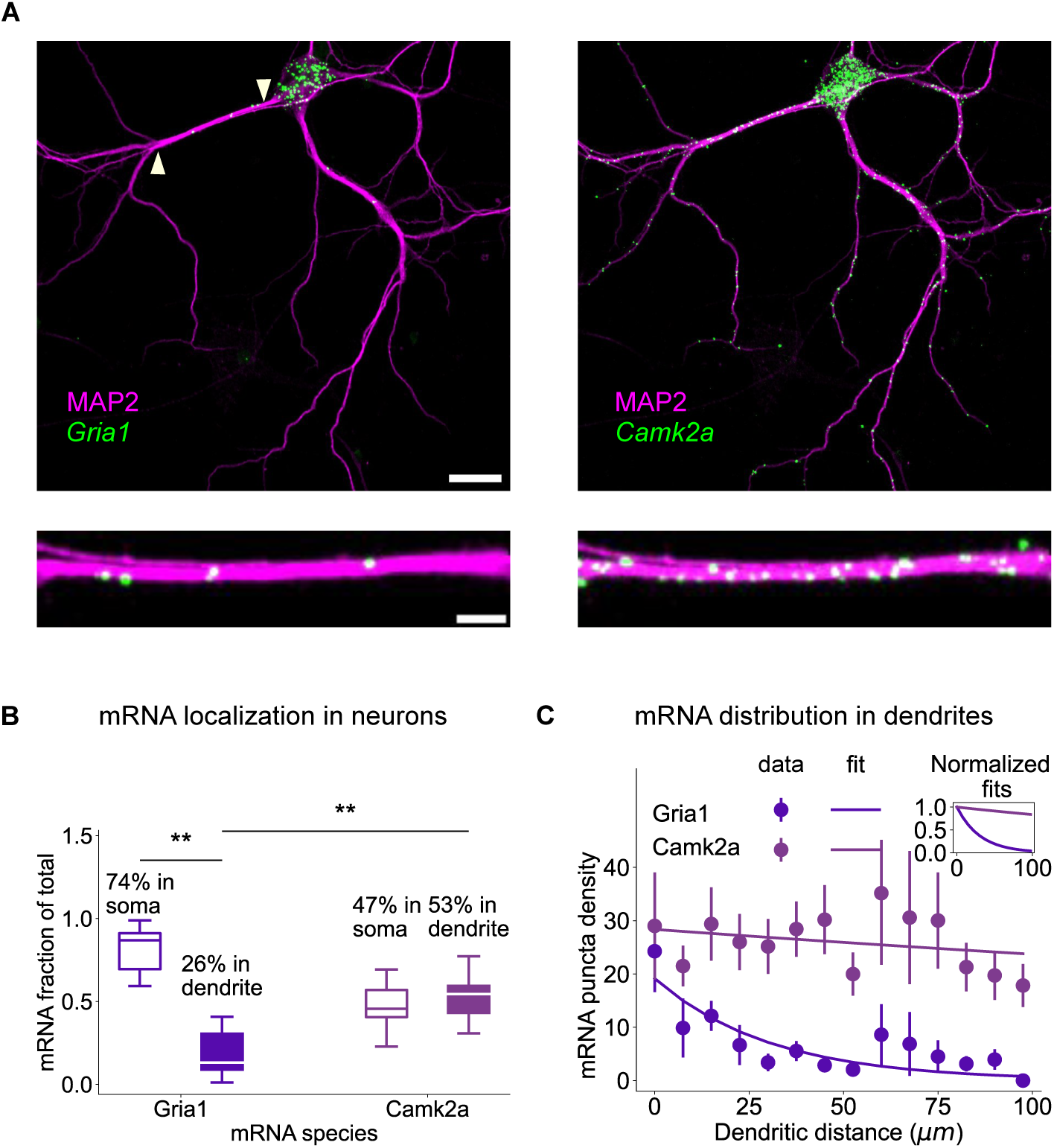
GluA1 mRNA is predominantly localised in the soma. **(A)** *Top*: Cultured rat hippocampal neurons at 18-21 DIV processed for FISH against *Gria1* (left, green) and *CamK2a*mRNA (right, green) and fluorescently immunostained (FI) MAP2 (magenta). Scale bar = 20 *µ*m. *Bottom*: Zooming into a representative dendrite shows a smaller number of *Gria1* mRNA (left) in comparison to *CamK2a* mRNA (right). Scale bar = 5 *µ*m. **(B)** Somatic (hollow) and dendritic (filled) fraction of total mRNA for *Gria1* and *CamK2a*. Departure from equal somatic and dendritic allocation was assessed using two-sided one-sample Wilcoxon signed-rank tests of dendritic fractions against 0.5, with Holm adjustment across the two mRNAs (*Gria1*: *W* = 21, *P*_adj_ = 0.00842; *Camk2a*: *W* = 66, *P*_adj_ = 0.417; *n* = 18 cells per mRNA). Dendritic fractions of *Gria1* and *Camk2a* mRNA were compared using a two-sided Wilcoxon signed-rank test (*W* = 20, *P* = 0.00842; *n* = 18 paired cells). Boxes show the 25th–75th percentiles, with median lines and whiskers extending to the extreme values; Percentages indicate mean somatic and dendritic fractions. (C) Exponential fit of mRNA puncta density distribution for *Gria1* (*n* = 23 dendrites, exponent *Gria1* = −0.03 0.007) compared to the *CamK2a* distribution (*n* = 23 dendrites, exponent *CamK2a* = −0.001 0.001), see Methods for mRNA exponential fit. *Inset*: Normalized fitted *Gria1* mRNA distribution compared to *CamK2a*. **: *p* 0.01. In C, data points are presented as mean SEM. Source data are provided as a Source Data file.

**Figure S8:**
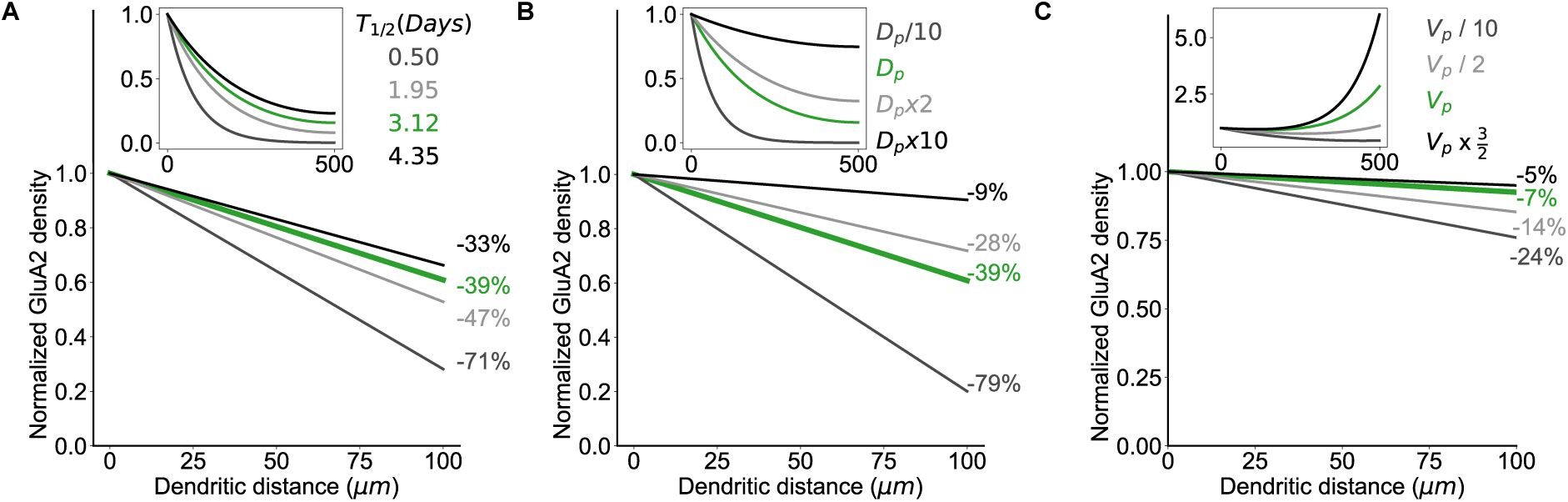
GluA2 distributions for different velocity, diffusion and life-time constants, related to Fig. 1F-H. **(A, B, C)** Parameters are as in Figure 1 in all the panels, we used the value at *x* = 0 to normalize the distributions.

**Figure S9:**
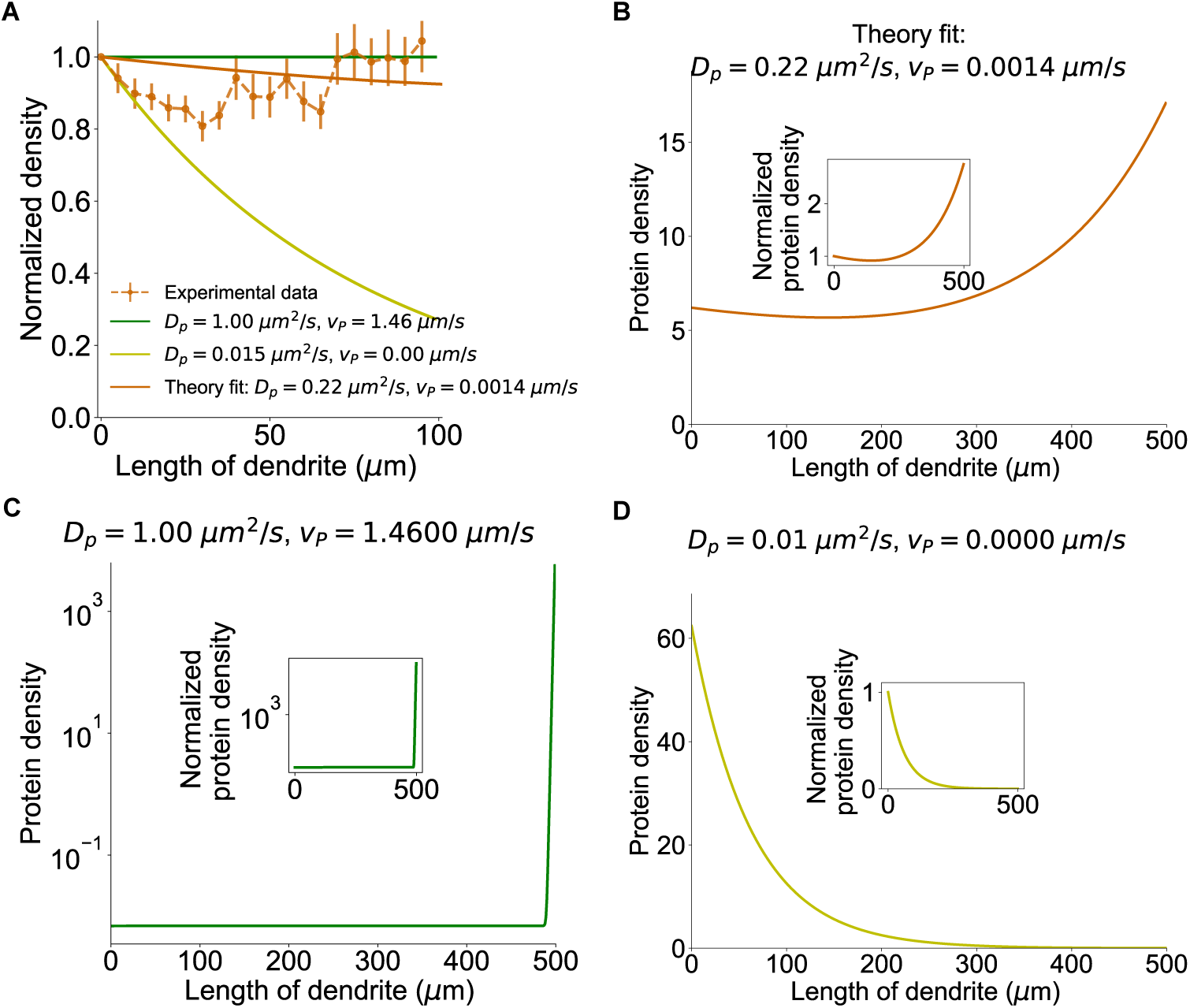
Impact of the active transport velocity on dendritic distribution of AMPARs. **(A)** Binned distribution of GluA2 subunit of AMPAR (data shown as mean SEM), slow and fast transport distribution and theory fit distribution. **(B,C,D)** Steady-state, dendritic distribution of GluA2-containing AMPAR (surface + intracellular) for three different sets of transport parameters. The degradation and the somatic flux parameters were set to *λ_P_* = 2.57 10*^−^*^6^*s^−^*^1^, *J*_Pin_ = 0.01*s^−^*^1^. The insets show the dendritic concentration normalized at the starting point of primary dendrite (*x* = 0). Source data are provided as a Source Data file.

**Figure S10:**
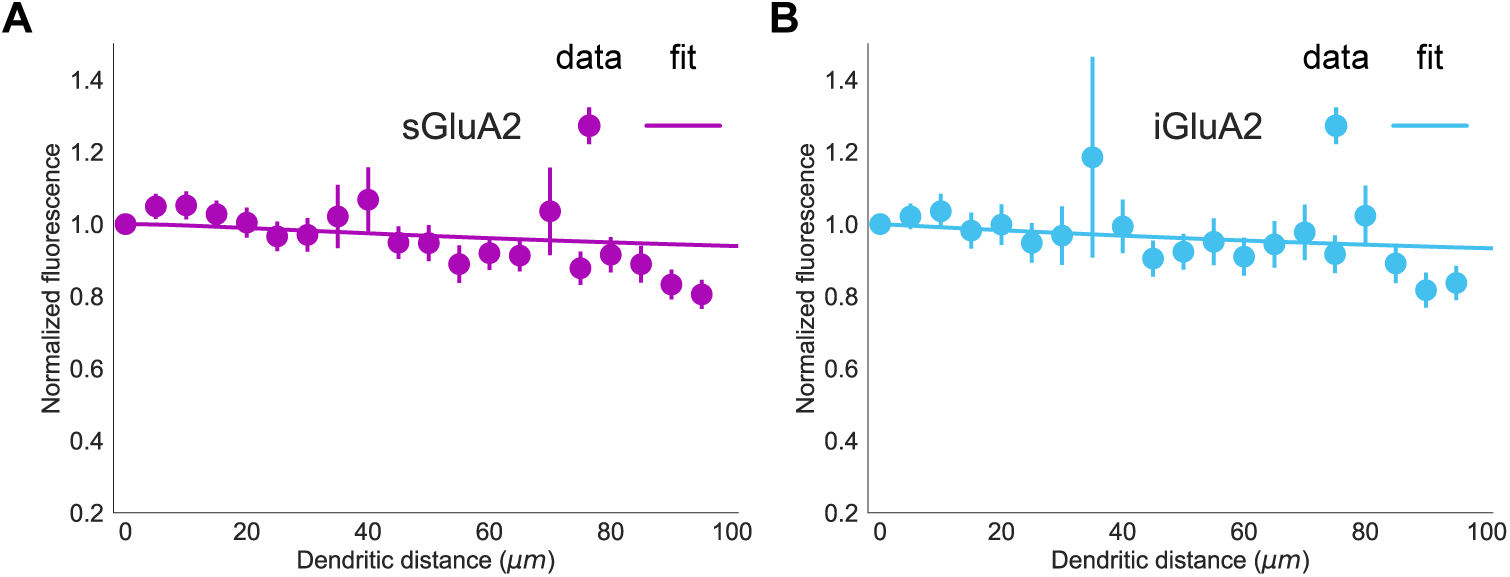
Fitting the global parameters of the full model to surface and intracellular GluA2 fluorescent signaling data. **(A)** Dendritic distribution of soma-normalized GluA2 confined to the surface (sGluA2 data and model fit) and **(B)** intracellular GluA2 (iGluA2 data and model fit, cyan, right, *n* = 176 dendrites). Data points are presented as mean *±* SEM. Source data are provided as a Source Data file.

**Figure S11:**
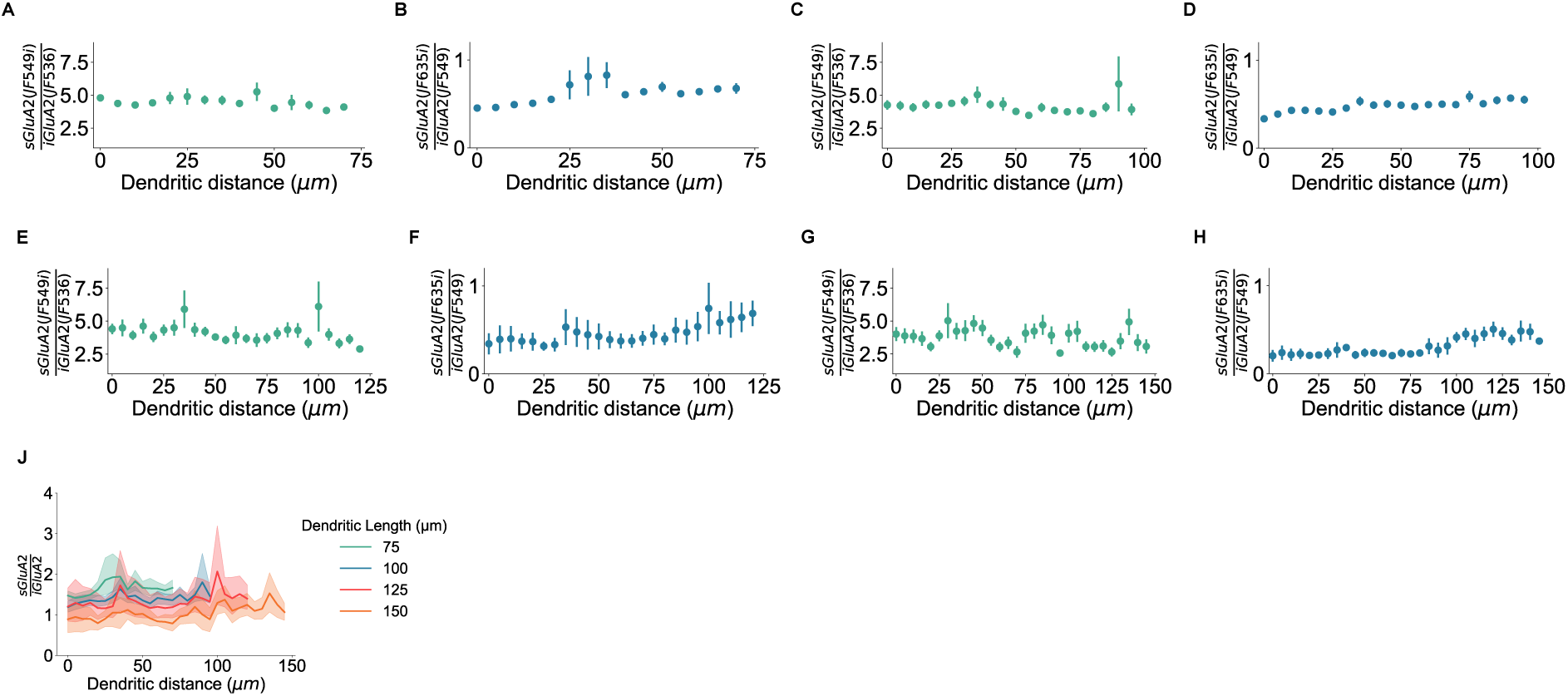
Surface to intracellular ratio is maintained throughout the dendrite. **(A, C, E, G)** Fluorescence intensity ratio between surface (JF549i) and intracellular GluA2 (JF635) along the dendrites of length **(A)** 75 *µm* (*n* = 255 dendrites), **(C)** 100 *µm* (*n* = 124 dendrites), **(E)** 125 *µm* (*n* = 43 dendrites) and **(G)** 150 *µm* (*n* = 15 dendrites). **(B, D, F, H)** Fluorescence intensity ratio between surface (JF635i) and intracellular GluA2 (JF549) along the dendrites of length **(B)** 75 *µm* (*n* = 165 dendrites), **(D)** 100 *µm* (*n* = 52 dendrites), **(F)** 125 *µm* (*n* = 11 dendrites) and **(H)** 150 *µm* (*n* = 5 dendrites). Data points are presented as mean SEM. **(J)** The estimated surface to intracellular ratio as geometric mean of the two data sets (solid lines represent mean curve, shaded region shows 95% confidence interval calculated using bootstrapping). Source data are provided as a Source Data file.

**Figure S12:**
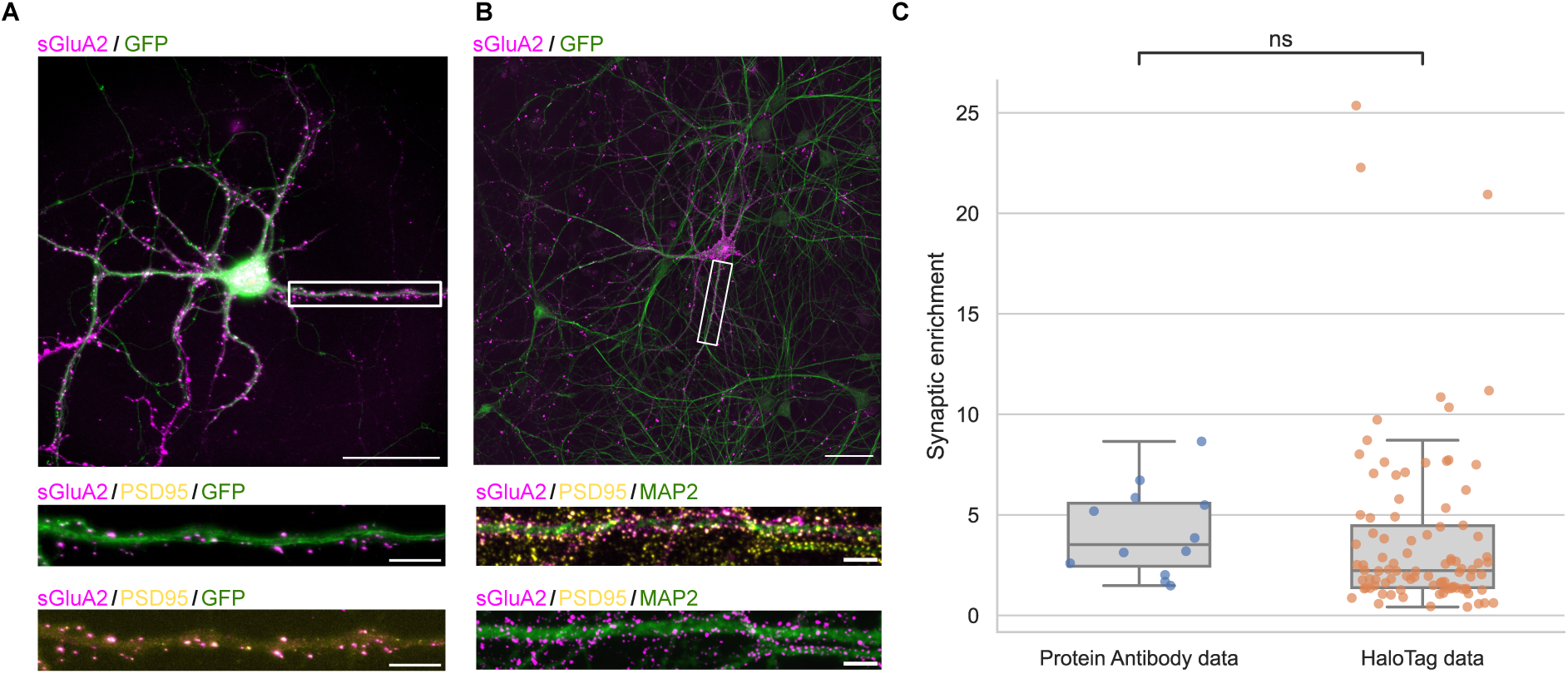
Synaptic enrichment calculated using GluA2-HaloTag is comparable to GluA2 live labeling by protein antibody against GluA2 labeling. **(A)** *Top:* Representative wide-field image of a hippocampal neuron from a primary dissociated culture labeled with *α*-GluA2 (scale bar: 50 *µ*m). *Bottom:* Zoomed in images of representative dendrite labeled for surface GluA2 (sGluA2) in magenta, post synaptic density (PSD95) in yellow and GFP in green (scale bar: 5 *µ*m). **(B)** *Top*: Cultured rat hippocampal neurons at 21 DIV visualized using CRISPR/Cas9 by endogenously tagging GluA2 with HaloTag. Dendrites are labeled using immunostaining to detect MAP2 (green). Surface HaloTag-GluA2 (magenta) was live-cell labeled using the cell-impermeable dyes JF549i. Scale bars 50 *µ*m. *Bottom*: Zooms of representative dendrites labeled for surface HaloTag-GluA2 (magenta), PSD95 (yellow) and MAP2 (green). Scale bars 2 *µ*m. **(C)** Comparison of synaptic enrichment (SE) calculated in neurons from *α*-GluA2 labeling and GluA2-HaloTag labeling (data in Fig. 2). Each individual point corresponds to a cell. Synaptic enrichment of antibody-labelled and HaloTag-labelled surface GluA2 was compared using a two-sided Mann–Whitney U test (U = 738, P = 0.0836; n = 12 and 94 cells, respectively). In C, boxes indicate the interquartile range, and whiskers extend to the most extreme values. Horizontal lines indicate medians. Source data are provided as a Source Data file.

**Figure S13:**
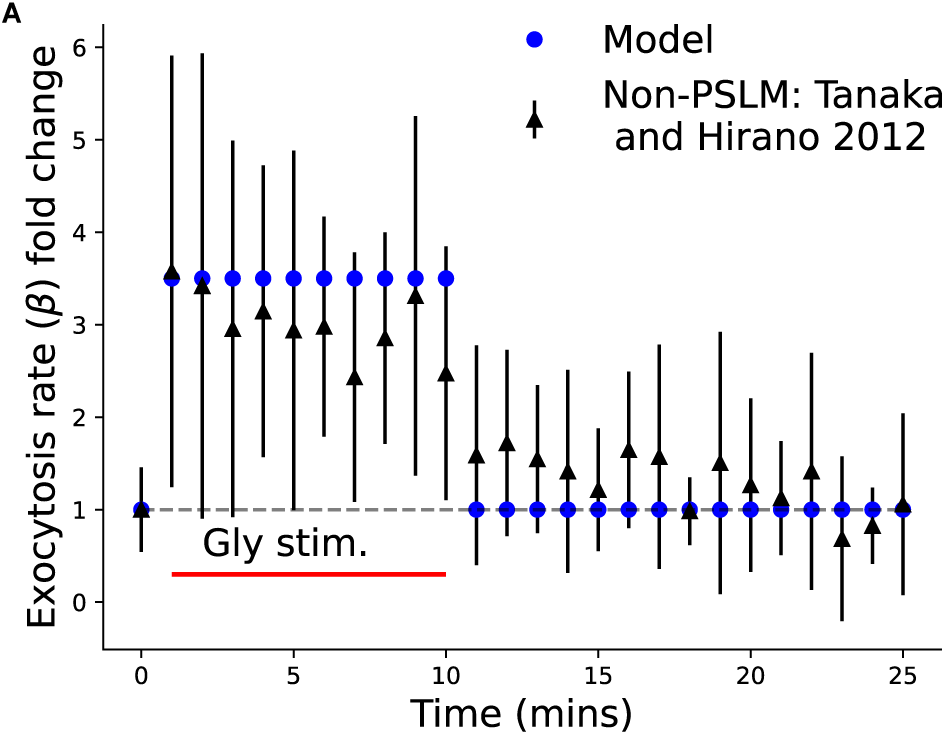
Simulating plasticity response of GluA1-homomeric AMPARs required changes in exocytosis rate *β* and synaptic incorporation rate *η*. **(A)** We used a step function where the exocytosis rate (*β*) was increased 3.5 fold to the basal rate. This matches well with the change in exocytosis frequency reported in (^66^ Fig. S3C). Data points are presented as mean SEM, blue circles represent the values used in simulations. Source data are provided as a Source Data file.

**Figure S14:**
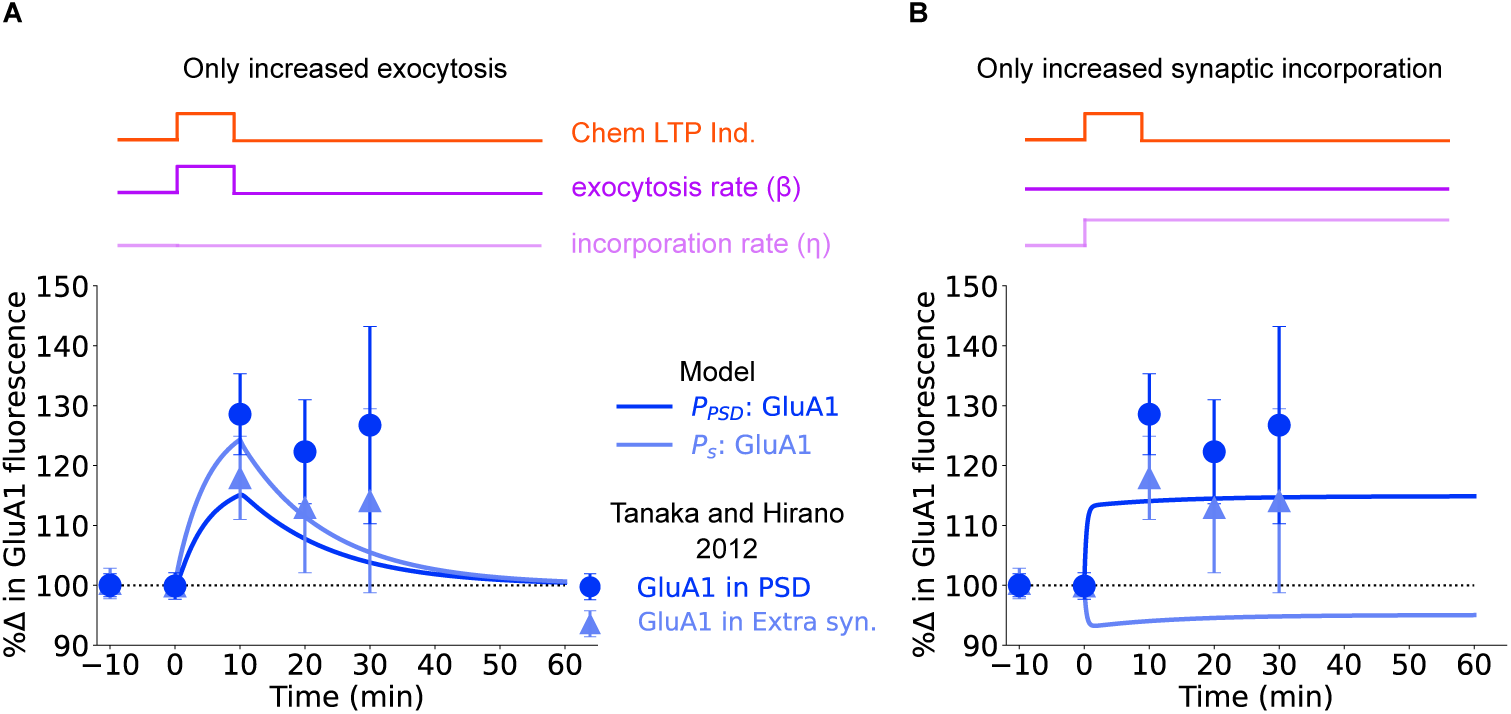
GluA1-homomeric AMPAR requires fast increase in exocytosis and enhanced diffusional trapping for LTP dependent changes (related to Fig. 3A). In both subpanels, a 10 min chemical LTP stimulation was applied which led to different changes in trafficking parameters. **(A)** The exocytosis rate (*β*) was stepwise increased 3.5 fold for the duration of stimulus followed by a stepwise decrease back to the basal level. The synaptic incorporation (*η*) rate was left unaltered. **(B)** The exocytosis rate (*β*) was kept at basal level; however the synaptic incorporation rate (*η*) was increased by 30%. None of the simulations could replicate the experimental observations. Data points are presented as mean SEM, solid lines represent mean response curves from the model. Source data are provided as a Source Data file.

**Figure S15:**
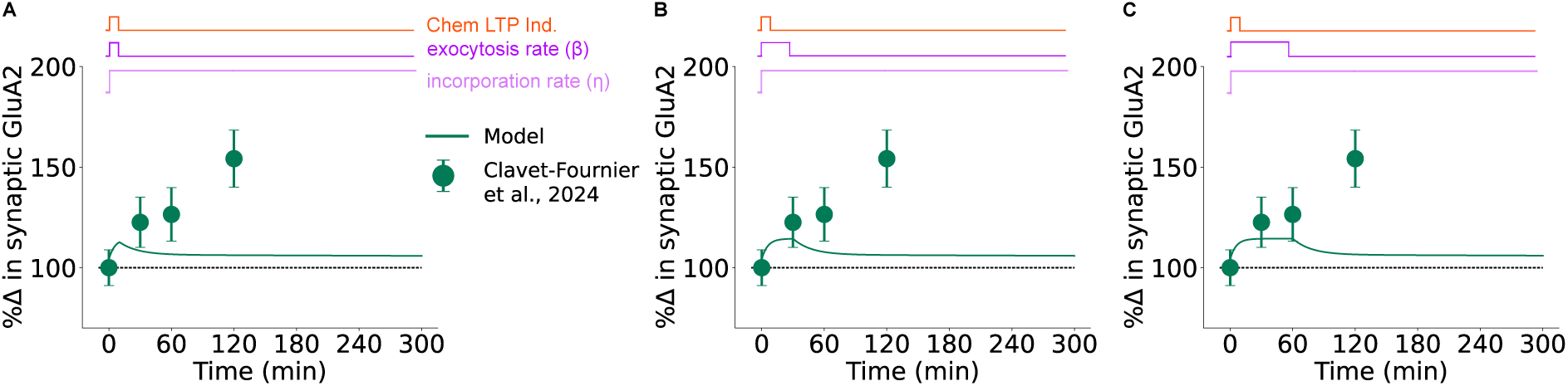
Constant increase in GluA2-AMPARs exocytosis cannot explain their temporal profile upon cLTP induction. We simulated the GluA2-containing AMPARs model with a constant increase in exocytosis rate by 3.5 folds for **(A)** 10 minutes, **(B)** 30 minutes, and **(C)** 1 hour. In addition, the synaptic incorporation rate *η* was increased by 1.3 folds for the entire duration of simulation. None of the simulations matched the experimental profile of synaptic GluA2 increase from^62^. Data points are presented as mean SEM, solid lines represent mean response curves from the model. Source data are provided as a Source Data file.

**Figure S16:**
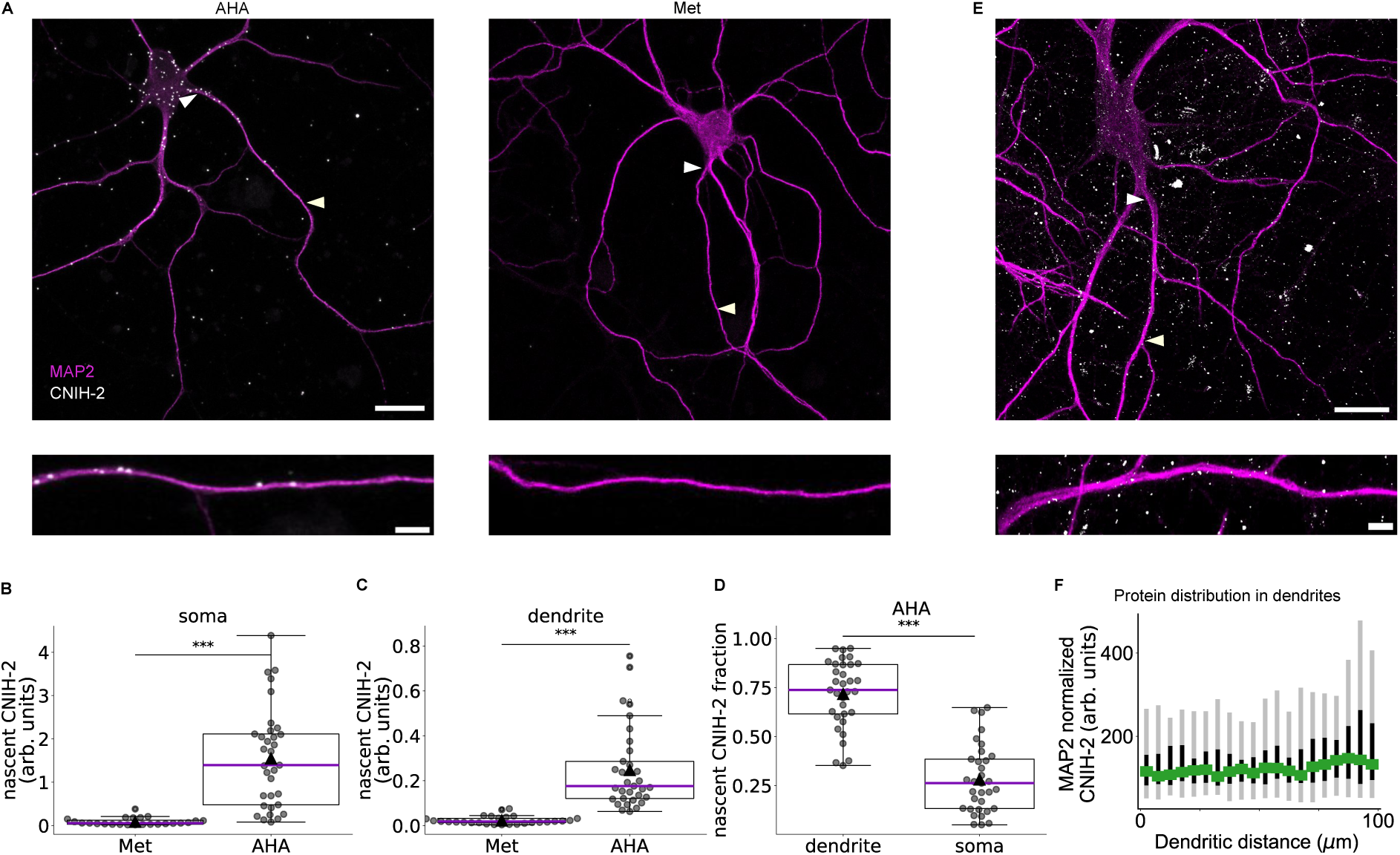
Auxiliary subunit CNIH-2 is translated locally in the dendrite. **(A)** *Top:* Representative images of cultured hippocampal neurons showing the distribution of the newly synthesized AMPAR auxiliary subunit CNIH-2 after 1 hr of metabolic labeling with AHA or, in control conditions, with methionine (Met). Scale bar = 20 *µ*m. *Bottom:* Representative dendrites showing that CNIH-2 nascent protein is abundant in a dendrite after 1h of metabolic labeling. Scale bar = 10 *µ*m. **(B, C)** Box and strip plot showing density of new CNIH-2 **(B)** in somata and **(C)** in dendrites. **(D)** Fraction of total nascent protein in AHA treatment found in dendrite vs somatic compartment. Boxes indicate the interquartile range, and whiskers extend to the most extreme values. Horizontal lines indicate medians; black triangles indicate means and individual points represent cell-level measurements. Nascent CNIH-2 densities in Met and AHA conditions were compared using two-sided Mann–Whitney *U* tests: soma (B), *U* = 23, *P* = 1.74 10*^−^*^9^ (*n* = 24 Met and 33 AHA cells); dendrites (C), *U* = 4, *P* = 3.56 10*^−^*^11^ (*n* = 28 Met and 33 AHA cells). Somatic and dendritic fractions in AHA-labelled cells (D) were compared using a two-sided Wilcoxon signed-rank test (*W* = 31, *P* = 8.27 10*^−^*^6^; *n* = 33 paired cells). **(E)** *Top*: Cultured rat hippocampal neurons at DIV 18-21 processed for antibody labeling of CNIH-2 protein (gray) and FI MAP2 (magenta). *Bottom*: Zoom into a representative dendrite shows a CNIH-2 distribution along the first 100 *µ*m. Scale bar = 20 *µ*m. **(F)** MAP2 normalized CNIH-2 intensity (5 *µ*m bins, black vertical lines show the 25th–75th percentiles with median represented as green squares and whiskers extending to the extreme as gray bars, *n* = 35 dendrites). ***: *p ≤* 0.001. Source data are provided as a Source Data file.

**Figure S17:**
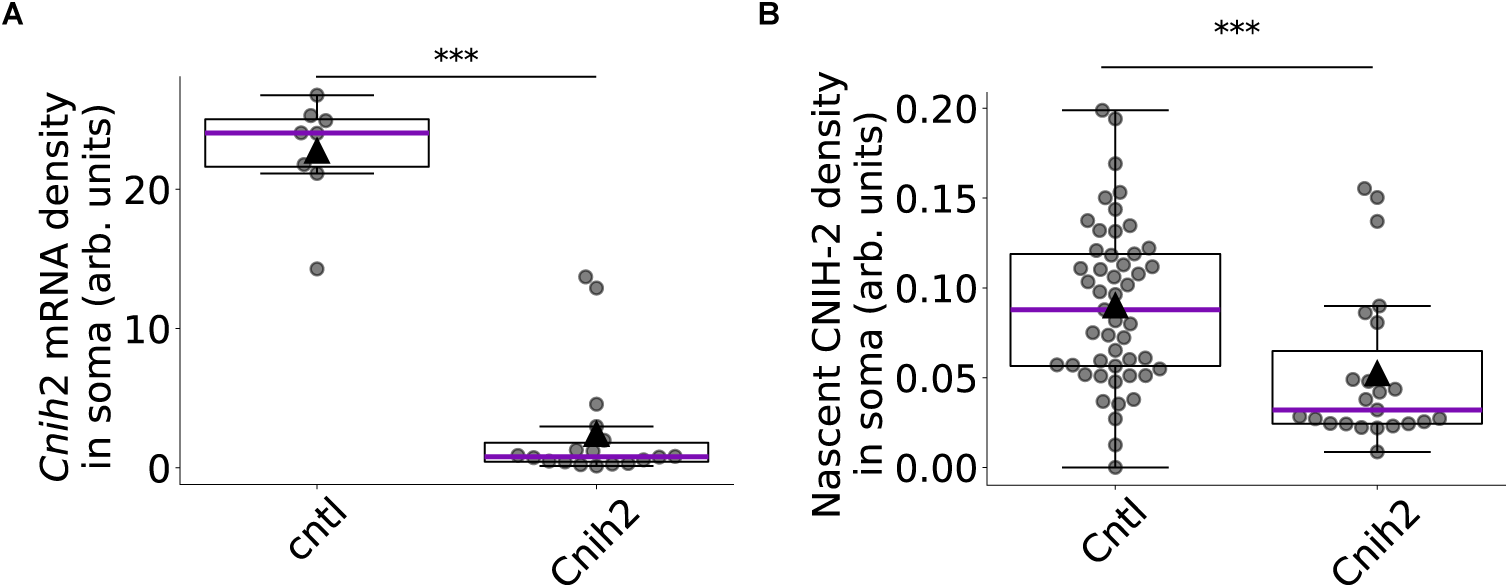
Validation of CNIH-2 knockdown using an shRNA strategy. **(A)** Bar graphs and strip plot showing the density of *Cnih2* mRNA in soma of neurons expressing a control shRNA or an shRNA against *Cnih2*. **(B)** Bar graphs and strip plot showing the density of nascent CNIH-2 proteins in soma of neurons expressing a control shRNA or an shRNA against *Cnih2*. A two-sided Mann–Whitney *U* test was used to assess statistical significance. For mRNA, *U* = 144, *P* = 1.28 10*^−^*^6^ (*n* = 8 control and 18 knockdown cells); for protein, *U* = 863, *P* = 3.05 10*^−^*^4^ (*n* = 49 control and 23 knockdown cells). Boxes indicate the interquartile range, and whiskers extend to the most extreme values. Horizontal lines indicate medians; black triangles indicate means and individual points represent cell-level measurements. ***: *p ≤* 0.001. Source data are provided as a Source Data file.

**Figure S18:**
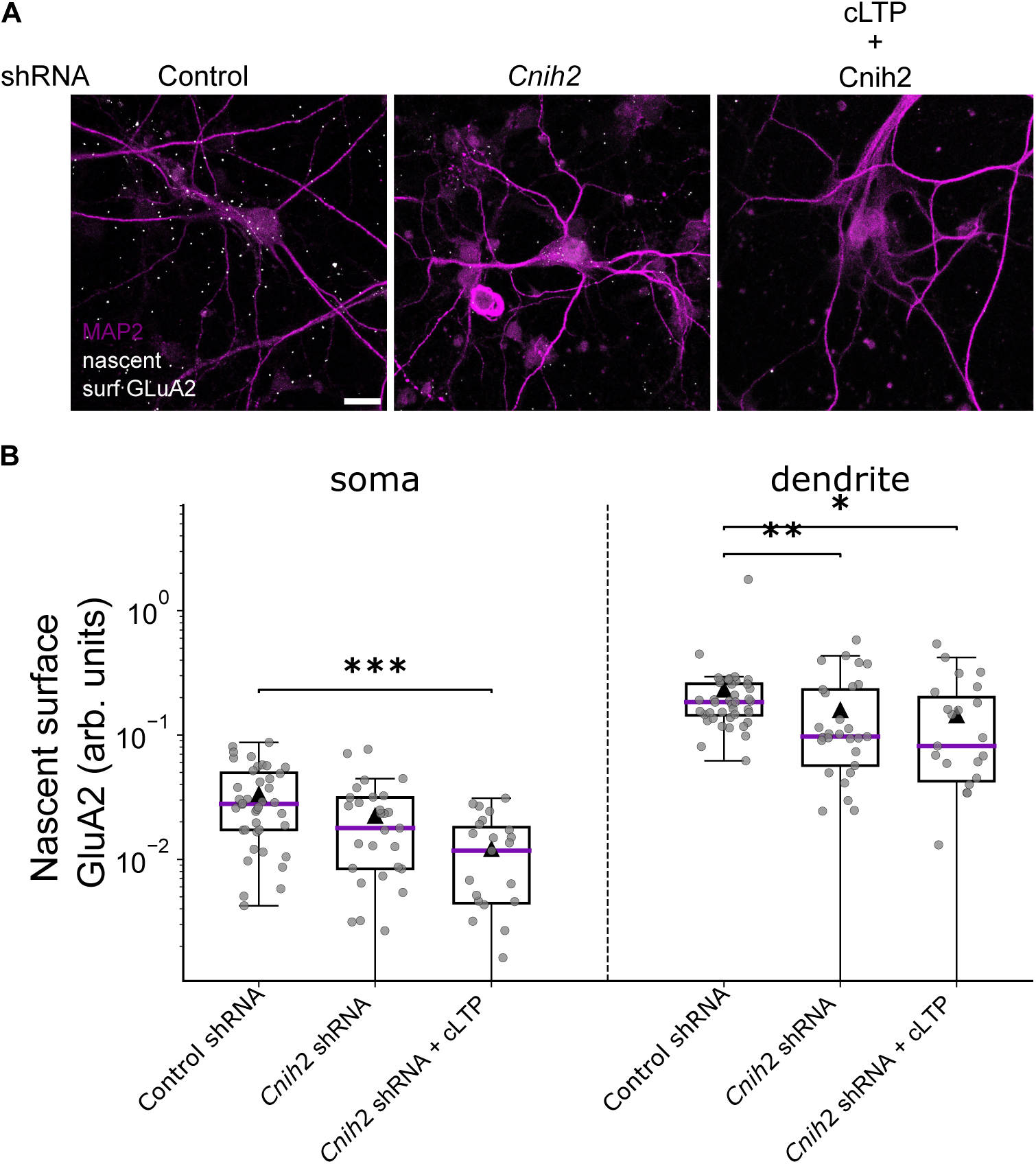
CNIH-2 local synthesis regulates surface insertion of GluA2-containing AMPARs upon chemical LTP induction. **(A)** Cultured hippocampal neurons showing newly synthesized surface GluA2 subunits in neurons expressing a control shRNA (first image) or an shRNA against CNIH-2 under basal condition (second image). Some neurons also underwent cLTP induction (third image). Without CNIH-2 proteins, newly synthesized GluA2-containing AMPARs are absent at the somatic and dendritic surface under both basal and cLTP conditions. Scale bar 20 *µ*m. **(B)** Box plots show the abundance of newly synthesized GluA2 containing receptors at the cell surface in soma and dendrite. Nascent surface GluA2 densities were compared using two-sided Mann–Whitney U tests, with Holm adjustment across all six comparisons. Comparisons against Control shRNA were: soma, *Cnih2* shRNA (*U* = 763, *P*_adj_ = 0.0795) and *Cnih2* shRNA + cLTP (*U* = 753, *P*_adj_ = 1.78 10*^−^*^4^); dendrites, *Cnih2* shRNA (*U* = 825, *P*_adj_ = 0.0148) and *Cnih2* shRNA + cLTP (*U* = 644, *P*_adj_ = 0.0352). Comparisons between *Cnih2* shRNA and *Cnih2* shRNA + cLTP were: soma (*U* = 449, *P*_adj_ = 0.0795) and dendrites (*U* = 368, *P*_adj_ = 0.5310). *n* = 40 Control shRNA, *n* = 29 *Cnih2* shRNA, and *n* = 23 *Cnih2* shRNA + cLTP cells per compartment. Boxes indicate the interquartile range, and whiskers extend to the most extreme values. Horizontal lines indicate medians; black triangles indicate means and individual points represent cell-level measurements. *: p *≤* 0.05, **: p *≤* 0.01, ***: p *≤* 0.001. Source data are provided as a Source Data file.

**Figure S19:**
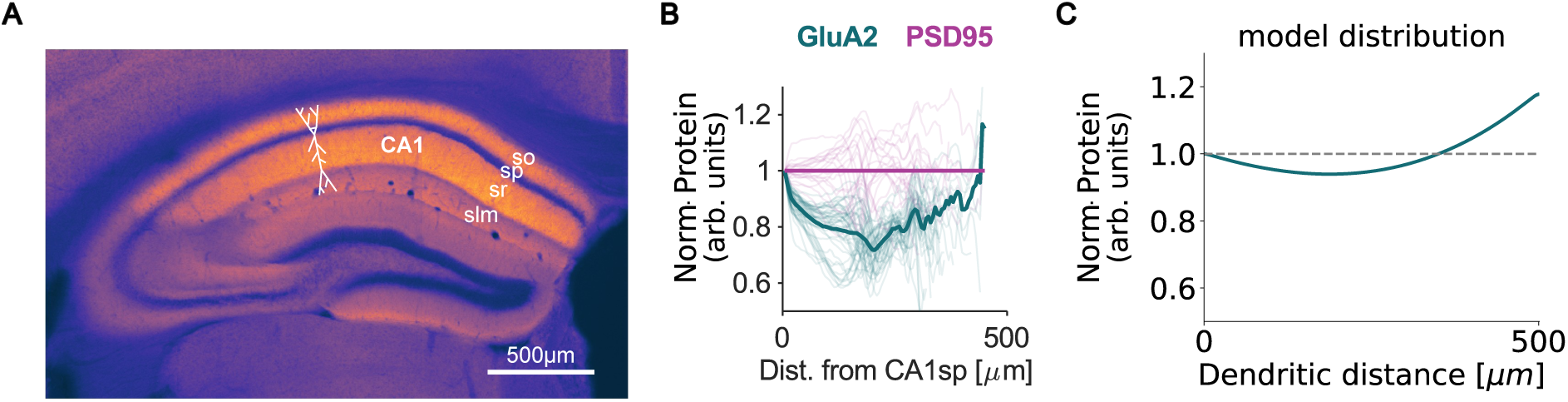
Dendritic distribution of endogenous GluA2 *in-vivo* shows increase toward distal site. **(A)** Example image of GluA2-HaloTag in a coronal section. Scale bar 500 *µ*m. **(B)** Normalized distribution of GluA2 protein shows an increase toward the distal tip (PSD95 is used to normalize the density of GluA2 protein). Dark solid lines represent mean distributions while light thin lines show single cell distributions. **(C)** Total GluA2 distribution from the extended model suggests an 15% increase toward the distal end. GluA2 and PSD95 HaloTag-data in this figure are from^63^.

**Figure S20:**
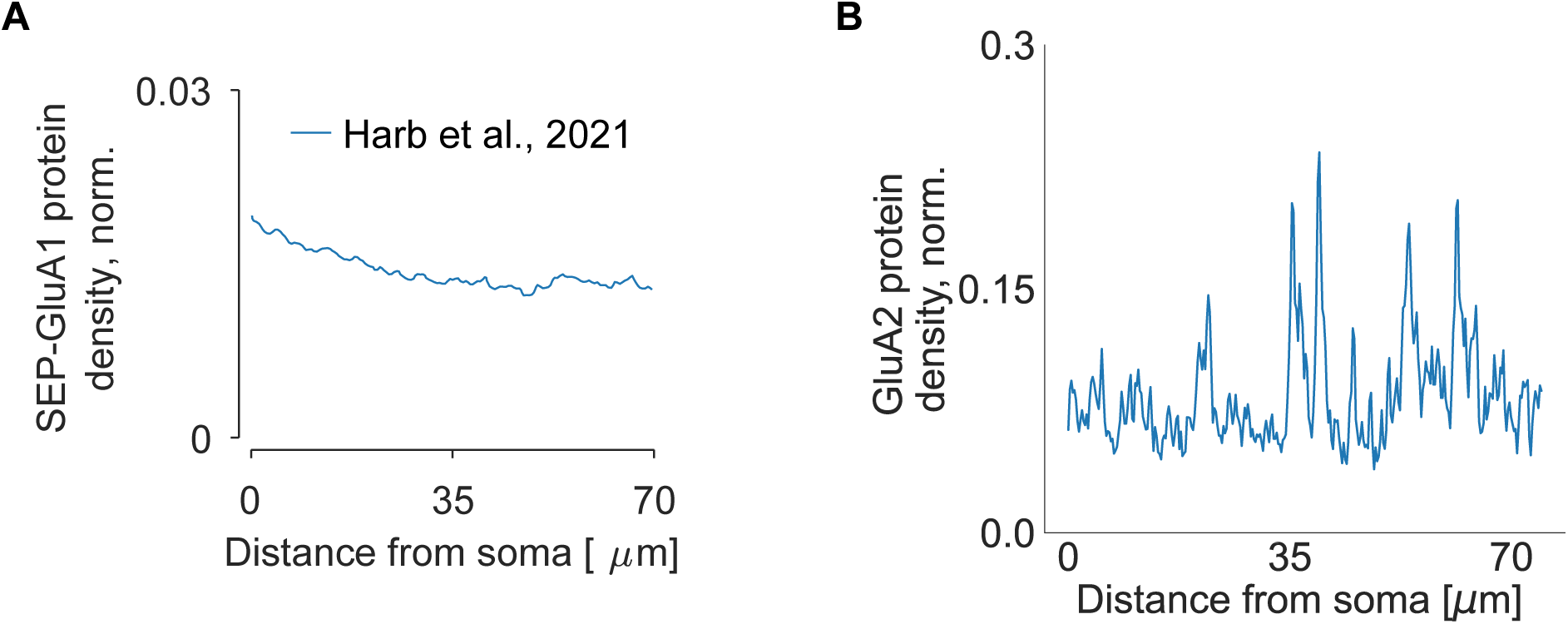
Dendritic distribution of endogenous GluA1-homomeric and GluA2-containing AMPARs in proximal dendrites. **(A)** Distribution of GluA1-homomeric AMPARs density (normalized to soma) in first 70 *µ*m (data from^114^). **(B)** Same for GluA2-containing AMPAR.

**Figure S21:**
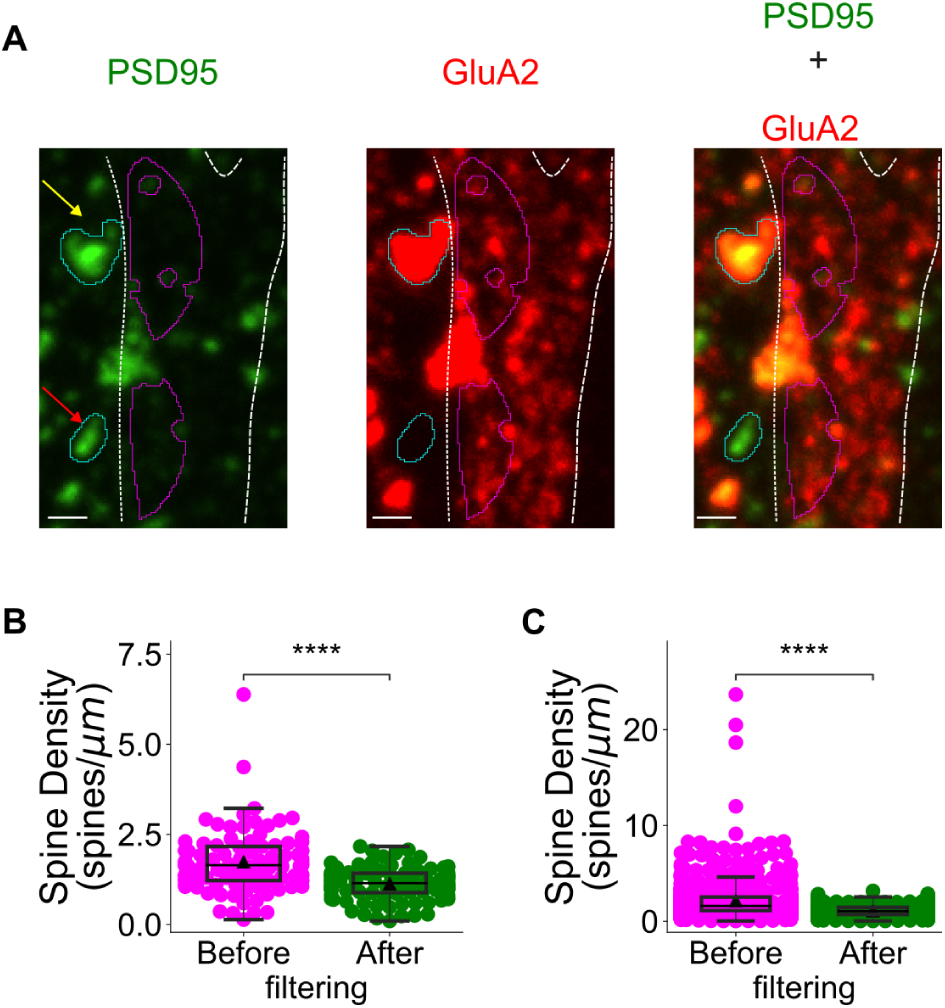
Filtering of empty/non-specific spines. **(A)** Representative image of cultured rat hippocampal neurons at 21 DIV visualized using CRISPR/Cas9 by endogenously tagging GluA2 with HaloTag. Dendrites are labeled using immunostaining to detect MAP2. *Left*: PSD95 (green), *middle*: Surface HaloTag-GluA2 (red), *right*: overlap. Yellow arrow points to a spine that has PSD95 and sGluA2 colocalized, red arrow points to the PSD95 void of sGluA2. **(B)** Box-strip plot of spine density per cell before and after filtering out the spine void of sGluA2 signal. (W = 2, P = 5.80 10*^−^*^10^; *n* = 94 paired cells). **(C)** The same as **(B)** but for individual ROIs. (W = 0, P = 2.54 10*^−^*^39^; *n* = 560 matched dendrites). Before and after filtering densities were compared using a two-sided Wilcoxon signed-rank test. Boxes indicate the interquartile range, and whiskers extend to the most extreme values. Horizontal lines indicate medians; black triangles indicate means and individual points represent cell-level (in B) and ROI-level (in C) measurements. Source data are provided as a Source Data file.

**Figure S22:**
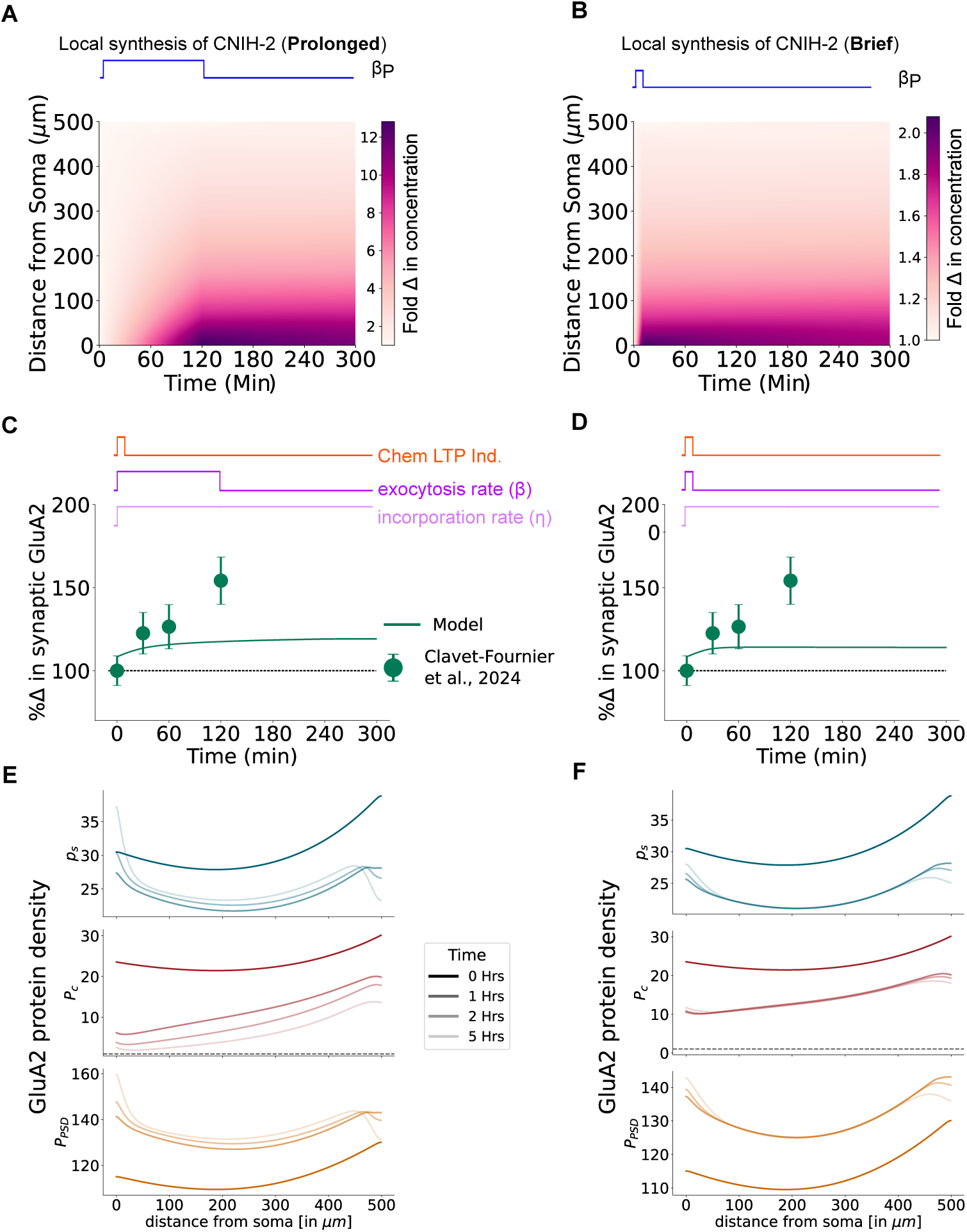
cLTP at all synapses in the model leads to competition for resources. **(A, B)** The spatiotemporal dynamics of CNIH-2 density upon chemical LTP induced increase in local CNIH-2 synthesis rate throughout the model dendrite that lasted for a **(A)** prolonged 2 hours duration or **(B)** brief 10 minutes duration. **(C, D)** The **(C)** prolonged and **(D)** brief time and space-dependent increase in CNIH-2 concentration was used to model the increase in the GluA2 exocytosis rate. The resulting GluA2-containing AMPAR distribution failed to match the temporal response reported in^62^. In addition, we stepwise increased the synaptic incorporation rate *η* by 70% (*η^cLTP^ /η* = 1.7). **(E, F)** Spatial distribution of GluA2 in surface (*P_s_*), intracellular (*P_c_*) and synaptic (*P_psd_*) compartment at t = 0, 1 hour, 2 hours and 5 hours in case of **(E)** prolonged and **(F)** brief CNIH-2 local synthesis upregulation. In both cases the synaptic GluA2 concentration saturated to a much lower value at earlier time points. In case of prolonged upregulation, lower synaptic GluA2 concentration was due to depletion of intracellular GluA2 concentration (in **E**, middle plot). At the same time, the brief upregulation did not cause enough exocytosis to meet the LTP induction demands (in **F**). Data points in (C) and (D) are represented as mean SEM, solid line represents mean response curve from the model. Source data are provided as a Source Data file.

